# Biobank-scale inference of ancestral recombination graphs enables genealogy-based mixed model association of complex traits

**DOI:** 10.1101/2021.11.03.466843

**Authors:** Brian C. Zhang, Arjun Biddanda, Pier Francesco Palamara

## Abstract

Accurate inference of gene genealogies from genetic data has the potential to facilitate a wide range of analyses. We introduce a method for accurately inferring biobank-scale genome-wide genealogies from sequencing or genotyping array data, as well as strategies to utilize genealogies within linear mixed models to perform association and other complex trait analyses. We use these new methods to build genome-wide genealogies using genotyping data for 337,464 UK Biobank individuals and to detect associations in 7 complex traits. Genealogy-based association detects more rare and ultra-rare signals (*N* = 133, frequency range 0.0004% - 0.1%) than genotype imputation from ∼65,000 sequenced haplotypes (*N* = 65). In a subset of 138,039 exome sequencing samples, these associations strongly tag (average *r* = 0.72) underlying sequencing variants, which are enriched for missense (2.3×) and loss-of-function (4.5×) variation. Inferred genealogies also capture additional association signals in higher frequency variants. These results demonstrate that large-scale inference of gene genealogies may be leveraged in the analysis of complex traits, complementing approaches that require the availability of large, population-specific sequencing panels.

## Introduction

Modeling of genealogical relationships between individuals plays a key role in a wide range of analyses, including the study of natural selection [1] and demographic history [2], genotype phasing [3], and imputation [4]. Data-driven probabilistic inference of genealogical relationships, however, is computationally difficult, and available methods for explicit inference of partial or complete genealogies rely on strategies that trade model simplification for computational scalability. These strategies have included probabilistic inference based on approximate models [5, 6, 7, 8, 9, 10, 11], scalable heuristic approaches [12, 13, 14, 15, 16], or combinations of probabilistic and heuristic components [17, 18]. Recent algorithms have enabled efficient estimation of coalescence times from ascertained genotype data [11], rapid genealogical approximations for hundreds of thousands of samples [15], and improved scalability of probabilistic genealogical inference [17]. However, available methods do not simultaneously offer all of these features, so that scalable and accurate genealogical inference in modern biobanks remains challenging. In addition, these data sets contain extensive phenotypic and environmental information, but applications of inferred genealogies have been primarily focused on evolutionary analyses. Early work suggested that genealogical data may be used to improve association and fine-mapping [19, 13], but the connections between genome-wide genealogical inference and recent methodological advances for complex trait analysis [20, 21, 22] remain under-explored.

To address these limitations, we introduce a new algorithm, called ARG-Needle, that accurately infers the ancestral recombination graph [23] (ARG) from large collections of genotyping or sequencing samples. We demonstrate that the ARG of a collection of samples may be coupled with a linear mixed model (LMM) framework to increase association power, detect association to unobserved genomic variants, and perform additional analyses such as the inference of narrow sense heritability or polygenic prediction. Using ARG-Needle, we infer the ARG for 337,464 genotyped British samples from the UK Biobank and perform a genealogy-wide scan for association to 7 complex traits. We show that although it is inferred using only array data, the ARG detects more independent associations to rare and ultra-rare variants (minor allele frequency, MAF < 0.1%) than imputation based on a reference panel of ∼65,000 sequenced haplotypes of matched predominantly European ancestry. We use 138,039 exome sequencing samples to confirm that these signals implicate sets of individuals carrying unobserved sequencing variants, which are strongly enriched for missense and loss-of-function variation. These variants also substantially overlap with likely causal associations detected using imputation from a large within-cohort exome sequencing panel. Using the ARG, we detect novel associations to variants as rare as MAF ≈ 4 × 10^*-*6^ and independent higher frequency variation that is not captured using imputation.

## Results

### Overview of the ARG-Needle algorithm

The ARG is a graph in which nodes represent the genomes of individuals or their ancestors and edges represent genealogical connections between them (see Supplementary Note 1). ARG-Needle infers the ARG for large genotyping array or sequencing data sets by iteratively “threading” [9] one haploid sample at a time to an existing ARG, as depicted in Fig. 1. Given an existing ARG, which we initialize to contain a single sample, we randomly select the next sample to be added (or threaded) to the ARG. We then compute a *threading instruction*, which at each genomic position provides the index of a sample in the ARG that is estimated to be most closely related to the target sample, as well as their time to most recent common ancestor (TMRCA). We use this threading instruction to add the target sample to the ARG and continue iterating these steps until all samples are in the ARG. The ARG-Needle threading algorithm is guaranteed to recover the true ARG if correct threading instructions are used (see Supplementary Note 1).

**Figure 1:**
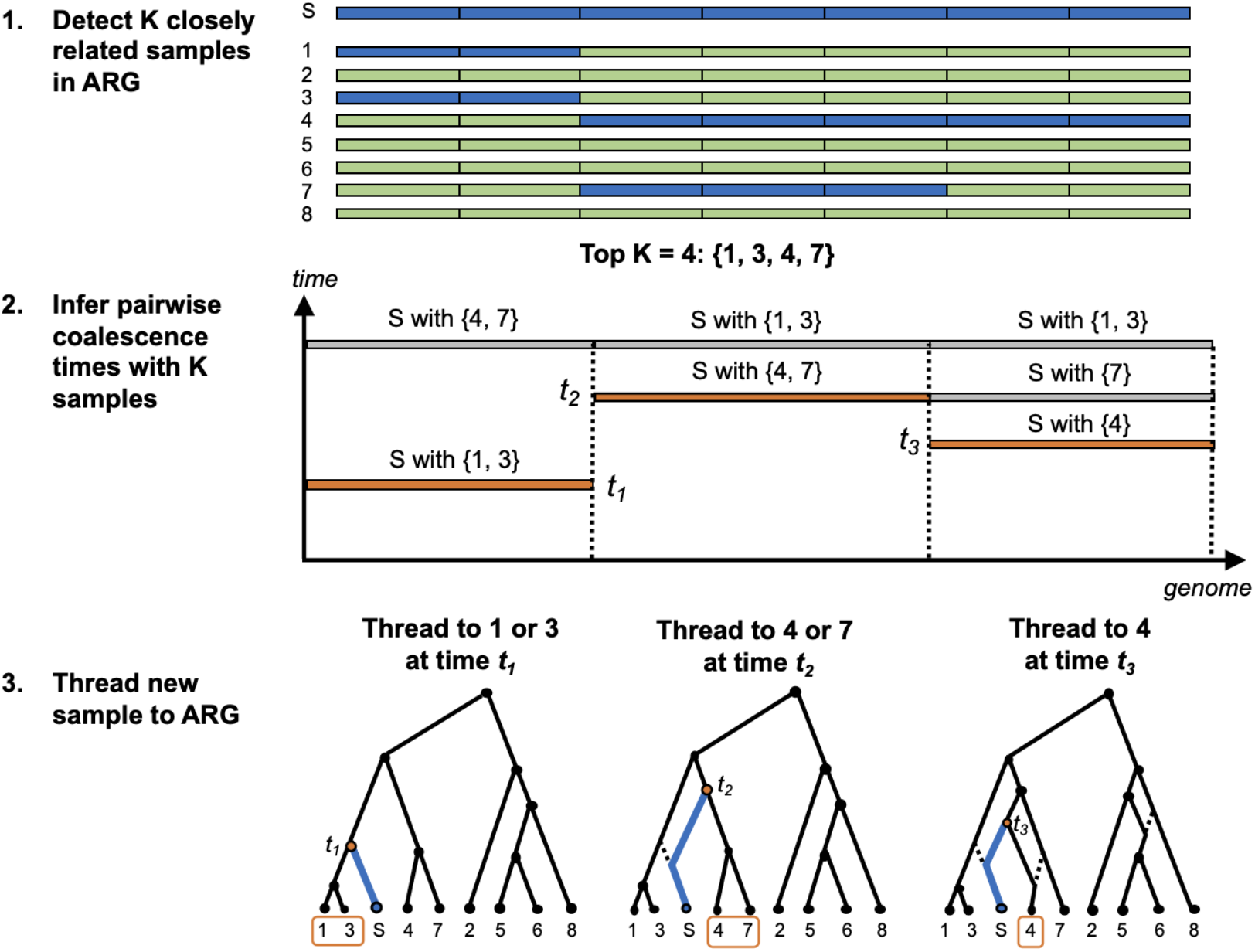
Overview of the ARG-Needle algorithm. ARG-Needle adds one haploid sample at a time to an existing ARG, each time performing three steps: 1. shortlisting a subset of most related samples already in the ARG through genotype hashing, 2. obtaining pairwise coalescence time estimates with these samples using ASMC [11], and 3. using the ASMC output to “thread” [9] the new sample to the ARG. We depict an example of adding sample S to an ARG, focusing on one genomic region. Step 1 divides the genome into “words” and checks for identical matches with sample S. Based on these matches (shown in blue), samples 1, 3, 4, and 7 are output as the *K* = 4 candidate most related samples already in the ARG. Step 2 computes pairwise coalescence time estimates between sample S and each of the samples 1, 3, 4, and 7. The minimum time for each position is highlighted. Step 3 uses these minimum times and samples to define a “threading instruction” that is performed to add sample S to the ARG. Threading connects the new sample to the ancestral lineage of each chosen sample at the chosen time. Dotted lines indicate past ARG edges that are inactive due to recombination. When all samples have been threaded, ARG-Needle performs a final post-processing step called ARG normalization (see Methods).

To compute the threading instruction of a sample, ARG-Needle performs genotype hashing [24, 25] to rapidly detect a subset of candidate closest relatives within the ARG, then uses the ASMC algorithm [11] to compute pairwise TMRCA values and to select the closest sample at each genomic position. When all samples have been threaded to the ARG, ARG-Needle uses a fast post-processing step, which we call ARG normalization, that applies a monotonic correction to the node times of the ARG while leaving the topology unchanged. ARG-Needle builds the ARG in time approximately linear in sample size, depending on hashing parameters (see Methods).

We also introduce an extension of ASMC [11], called ASMC-clust, that builds genome-wide genealogies by forming a tree at each site using hierarchical clustering on pairwise TMRCAs output by ASMC. This approach scales quadratically with sample size and produces less correlated marginal trees, but the joint modeling of all pairs of samples leads to improved accuracy in certain scenarios. Additional details for the ARG-Needle and ASMC-clust algorithms are provided in Methods and Supplementary Note 1.

### Accuracy of ARG reconstruction in simulated data

We used extensive coalescent simulations to compare the accuracy and scalability of ARG-Needle, ASMC-clust, Relate [17], and tsinfer [15] (Fig. 2 and Supplementary Figs. S1-S3). We generated synthetic array data sets of up to 32,000 haploid samples using a European demographic model (see Methods) and measured ARG reconstruction accuracy. To this end, we considered several metrics used in the past to compare ARGs, including: the Robinson-Foulds distance [26], which reflects dissimilarities between the possible mutations that can be generated by two ARGs; the root mean squared error (RMSE) between true and inferred pairwise TMRCAs, which captures the accuracy in predicting allele sharing between individuals; and the Kendall-Colijn (KC) topology-only distance [27]. We found that the KC distance is systematically lower for trees containing polytomies (Supplementary Fig. S1b-d, see Methods and Supplementary Note 2), which are not output by Relate, ASMC-clust, or ARG-Needle by default. We therefore applied a simple heuristic to allow all methods to output polytomies (see Methods). As further discussed in Supplementary Note 2, although these three metrics capture the similarity between marginal trees and are in some cases interpretable in terms of accuracy in downstream analyses, they are not specifically developed for applications related to ARGs. We therefore also developed a new metric, called the ARG total variation distance, which generalizes the Robinson-Foulds distance to better capture the ability of a reconstructed ARG to predict mutation patterns that may be generated by the true underlying ARG (see Supplementary Note 2). Across these metrics, ARG-Needle and ASMC-clust achieved best performance for our primary array data simulations (Fig. 2a-d).

**Figure 2:**
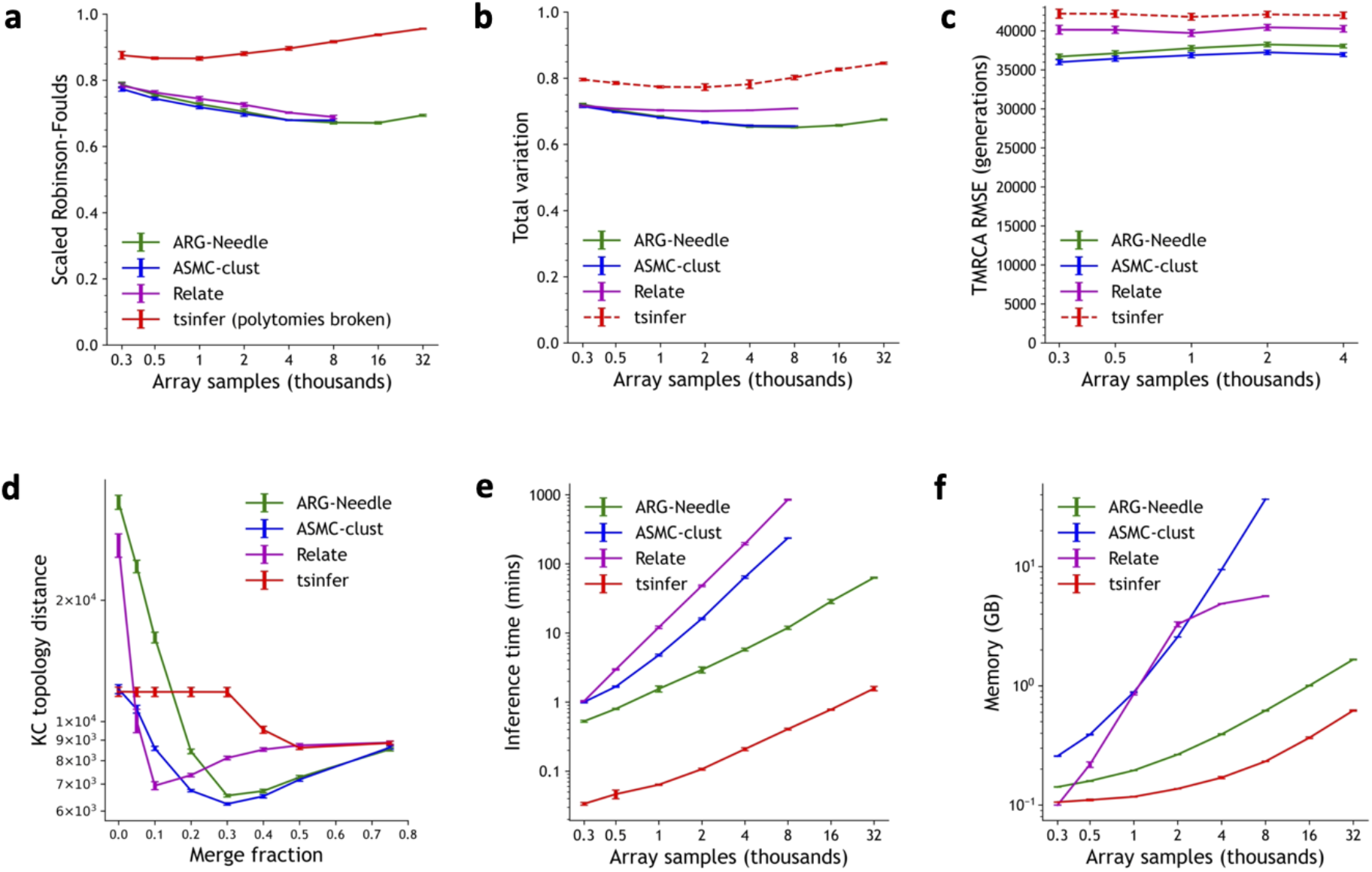
Comparison of ARG inference algorithms in simulation. We benchmark ARG inference performance for ARG-Needle, ASMC-clust, Relate, and tsinfer in realistic array data simulations across a variety of metrics related to accuracy and computational resources (lower values indicate better performance for all metrics), including **a**. the Robinson-Foulds distance, **b**. the ARG total variation distance (see Methods), **c**. pairwise TMRCA root mean squared error, **d**. the Kendall-Colijn topology-only metric, **e**. runtime, and **f**. peak memory. In **c**, we only run up to *N* = 4,000 samples. In **d**, we fix *N* = 4,000 samples and vary the fraction of branches that are merged to form polytomies, using a heuristic that preferentially merges branches that are less confidently inferred (see Supplementary Note 2). Both **c** and **d** involve 25 random seeds. All other examples use 5 random seeds and run up to 32,000 samples for ARG-Needle and ASMC-clust, and 8,000 samples for ASMC-clust and Relate due to runtime or memory constraints. Error bars represent 2 s.e. Dotted lines for tsinfer correspond to evaluation metrics which consider branch lengths. Relate’s default settings cap the memory for intermediate computations at 5 GB (see **f**). For additional simulations, see Supplementary Figs. S1-S2.

We next measured the speed and memory footprint of these methods. ARG-Needle requires lower computation and memory than Relate and ASMC-clust, which both scale quadratically with respect to sample size (Fig. 2e,f and Supplementary Fig. S1a). It runs slower than tsinfer but with a similar (approximately linear) scaling (also see Methods and Supplementary Note 1). Finally, we performed simulations in a variety of additional settings, including a constant demographic history, varying recombination rates, and with sequencing data (Supplementary Figs. S1-S2), as well as additional tests to measure the effects of ARG normalization (Supplementary Figs. S2-S3). ARG-Needle tended to achieve best performance across all accuracy metrics in array data, sometimes tied or in close performance with ASMC-clust or Relate. In sequencing data, ASMC-clust performed best on the ARG total variation and TMRCA RMSE metrics, with ARG-Needle and Relate close in performance, while Relate and tsinfer performed better on the Robinson-Foulds metric.

### A genealogical approach to linear mixed model analysis

Linear mixed models (LMMs) represent the state of the art for the analysis of polygenic traits, including heritability estimation [28, 29], polygenic prediction [30], and association [20, 31]. We devseloped an approach that uses the ARG of a set of genomes to perform mixed-linear-model association (MLMA [31]) of complex traits (see Methods). More in detail, we use an ARG built from an incomplete set of markers, in our case genotyping array data, to infer the presence of unobserved variants, and test these inferred variants for association within a mixed model framework. This increases association power in two ways: the ARG is used to uncover previously unobserved variants and the LMM utilizes estimates of genomic similarity across samples to model polygenicity, while also accounting for relatedness and population stratification [31]. This strategy generalizes approaches developed in the past that used haplotype sharing and genealogical relationships to improve association and fine-mapping [32, 33, 34, 25, 19, 13]. We refer to association analyses that test variants in the ARG as genealogy-wide association scans and to analyses that incorporate mixed-linear-model testing as ARG-MLMA. In simulations, we observed that genealogy-wide association and ARG-MLMA can achieve higher statistical power to detect signals that are linked to low frequency causal variants (MAF = 0.25%) than testing based on SNP array variants or variants imputed from a sequenced reference panel (Fig. 3a and Supplementary Fig. S4).

**Figure 3:**
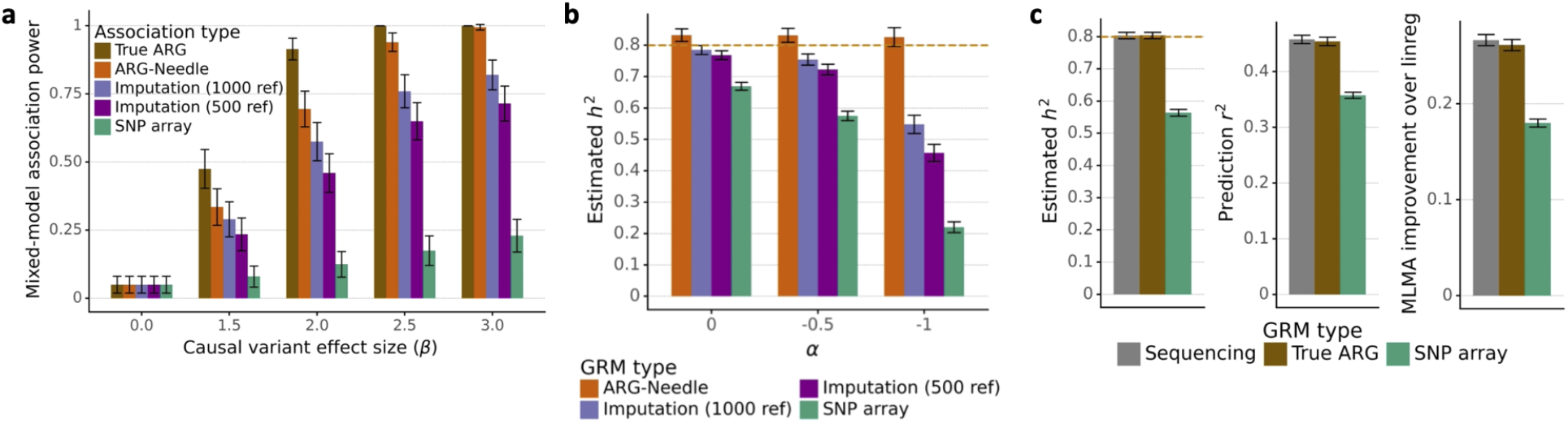
ARG-based analysis of simulated complex traits. **a**. Power to detect a low-frequency causal variant (MAF = 0.25%) in simulations of a polygenic phenotype. We compare ARG-MLMA of ground-truth ARGs and ARG-Needle inferred ARGs with MLMA of imputed and SNP array variants as we vary the effect size *β* (200 independent simulations of *h*^2^ = 0.8, *α* = -0.25, *N* = 2,000 haploid samples, and 22 chromosomes of 0.5 Mb each, see Methods). **b**. Heritability estimation using ARG-GRMs from ARG-Needle inference on SNP array data, compared to using imputed or array SNPs (5 simulations of 25 Mb, 5,000 haploid samples, *h*^2^ = 0.8, and varying *α*). **c**. ARG-GRMs computed using ground-truth ARGs perform equivalently to GRMs computed using sequencing data in heritability estimation, polygenic prediction, and mixed-model association. For association, we show the relative improvement in mean - log_10_(*p*) of MLMA compared to linear regression (see Methods). Heritability and prediction involve 5 simulations of 50 Mb, and association involves 50 simulations of 22 chromosomes, each of 2.5 Mb. In all cases, *N* = 10,000 haploid samples, *h*^2^ = 0.8, and *α* = -0.5. *ref* indicates the number of haploid reference samples used for imputation. Error bars represent 2 s.e. (from meta-analysis in the case of heritability estimation). Additional results are shown in Supplementary Figs. S4-S7.

We sought to verify that genomic variants that can be inferred using an ARG built from genotyping array data collectively capture more phenotypic variance than array markers alone. To this end, we developed strategies to use all variants that may be present in an ARG to obtain estimates of genomic similarity across individuals, which we aggregate in a genomic relatedness matrix (GRM, see Methods). We refer to GRMs built using this approach as ARG-GRMs and provide details of their derivation, properties, and computationally efficient estimation in Supplementary Note 3. We used ARG-GRMs to measure the amount of phenotypic variance captured by the ARG, performing LMM inference of narrow sense heritability (Supplementary Fig. S5a). In simulations, ARG-GRMs built using ARGs inferred by ARG-Needle in array data captured more phenotypic variance than GRMs built using array data, consistent with results observed for ARG-MLMA [28, 35, 36] (see Methods, Fig. 3b, and Supplementary Fig. S7). We also performed additional simulations to test whether the modeling of unobserved genomic variation using ARG-GRMs may be leveraged to obtain performance gains in other LMM analyses. We observed that ARG-GRMs built using true ARGs perform as well as GRMs computed using sequencing data in LMM-based heritability estimation, polygenic prediction, and association power (see Methods, Fig. 3c, and Supplementary Fig. S6). However, we note that applying these analyses to inferred ARGs and large data sets will require further methodological advances (see Discussion).

Overall, these experiments suggest that accurate genealogical inference combined with linear mixed models allows increasing association power by testing variants that are not well tagged using available markers while modeling polygenicity. The ARG may also be potentially utilized to obtain improved estimates of genomic similarity, to the benefit of additional LMM-based complex trait analyses.

### Genealogy-wide association scan of rare and ultra-rare variants in the UK Biobank

Using ARG-Needle, we built the genome-wide ARG from genotyping array data for 337,464 unrelated White British individuals in the UK Biobank (see Methods). We performed ARG-MLMA for standing height and 6 molecular traits, comprising alkaline phosphatase, aspartate aminotransferase, LDL/HDL cholesterol, mean platelet volume, and total bilirubin. To scale this analysis to the entire data set, we built on a recent method for large-scale MLMA [37, 22], which uses an array-based GRM to model polygenicity (see Methods and Discussion). We compared ARG-MLMA to standard MLMA testing of variants imputed using the combined Haplotype Reference Consortium (HRC) and UK10K reference panels (hereafter HRC+UK10K) [38, 39, 40], comprising ∼65K haploid samples. We filtered the imputed variants using standard criteria and focused on rare (0.01% ≤ MAF < 0.1%) and ultra-rare (MAF < 0.01%) genomic variants. We used a permutation-based approach [41] to establish genome-wide significance thresholds of *p* < 4.8 × 10^−11^ for ARG variants (sampled with mutation rate *μ* = 10^−5^) and *p* < 1.06 × 10^−9^ for imputed variants and performed extensive LD-based filtering to extract a stringent set of approximately independent associations (hereafter “independent associations”) resulting from each analysis (see Methods). To aid the localization and validation of these independent associations, we leveraged a subset of 138,039 individuals for whom whole exome sequencing (WES) data was also available (we refer to this data set as WES-138K). For each detected independent variant, we selected the exome sequenced variant with the largest correlation, which we refer to as its *WES partner*.

Applying this approach, we detected 133 independent signals using the ARG and 65 using imputation, implicating a set of 149 unique WES partners (Supplementary Tables 2-3). Of these WES variants, 38 were implicated using both approaches, confirming a common underlying source for these associations (Fig. 4a, for region-level results see Supplementary Fig. S8a). The fraction of WES partners uniquely identified using the ARG was larger among ultra-rare variants (84%) compared to rare variants (40%), reflecting a scarcity of ultra-rare variants in the sequenced HRC+UK10K panel. We observed a strong correlation between the phenotypic effects estimated in the 337K individuals using ARG-derived or imputed associations and those directly estimated for the set of WES partners in the WES-138K data set (Fig. 4b), with a stronger correlation (bootstrap *p* = 0.002) for ARG-derived variants (average 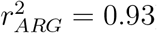) compared to imputed variants (average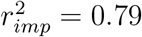). Only 73% of the WES partners for ARG-derived rare variant associations were significantly associated (at *p* < 5 × 10^−8^) in the smaller WES-138K data set, a proportion that dropped to 59% for ultra-rare variants. Variants detected using genealogy-wide association had a larger average phenotypic effect than those detected via imputation (bootstrap *p* < 0.0001; average 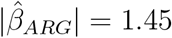; average 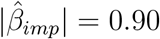), reflecting larger effects observed in ultra-rare variants. In addition, the set of WES partners implicated by either ARG or imputation were ∼2.3× enriched for missense variation, and ARG-derived WES partners were ∼4.5× enriched for loss-of-function variation compared to exome-wide variants of the same frequency (bootstrap *p* < 0.001, Fig. 4c), supporting their likely causal role.

**Figure 4:**
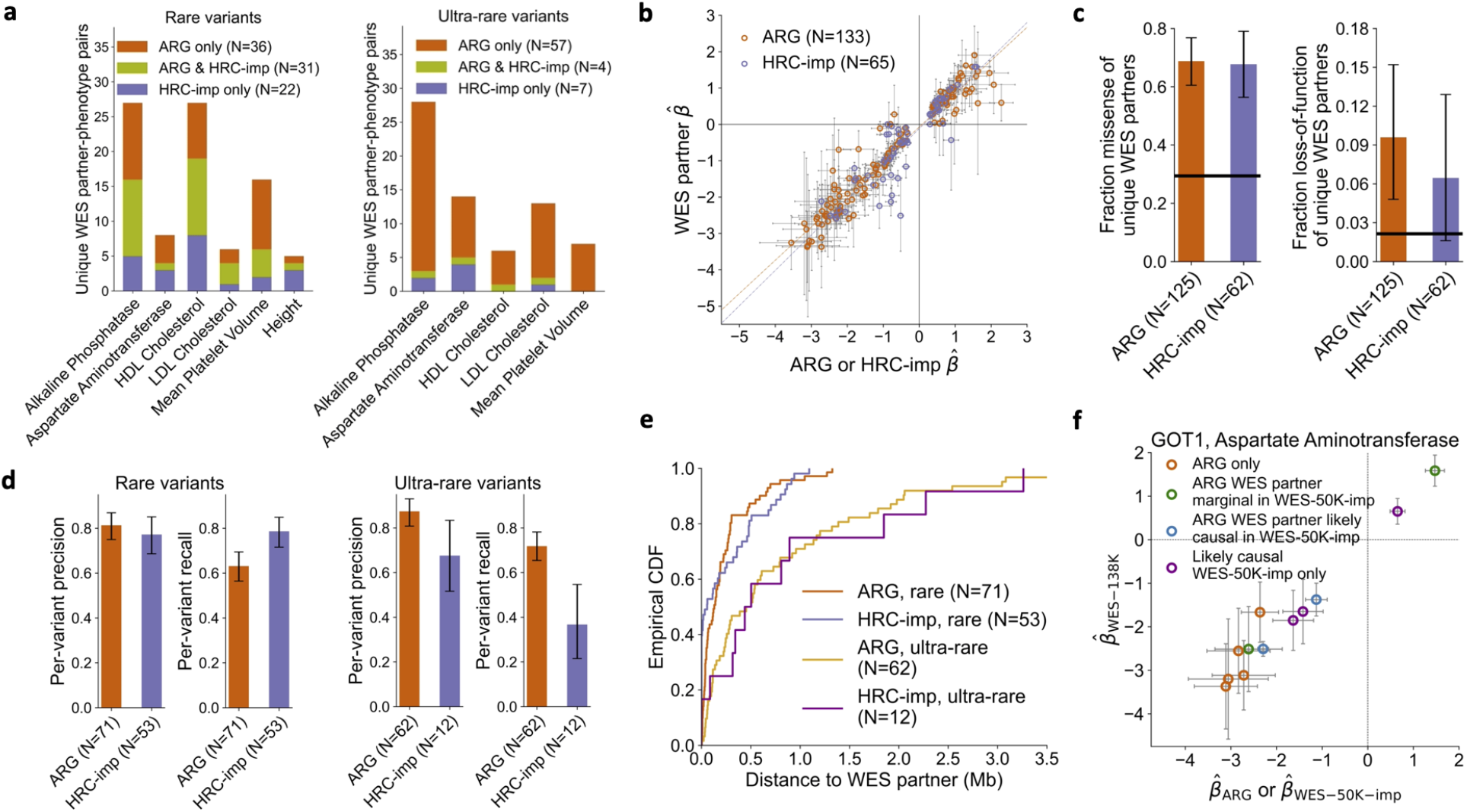
Association of ARG-derived and imputed rare and ultra-rare variants with 7 quantitative traits in UK Biobank. **a**. Counts of unique WES partners for ARG and HRC+UK10K imputed (“HRC-imp”) independent associations, partitioned by traits and frequency and showing overlap. Total bilirubin was not associated at these frequencies. **b**. Scatter plot of 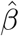 (estimated effect) for independent variants against 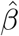 for their WES partners, with linear model fit. **c**. Fraction of missense and loss-of-function variants for the unique WES partners of independent variants. Horizontal black lines represent genome-wide averages. **d**. Average per-variant precision and recall of predicting WES carrier status, partitioned by frequency. **e**. Cumulative distribution function for the distance between independent variants and their WES partners. **f**. Scatter plot of 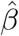 for ARG-derived independent variants with aspartate aminotransferase in the *GOT1* gene against 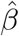 for their WES partners. We color points based on whether the WES partner is likely causal in WES-50K-imp (imputation from WES-50K [42]), not likely causal but marginally significant in WES-50K-imp, or not marginally significant in WES-50K-imp (“ARG only”). We also plot the 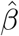 for the additional likely causal variants in WES-50K-imp against the 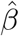 in WES-138K. Error bars represent 1.96 s.e. in **b** and **f** and represent bootstrap 95% confidence intervals in **c** and **d**. Additional results are shown in Fig. S8.

We also used the WES-138K data set to measure the extent to which carrying an associated ARG-derived or imputed variant is predictive of carrying the corresponding sequence-level WES partner variant (Fig. 4d). We quantified this using variant-level precision and recall statistics (see Methods). ARG-derived and imputed rare variants had similar levels of variant-level precision, while imputation had higher recall (bootstrap *p* = 0.0009). For ultra-rare variants, ARG-derived signals performed better than imputed variants for both precision (bootstrap *p* = 0.01) and recall (bootstrap *p* < 0.001). Similarly, ARG-derived and imputed rare variants provided comparable tagging for their WES partners (Supplementary Fig. S8b). ARG-derived ultra-rare variants, on the other hand, provided stronger tagging compared to imputed ultra-rare variants (average *r*_*ARG*_ = 0.77, average *r*_*imp*_ = 0.42, bootstrap *p* < 0.001; average *r*_*ARG*_ = 0.72 for combined rare and ultra-rare variants). Compared to ARG-derived variants, genotype imputation has the advantage that associated variants that are sequenced in the reference panel may be directly localized in the genome. We found that for 21/53 of rare and 2/12 of ultra-rare independent imputation signals the WES partner had been imputed (having the same physical position as the associated variant), while the remaining signals likely provide indirect tagging for underlying variants. Comparing ARG-derived and imputed variants in terms of the distance to their WES partners, however, revealed similar distributions (Fig. 4e and Supplementary Fig. S8c). This suggests that genealogy-wide associations have the same spatial resolution as associations obtained using genotype imputation in cases where the variant driving the signal cannot be directly imputed, unless sequencing data is available to further localize the signal, as done here using WES partners.

A recent study leveraged exome sequencing data from a subset of ∼50K participants (hereafter WES-50K) to perform genotype imputation for ∼459K European samples, achieving association power equivalent to exome sequencing of ∼250K samples [42]. We considered the set of WES partners implicated using ARG-derived independent signals and not using HRC+UK10K imputation, and found that, of these, 14/30 rare and 28/54 ultra-rare variants were also detected as likely-causal associations (at *p* < 5 × 10^−8^) in [42] (see Supplementary Table 2). For the remaining 42 independent associations that are detected using the ARG but are not reported in [42], WES partners are often very rare variants (median MAF = 3.8 × 10^−5^; Supplementary Fig. S8d) of large phenotypic effect (median |*β*| = 1.12). These variants are difficult to impute, even when large reference panels are available: 18/42 such variants were absent or singletons in the WES-50K data set used for imputation or had poor imputation quality score. The set of associations uniquely detected using the ARG often extended allelic series at key genes linked to the analyzed traits. For instance, restricting to loss-of-function or missense WES partners for independent ARG signals that are not present or marginally significant in [42], 5 novel variants with aspartate aminotransferase are mapped to the *GOT1* gene (Fig. 4f) and 3 with alkaline phosphatase are mapped to *ALPL* (Supplementary Fig. S8e). A subset of strong independent associations uniquely detected by the ARG had weak correlation with their WES partners (e.g. a signal for aspartate aminotransferase with *p* = 3.2 × 10^−39^, ARG-MAF = 0.00053, WES partner *r* = 0.2, WES-138K MAC = 6, WES-50K MAC = 1). These signals may tag variants, such as structural or regulatory variation, that are unobserved in the WES-138K data set.

In summary, a genealogy-wide association scan using an ARG inferred from common SNPs revealed more independent rare and ultra-rare associations than a scan performed using genotype data imputed from a reference panel comprising ∼65K sequenced haplotypes. Using genealogy-wide association, we also detected ultra-rare variant associations that were not detected using imputation from a subset of ∼50K exome sequenced participants from the same cohort. ARG-derived associations accurately predicted the effect of underlying sequencing variants as well as the subset of carrier individuals. Leveraging a subset of exome sequenced samples enabled further fine-mapping of several genomic regions implicated using the ARG.

### Genealogy-wide association for low and high frequency variants

Lastly, we performed genealogy-wide association for low (0.1% ≤ MAF < 1%) and high (MAF ≥ 1%) frequency variants. Variants within these frequencies are more easily imputed using reference panels that are not necessarily large and population-specific. Consistent with this, extending our previous analysis to consider low-frequency variants yielded a similar number of independent associations for ARG-derived and HRC+UK10K-imputed variants (*N*_*ARG*_ = 102, *N*_*imp*_ = 100, see Supplementary Tables 4-5, Supplementary Fig. S9a-c). Associations detected using the ARG had slightly larger effects compared to those found using imputation (bootstrap *p* = 0.045; average |*β*_*ARG*_| = 0.31; average |*β*_*imp*_| = 0.27) but provided lower tagging to their WES partners (bootstrap *p* < 0.001; average *r*_*ARG*_ = 0.56; average *r*_*imp*_ = 0.73), reflecting the large fraction (42/100) of WES partners that could be directly imputed.

We hypothesized that although imputation of higher frequency variants is generally accurate, branches in the marginal trees of the ARG may tag underlying causal variants better than the set of available polymorphic markers, revealing complementary signal. To test this, we performed MLMA for height using HRC+UK10K imputed variants filtered using the criteria used in [40], including MAF > 0.1% and info score > 0.3 (see Methods). For these variants, we established a permutation-based genome-wide significance threshold of 4.5 × 10^−9^ (95% CI: [2.2 × 10^−9^, 9.6 × 10^−9^]). To facilitate direct comparison, we selected ARG-MLMA parameters that correspond to a comparable genome-wide significance threshold (see Methods). We thus ran ARG-MLMA by sampling mutations from the ARG at rate *μ* = 1 × 10^−5^ and restricting to MAF > 1%, for which we obtained a permutation-based threshold of 3.4 × 10^−9^ (95% CI: [2.4 × 10^−9^, 5 × 10^−9^]). In downstream analyses, we adopted a significance threshold of 3 × 10^−9^.

We assessed the total number of 1 Mb regions that contain an association (*p* < 3 × 10^−9^) for either genotype array, imputed, or ARG-derived variants. We found that ARG-MLMA detected 98.9% of regions found by both SNP array and imputation, as well as 71% of regions found by imputation but not array data, and detected an additional 8% of regions not found by either imputation or array data (Supplementary Fig. S9d). A significant fraction (54/92, permutation *p* < 0.0001) of regions identified using the ARG but not HRC+UK10K-imputation contained significant (*p* < 3 × 10^−9^) associations in a larger meta-analysis by the GIANT Consortium [43] (*N* ≈ 700K), which comprised the UK Biobank and additional cohorts. By inspecting associated loci, we observed that ARG-MLMA captures association peaks and haplotype structure found using genotype imputation but not array data (Fig. 5a-b,f, Supplementary Fig. S11d-h, Supplementary Fig. S10) as well as association peaks uniquely identified using ARG-MLMA (Fig. 5e, Supplementary Fig. S11a-c).

**Figure 5:**
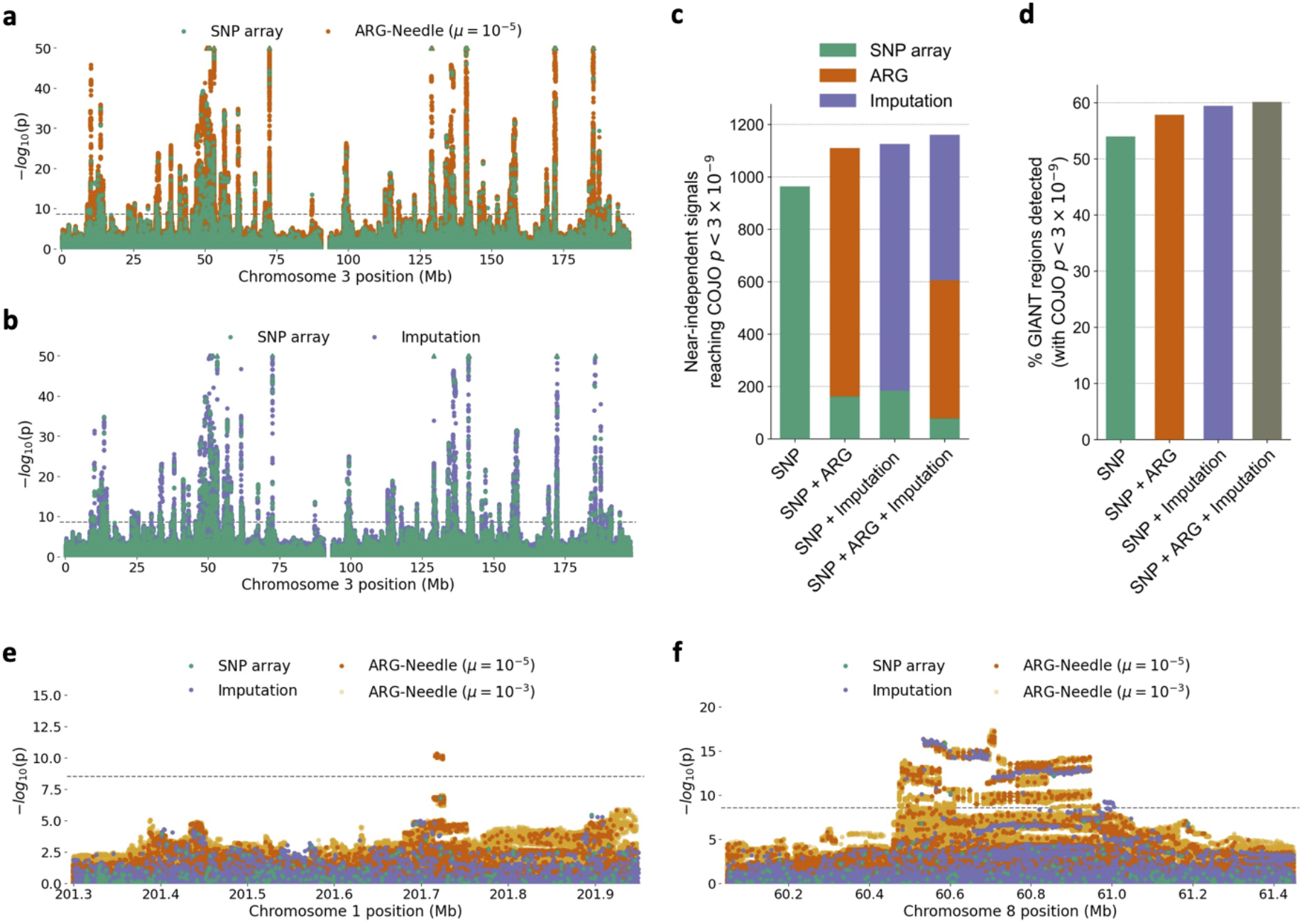
Genealogy-wide association of higher frequency variants with height in UK Biobank. **a**-**b**. Chromosome 3 Manhattan plots showing MLMA of ARG-Needle on SNP array data vs. array SNPs (**a**) and HRC+UK10K imputed variants vs. array SNPs (**b**). **c**-**d**. Near-independent associations (COJO *p* < 3 × 10^−9^) when considering array SNPs alone, array SNPs and ARG-Needle variants, array SNPs and imputed variants, and all three types of variants. **c**. Total number of independent variants found and attribution based on data type. **d**. Percent of 1 Mb regions containing COJO associations in a GIANT consortium meta-analysis of ∼700K samples that are detected using our methods. **e**-**f**. Manhattan plots of two example loci. **e**. An association peak found by ARG-MLMA that was significant (*p* < 3 × 10^−9^) in the GIANT meta-analysis. **f**. ARG-MLMA detects haplotype structure that is found using imputation, while indicating a new association peak. For the Manhattan plots, the order of plotting is ARG-Needle with *μ* = 10^−3^ (used for follow-up), then ARG-Needle with *μ* = 10^−5^ (used for discovery), then imputation, then SNP array variants on top. Dotted lines correspond to *p* = 3 × 10^−9^ (see Methods) and triangles indicate associations with *p* < 10^−50^. See also Supplementary Figs. S10-S11.

We performed LD-based filtering as well as conditional and joint (COJO [44]) association analysis (Fig. 5c, see Methods) to obtain a set of approximately independent association signals. Analyses including either or both ARG-derived and imputed variants in addition to array markers resulted in an increase in the number of significant (*p* < 3 × 10^−9^) COJO variants (*N*_*SNP*_ = 964, *N*_*SNP*+*ARG*_ = 1,110, *N*_*SNP*+*imp*_ = 1,126, *N*_*SNP*+*ARG*+*imp*_ = 1,161). The fraction of COJO-associated array markers was substantially reduced by the inclusion of ARG-derived or imputed variants, suggesting that both ARG and imputation provide better tagging of underlying signal than array markers alone. ARG-derived and imputed variants, on the other hand, resulted in comparable proportions of COJO associations when jointly analyzed (Fig. 5c). We sought to validate this increase in the number of independent signals by leveraging COJO association summary statistics from the GIANT analysis [43]. To this end, we considered the set of 1 Mb regions harboring significant COJO associations and observed that the additional COJO signals detected when including ARG-derived or imputed variants concentrated within regions that also harbor significant (*p* < 3 × 10^−9^) COJO signal in the GIANT analysis (Fig. 5d and Supplementary Fig. S9e).

In summary, higher frequency variant analysis using the ARG inferred by ARG-Needle from SNP array data revealed associated haplotypes and peaks that were not found through association of array data alone, and complemented genotype imputation in detecting independent association signals.

## Discussion

We developed ARG-Needle, a method for accurately inferring genome-wide genealogies from genomic data that scales to large biobank data sets. We performed extensive simulation-based benchmarks, showing that ARG-Needle is both accurate and scalable when applied to ascertained genotyping array and sequencing data. We also developed a framework that combines inferred genealogies with linear mixed models to increase statistical power to detect phenotypic associations, particularly for unobserved rare and ultra-rare variants, and showed that this strategy may be utilized in analyses of heritability and polygenic prediction. We applied these new tools to build genome-wide ARGs from genotyping array data for 337,464 UK Biobank individuals and performed a genealogy-wide association scan for height and 6 molecular phenotypes. Using the inferred ARG, we detected more associations to rare and ultra-rare variants of large effect than using genotype imputation from ∼65,000 ancestry-matched sequenced haplotypes, down to a frequency of ∼4 × 10^−6^. We validated these signals using 138,039 exome sequencing samples, showing that they strongly tag underlying variants that are enriched for predicted missense and loss-of-function variation. Associations detected using the ARG overlap with and extend fine-mapped associations detected using genotype imputation based on the sequencing of a large fraction of analyzed individuals. Applied to the analysis of higher frequency variants, the ARG revealed haplotype structure and independent signals complementary to those obtained using genotype imputation.

These results highlight connections between genealogical modeling and linear mixed model analysis of complex traits. When causal variants are not directly observed, genome-wide association analyses rely on the correlation between available markers and underlying variation [45] and the MLMA strategy is used to account for polygenicity, relatedness, and population structure [31]. In genealogy-wide association, on the other hand, the signal of LD is amplified by further modeling the signature of past recombination events to directly infer the presence of hidden genomic variation. Through ARG-GRMs, the genealogy may be used as a further route to amplifying power in MLMA, by obtaining better estimates of genomic similarity and shared polygenic effects.

These analyses also demonstrate that genealogical inference provides a complementary strategy to genotype imputation approaches, which rely on haplotype sharing between the analyzed samples and a sequenced reference panel to extend the set of available markers. Imputation has been successfully applied in the analysis of complex traits [4, 35], but its efficacy, particularly in the study of rare and ultra-rare variants, hinges on the availability of large, population-specific sequencing panels [39, 42]. Although sequencing data sets are rapidly growing [46], they are not widely available for all populations. Genealogy-wide association based on ARGs inferred from incomplete genomic data may therefore offer new avenues to better utilize genomic resources for groups that are underrepresented in modern sequencing studies [47].

We highlight several limitations and directions of future development for this work. First, although genealogy-wide association accurately detects the subset of individuals carrying independently associated variants, the signal is usually localized within a genomic region, whereas genotype imputation may implicate individual variants if they are present in the sequenced reference panel. However, when sequencing data is available, it may be utilized to further localize ARG-derived signals, as done using WES partners in our analyses. Second, although we have shown in simulation that ARG-GRMs built from true ARGs may be used to estimate heritability, perform polygenic prediction, and increase association power, preliminary work we performed suggests that real data applications of this approach will require further methodological improvements to achieve sufficient accuracy and scalability. To this end, it may be possible to build on recent advances in methods for highly scalable linear mixed model algorithms and for improved modeling of complex architectures and environmental factors [48, 49, 50, 51]. Third, although we have focused on leveraging an ARG inferred from array data alone, ARG-Needle currently enables building an ARG using a mixture of sequencing and array data. This approach may be used to perform ARG-based genotype imputation, which is likely to improve upon approaches that do not accurately model the TMRCA between target and reference samples [52]. We performed preliminary simulation-based analyses using this strategy, and obtained promising results (see Methods, Supplementary Fig. S12). Fourth, reconstructing biobank-scale ARGs will likely aid the study of additional evolutionary properties of disease-associated variants, including analyses of natural selection acting on complex traits [11, 53, 54], which we have not explored in this work. Finally, our analysis focused on the UK Biobank data set, which provides an excellent testbed due to the large volumes of high-quality data of different types that are available for validation. Future applications of our methods will involve analysis of cohorts that are less strongly represented in current sequencing studies. Nevertheless, we believe that the results described in this work represent an advance in large-scale data-driven genealogical inference and provide new tools for the analysis of complex traits.

## Methods

### ARG-Needle and ASMC-clust algorithms

We introduce two algorithms to construct the ARG of a set of samples, called ARG-Needle and ASMC-clust. Both approaches leverage output from the ASMC algorithm [11], which takes as input a pair of genotyping array or sequencing samples and outputs a posterior distribution of the time to most recent common ancestor (TMRCA) across the genome. These pairwise TMRCAs are equivalent to an ARG between two samples, which ARG-Needle and ASMC-clust use to assemble the ARG for all individuals.

ASMC-clust runs ASMC on all pairs of samples and performs hierarchical clustering of TMRCA matrices to obtain an ARG. At every site, we apply the unweighted pair group method with arithmetic mean (UPGMA) clustering algorithm [55] on the *N* × *N* posterior mean TMRCA matrix to yield a marginal tree. We combine these marginal trees into an ARG, using the midpoints between sites’ physical positions to decide when one tree ends and the next begins. By using an *O*(*N* ^2^) implementation of UPGMA [56, 57], we achieve a total runtime and memory complexity of *O*(*N* ^2^*M*). We also implement an extension that achieves *O*(*NM*) memory at the cost of increased runtime (see Supplementary Note 1).

ARG-Needle starts with an empty ARG and repeats three steps to add additional samples to the ARG: (1) detecting a set of closest genetic relatives via hashing, (2) running ASMC, and (3) “threading” the new sample into the ARG (Fig. 1). Given a new sample, the first step performs a series of hash table queries to determine the candidate closest samples already in the ARG [24]. We divide up the sites present in the genetic data into non-overlapping “words” of *S* sites and store hash tables mapping from the possible values of the *i*th word to the samples that carry that word. We use this approach to rapidly detect samples already in the ARG that share words with the target sample and return the top *K* samples with the most consecutive matches. A tolerance parameter *T* controls the number of mismatches allowed in an otherwise consecutive stretch. We also allow the top *K* samples to vary across the genome due to recombination events, by partitioning the genome into regions of genetic distance *L*. Assuming this results in *R* regions, the hashing step outputs a matrix of *R* × *K* sample IDs containing the predicted top *K* related samples over each region.

The sample IDs output by the first step inform the second step of ARG-Needle, in which ASMC is run over pairs of samples. In each of the *R* regions, ASMC computes the posterior mean and maximum a posteriori (MAP) TMRCA between the sample being threaded and each of the *K* candidate most related samples. We add up to 1.0 cM on either side of the region, to provide additional context for the ASMC model.

In the third step, ARG-Needle finds the minimum posterior mean TMRCA among the *K* candidates at each site of the genome. The corresponding IDs determine which sample in the ARG to thread to at each site. Because the posterior mean assumes continuous values and changes at each site, we average the posterior mean over neighboring sites where the ID to thread to and the associated MAP remain constant. This produces piecewise constant values which determine how high above the sample to thread, with changes corresponding to inferred recombination events. The sample is then efficiently threaded into the existing ARG, utilizing custom data structures and algorithms.

Of the three steps of ARG-Needle, the second step (ASMC) dominates runtime for small *N* and the first step (hashing) dominates runtime for large *N*. The parameters *K, L*, and *S* each carry an accuracy versus runtime tradeoff, with large *K*, small *L*, and small *S* leading to a more accurate but slower algorithm. We decided on default parameters of *K* = 64 and *L* = 0.5 cM for array data and *K* = 64 and *L* = 0.1 cM for sequencing data. In experiments with simulated genotypes, we fixed the hash word size at *S* = 16. As threading proceeds, increasingly close relationships and thus increasingly long shared haplotypes are detected between the sample and other individuals in the ARG. In real data analysis we therefore increase *S* as threading proceeds, which reduces computational cost without a significant loss in accuracy. We also implemented a larger primary hash word size *S*_1_, leveraged for increased speed, and a smaller “backup hash word size” *S*_2_, used for samples where more fine-grained hashing was needed (see Supplementary Note 1). We set *S*_1_ = 16 and *S*_2_ = 8 when threading the first 50K individuals and set *S*_1_ = 64 and *S*_2_ = 16 for the remaining 287K individuals. We set *T* = 1 in simulations and real data analyses to enable robustness to a single genotyping error or recent mutation event in otherwise closely related samples.

For additional details on all three steps in the ARG-Needle algorithm, see Supplementary Note 1.

### ARG normalization

ARG normalization applies a monotonically increasing mapping from existing node times to transformed node times (similar to quantile normalization), further utilizing the demographic prior provided in input. We compute quantile distributions of node times in the inferred ARG as well as in 1,000 independent trees simulated using the demographic model provided in input under the single-locus coalescent. We match the two quantile distributions and use this to rewrite all nodes in the inferred ARG to new corresponding times (see Supplementary Note 1). ARG normalization preserves the time-based ordering of nodes and therefore does not alter the topology of an ARG. It is applied by default to our inferred ARGs and optionally to ARGs inferred by Relate and tsinfer (see Supplementary Figs. S2-S3).

### Simulated genetic data

We used the msprime coalescent simulator [58] to benchmark ARG inference algorithms. For each run, we first simulated sequence data with given physical length *L* for *N* haploid individuals. Our primary simulations used a mutation rate of *μ* = 1.65 × 10^−8^ per base pair per generation, a constant recombination rate of *ρ* = 1.2 × 10^−8^ per base per generation, and a demographic model inferred using SMC++ on CEU 1,000 Genomes samples [10]. These simulations also output the simulated genealogies, which we refer to as “ground-truth ARGs” or “true ARGs”.

To obtain realistic SNP data, we subsampled the simulated sequence sites to match the genotype density and allele frequency spectrum of UK Biobank SNP array markers (chromosome 2, with density defined using 50 evenly spaced MAF bins). When running ASMC, we used decoding quantities precomputed for version 1.1, which were obtained using a European demographic model and UK Biobank SNP array allele frequencies [11].

### Comparisons of ARG inference methods

In comparisons of ARG inference methods, we used simulations with *L* = 1 Mb for sequencing data and *L* = 5 Mb for array data. We ran Relate with the mutation rate, recombination rate, and demographic model used in simulations. Relate includes a default option, which we kept, that limits the memory used for storing pairwise matrices to 5 GB. For metrics which involve both topology and branch lengths, we used the default branch lengths output by tsinfer. Because these are not intended to be used as accurate estimates, we used dotted lines for tsinfer in Fig. 2b-c and Supplementary Fig. S2. For each choice of sample size, we generated genetic data using either 5 or 25 random seeds and applied ARG-Needle, ASMC-clust, Relate, and tsinfer to infer ARGs. Due to scalability differences, we ran ASMC-clust and Relate in up to *N* = 8,000 haploid samples (*N* = 4,000 for sequencing) and ARG-Needle and tsinfer up to *N* = 32,000 haploid samples. We ran all analyses on Intel Skylake 2.6 GHz nodes on the Oxford Biomedical Research Computing cluster.

A commonly used metric for comparing topologies of trees is the Robinson-Foulds metric [26], which counts the number of unique mutations that can be generated by one tree but not the other (lower implies higher accuracy). Because the presence of polytomies can skew this metric, we randomly break polytomies of tsinfer-inferred ARGs before comparing with Robinson-Foulds, as was done in [15]. We report a genome-wide average of the Robinson-Foulds metric, where we divide by the maximum possible value of 2*N* - 4 to obtain a rescaled quantity between 0 (minimum) and 1 (maximum).

We generalized the Robinson-Foulds metric to measure mutational dissimilarity while considering branch lengths, to better capture the accuracy in predicting unobserved variants using an inferred ARG. To this end, we consider the probability distribution of mutations induced by uniform sampling over an ARG and compare the resulting distributions for the true versus inferred ARG using the total variation distance, a common metric for comparing probability measures. Polytomies do not need to be broken using this metric, as they simply concentrate the probability mass on fewer predicted mutations. We refer to this metric as ARG total variation distance (see Supplementary Note 2 for further details).

For pairwise TMRCA comparisons, at each position we compute the TMRCA of each pair of samples in the true ARG and the inferred ARG, resulting in two vectors of length *N*(*N* - 1)/2. We average the squared Euclidean distance between these vectors over all positions, then take a square root, resulting in a pairwise TMRCA root mean squared error (RMSE).

We also considered the Kendall-Colijn (KC) topology-only distance to compare ARG topologies, calculated between marginal trees of two ARGs and averaged over all positions. We observed that the performance of methods that output binary trees (Relate, ASMC-clust, and ARG-Needle) under this metric significantly improved when we selected inferred branches at random and collapsed them to create polytomies (Supplementary Fig. S1c), suggesting that the KC distance tends to reward inferred ARGs that contain polytomies. Consistent with [15], we also observed that when polytomies are randomly broken, tsinfer’s performance on the KC distance deteriorates (Supplementary Fig. S1b). We therefore developed a heuristic approach to form polytomies in inferred ARGs by collapsing branches based on their size and height, which capture the confidence in the inferred tree branches. We applied this heuristic to all methods to compare their performance under the KC topology-only distance while accounting for different prevalence of polytomies in the inferred ARGs (Fig. 2d, Supplementary Fig. S1d).

Supplementary Note 2 provides further details on the computation of these metrics and their interpretation in the context of ARG inference and downstream analyses.

### ARG-based mixed linear model association (ARG-MLMA)

We developed an approach to perform mixed linear model association of variants extracted from the ARG, which we refer to as ARG-MLMA. In this approach we sample mutations from a given ARG using a specified rate *μ* and apply a mixed model association test to these variants. The choice of *μ* may be used to decrease the number of tests performed, while also accounting for uncertainty of the inferred variants within the ARG.

For simulation experiments (Fig. 3a and Supplementary Fig. S4) we traversed the ARG and wrote out all possible mutations to disk, which is equivalent to adopting a large value of *μ*, and used GCTA to perform LMM testing of these mutations. When comparing against ARG-based linear regression, we tested the same written mutations using PLINK. While the ARG-MLMA approach can in principle be combined with ARG-GRMs, for these simulations we used a GRM built using array markers to model polygenicity (Supplementary Fig. S4b), as done in larger UK Biobank analyses. We used sequencing variants from chromosomes 2-22 to form a polygenic background with narrow-sense heritability *h*^2^ = 0.8 and negative selection parameter [59] *α* = -0.25. In detail, we drew effects *β*_*i*_ ∼ *𝒩*(0, [*p*_*i*_(1 - *p*_*i*_)]^*α*^), computed *y*_*g*_ = ∑_*i*_ *β*_*i*_*x*_*i*_ using unnormalized haploid genotypes *x*_*i*_, and scaled *y*_*g*_ to have variance *h*^2^. We then added a single causal sequencing variant on chromosome 1 (chosen at random from those with allele frequency *p* ∈ {0.01, 0.005, 0.0025}) with effect size *β* and added independent normally-distributed noise for the remaining environmental variance. We varied the value of *β* and measured association power for each method as the fraction of runs (out of 200) detecting a significant association on chromosome 1. Significance thresholds for each method were calibrated to yield a family-wise error rate of 0.05 under the null condition *β* = 0. We compared between association of array data, imputed data, the true ARG, and an ARG inferred by ARG-Needle using array data (Fig. 3a and Supplementary Fig. S4). For each method, we tested using both linear regression and MLMA with a leave one chromosome out (LOCO) GRM built from array markers on chromosomes 2-22.

For ARG-MLMA analyses in the UK Biobank we adopted *μ* = 10^−5^, also adding variants sampled with *μ* = 10^−3^ to locus-specific Manhattan plots to gain further insights in the association regions. To achieve greater scalability, we leveraged the BOLT-LMM software package [37, 22], using the following procedure. We first regressed out covariates from the phenotype, then ran BOLT-LMM on SNPs from all chromosomes to extract BOLT-LMM’s calculated calibration factor. We additionally ran BOLT-LMM 22 times, once with each chromosome excluded, and extracted estimated prediction effects (`--predBetasFile` flag) to form LOCO polygenic predictors. We then obtained LOCO residuals by subtracting these LOCO predictions from the processed phenotype. We finally used ARG-Needle to test clades of the ARG for association against these residuals, traversing the ARG and calculating BOLT-LMM’s non-infinitesimal test statistics for each clade of interest. Our methods include runtime optimizations for sparse clades, as well as options to sample clades based on MAF or a mutation rate.

### Construction of ARG-GRMs

We consider *N* haploid individuals, *M* sites, and genotypes *x*_*ik*_ for individual *i* at site *k*, where variant *k* has mean *p*_*k*_. We assume an infinitesimal genetic architecture so that the genetic component of a trait is given by *g*_*i*_ = ∑_*k*_ *β*_*k*_*x*_*ik*_, where *β*_*k*_ is drawn with mean zero and MAF-dependent variance proportional to (*p*_*k*_(1 - *p*_*k*_))^*α*^, where *p*_*k*_ is the MAF of variant *k* and *α* captures the strength of negative selection [28, 59]. Using available markers, a common estimator for the *ij*-th entry of the *N*×*N* genomic relatedness matrix (GRM [21]) may be computed as

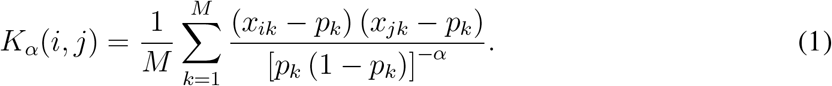

Given an ARG, we compute the ARG-GRM as the expectation of the marker-based GRM that would be obtained using sequencing data, i.e. when all variants are observed, assuming that mutations are sampled uniformly over the area of the ARG. The rationale of this approach is that when sequencing data is not available but an accurate ARG can be estimated from an incomplete set of markers, the ARG-GRM may provide a good estimate for the sequence-based GRM. We briefly describe how ARG-GRMs are derived from the ARG for the special case of *α* = 0. We discuss the more general case and provide further derivations in Supplementary Note 3.

Assuming *α* = 0, (1) is equivalent (up to invariances described in Supplementary Note 3) to the matrix containing the Hamming distance between the sequences of pairs of samples, given by

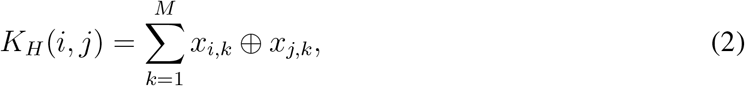

where ⊕ refers to the XOR function. Assume there are *L* total base pairs in the genome, a constant mutation rate per base pair and generation of *μ*, and denote the TMRCA of *i* and *j* at base pair *k* with *t*_*ijk*_ (in generations). The *ij*-th entry of the ARG-GRM is obtained as the expected number of mutations that are carried by only one of the two individuals, which corresponds to the expected hamming distance for sequences *i* and *j*:

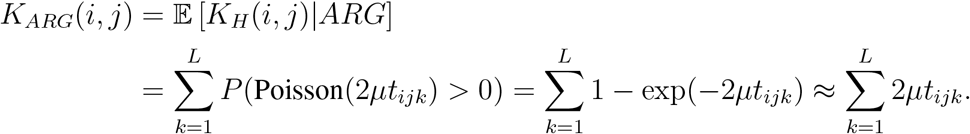

Computing the above ARG-GRM requires iterating over the entire ARG (Supplementary Fig. S5a). We instead computed a Monte Carlo ARG-GRM by uniformly sampling new mutations on the ARG with a high mutation rate and using these mutations to build the ARG-GRM, using (1). We used simulations to verify that Monte Carlo ARG-GRMs converge to exactly computed ARG-GRMs for large mutation rates, saturating at *μ* ≈ 1.65 × 10^−7^ (Supplementary Fig. S5b-c), the default value we used for ARG-GRM computations. Stratified Monte Carlo ARG-GRMs may also be computed by partitioning the sampled mutations based on e.g. allele frequency, LD, or allele age [60, 35, 61, 29]. For simulations adopting MAF-stratification (e.g. Supplementary Fig. S6g), we used MAF boundaries given by {0, 0.01, 0.05, 0.5}, normalized genotypes using *α* = -1, and then provided the MAF-stratified ARG-GRMs in input to GCTA [21].

### ARG-GRM simulation experiments

We simulated polygenic traits from haploid sequencing samples for various values of *h*^2^ and *α*. We varied the number of haploid samples *N* but fixed the ratio *L*/*N* throughout experiments, where *L* is the genetic length of the simulated region. For heritability and polygenic prediction experiments, we adopted *L*/*N* = 5 × 10^−3^ Mb/individuals. For association experiments, we simulated a polygenic phenotype from 22 chromosomes, with each chromosome consisting of equal length *L*/22 and *L*/*N* = 5.5 × 10^−3^ Mb/individuals. Mixed-model prediction *r*^2^ and association power may be roughly estimated as a function of *h*^2^ and the ratio *N*/*M*, where *M* is the number of markers [62, 30, 37]. We thus selected values of *M* and *L* such that the *N*/*M* ratio is kept close to that of the UK Biobank (*L* = 3 × 10^3^ Mb, *N* ≈ 6 × 10^5^).

We computed GRMs using ARGs, SNP data, imputed data, and sequencing data and provided them in input to GCTA with the simulated phenotype. For heritability estimation, we ran GCTA with flags `--reml-no-constrain` and `--reml-no-lrt`. For polygenic prediction, we ran leave-one-out prediction using cvBLUP [63] within GCTA, then computed *r*^2^ between the resulting predictions and the phenotype. For ARG-GRM association experiments (Fig. 3c and Supplementary Fig. S6c,f), we performed MLMA of array data SNPs, testing each chromosome while using a LOCO GRM built on the other 21 chromosomes. We measured power improvement as the relative increase of mean - log_10_(*p*) for MLMA compared to linear regression of array data SNPs and compared ARG-GRMs to GRMs of array and sequencing data.

We observed that MAF-stratification for ARG-GRMs of true ARGs enabled robust heritability estimation and polygenic prediction if *α* is unknown (Supplementary Fig. S6g). In experiments involving inferred ARGs (Fig. 3b and Supplementary Fig. S7), we applied MAF-stratification for ARG-Needle ARGs and imputed data, but not for SNP data, for which GCTA did not converge. In other experiments (Fig. 3c and Supplementary Fig. S6a-f) we used the true value of *α* to build GRMs. Imputed data comparisons used IMPUTE4 [40].

### ARG-based genotype imputation

Given a collection of sequencing and array samples, we perform ARG-based imputation as follows (see Supplementary Fig. S12a). We use ARG-Needle in sequencing mode to first thread the sequencing samples, then thread the array samples using array mode. For each sequencing site not genotyped by the array samples, we consider the marginal tree at that position in the inferred ARG. We select the branches in the marginal tree for which an unseen mutation best explains the observed sequencing data in terms of Hamming distance. Each of these branches implies genotypes of 0 or 1 for the array samples, so we output a weighted average of the implied genotypes, weighting branches by their length in the marginal tree. We applied this framework to perform genotype imputation using ground-truth ARGs and ARG-Needle inferred ARGs in 10 Mb of simulated data and compared to IMPUTE4 [40] and Beagle 5.1 [64] imputation using the binned aggregate *r*^2^ metric [39] (Supplementary Fig. S12b-c).

### ARG-Needle inference in the UK Biobank

Starting from 488,337 samples and 784,256 available autosomal array variants (including SNPs and short indels), we removed 135 samples (129 withdrawn, 6 due to missingness > 10%) and 57,126 variants (missingness > 10%). We phased the remaining variants and samples using Beagle 5.1 [65] and extracted the subset of 337,464 unrelated White British samples reported in [40]. We built the ARG of these samples using ARG-Needle, using parameters described above. We parallelized the ARG inference by splitting phased genotypes into 749 non-overlapping “chunks” of approximately equal numbers of variants. We added 50 variants on either side of each chunk to provide additional context for inference and independently applied ARG normalization on each chunk.

### Computation of permutation-based significance thresholds

We used a permutation-based approach to establish genome-wide significance thresholds corresponding to a family-wise error rate of 0.05 (Supplementary Table 1). In detail, we ran 1,000 null simulations using random phenotypes drawn from a standard normal distribution, performed univariate linear regression against imputed or ARG data [66, 13, 41], and computed the minimum *p*-value. We then estimated the genome-wide significance threshold using the most significant 5% quantile of these minimum *p*-values. We verified that performing this analysis using either the entire genome, chromosome 1, or the first chunk of the genome produced compatible results in a limited number of settings. To reduce computational costs, we thus performed these analyses using a subset of the genome and extrapolated to the whole genome. When a MAF cut-off was applied, significance thresholds remained compatible across several choices of sample size. We thus used 50, 000 haploid samples to estimate significance thresholds when MAF filtering was applied. Significance thresholds for several choices of filtering parameters are reported in Supplementary Table 1; specific thresholds used in individual analyses are detailed below.

### Genealogy-wide association scan in the UK Biobank

To process phenotypes (standing height, alkaline phosphatase, aspartate aminotransferase, LDL/HDL cholesterol, mean platelet volume, and total bilirubin) we first stratified by sex and performed quantile normalization. We then regressed out age, age squared, genotyping array, assessment center, and the first 20 genetic principal components computed in [40]. We built a non-infinitesimal BOLT-LMM mixed model using SNP array variants, then tested HRC+UK10K imputed data [38, 39, 40] and variants inferred using the ARG (ARG-MLMA, described above). For association of imputed data (including SNP array) we restricted to variants with Hardy-Weinberg equilibrium *p* > 10^−12^, missingness < 0.05, and info score > 0.3 (matching the filtering criteria adopted in [40]). For ARG-MLMA we tested variants sampled at rate *μ* = 10^−5^ and plotted variants sampled at *μ* = 10^−3^ in some follow-up analyses. For all analyses we did not test variants with a minor allele count (MAC) < 5 and used minor allele frequency (MAF) thresholds detailed below.

### Rare and ultra-rare variant association analysis

Using filtering criteria above and no additional MAF cutoff, we obtained genome-wide permutation significance thresholds of *p* < 4.8 × 10^−11^ (95% CI: [4.06 × 10^−11^, 5.99 × 10^−11^]) for ARG and *p* < 1.06 × 10^−9^ (95% CI: 5.13 × 10^−10^, 2.08 × 10^−9^]) for imputation. After performing genome-wide MLMA for the 7 traits, we selected genomic regions harboring rare (0.01% ≤ MAF < 0.1%) or ultra-rare (MAF < 0.01%) variant associations. We then formed regions by grouping any associated variants within 2 Mb of each other and adding 1 Mb on either side of the leftmost and rightmost variant in each of these groups. We next performed several filtering and association analyses to extract sets of approximately independent signals, using a procedure similar to that of [42], using PLINK (v1.90b6.21) with specified flags for several of these steps. For each region, we extracted hard-called raw genotypes for all genome-wide significant signals from either ARG or imputed data, tested for association (`-–assoc` flag), and performed two-stage LD-clumping of the variants. The first clumping step used parameters `--clump-p1 0.0001 --clump-p2 0.0001` `--clump-r2 0.5 --clump-kb 10`; the second used same parameters except for `--clump-kb 100000`. For each variant *i*, we considered each other variant *j* and computed the approximate chi-squared statistic that would be obtained by including *j* as covariate [44, 42]:

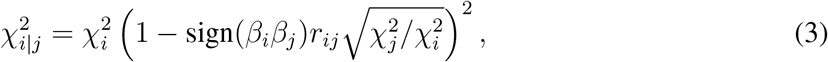

where 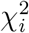 and 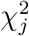 denote the respective chi-square statistics and sign(*β*_*i*_*β*_*j*_) is 1 if the effect sizes for the two variants have the same sign, -1 otherwise. LD was computed using the `--r` flag. We only retained variants *i* such that 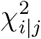 remained significant for all choices of *j*. We refer to the set of variants remaining after these filtering steps as approximately independent (“independent” for short, reported in Supplementary Tables 2-3). Of the 7 phenotypes, total bilirubin did not yield any rare or ultra-rare independent signals and height did not yield any independent ultra-rare signals.

We next leveraged the UK Biobank whole exome sequencing (WES) data to validate and localize independent associations. We extracted 138,039 exome sequenced samples that overlap with the analyzed set of White British individuals and performed lift-over of exome sequencing positions from genome build hg38 to hg19. We then computed pairwise LD (`--r` flag) between the set of independent associated variants and the set of all WES variants. The “WES partner” of an independent variant was selected to be the WES variant with largest *r*^2^ to it. For each pair of independent ARG or imputation signals with their WES partners, we computed the distance to the WES partner, the LD with the WES partner, the values of association 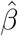 (with standard error), and the confusion matrix of genotype overlap between the independent signal and the WES partner, which we used to determine precision and recall of predicting the carriers in the WES variant (reported in Supplementary Tables 2 and 3). Because the BOLT-LMM non-infinitesimal model does not produce 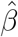 estimates, we instead obtained them using PLINK (`-–assoc` flag).

Variant annotations for the WES partners used in Fig. 4c were obtained using the Ensembl Variant Effect Predictor (VEP) tool [67]; variants reported by VEP as either of `frameshift_variant`, `splice_acceptor_variant`, `splice_donor_variant`, `stop_gained` were classed as loss-of-function (LoF) and variants reported as either of `stop_lost`, `start_lost`, `missense_variant`, `inframe_deletion`, `inframe_insertion`were classed as missense. Gene annotations for each WES variant were also obtained from VEP, where we allow for each WES partner to be annotated with multiple genes if they overlap the WES variant position. We used the WES variant position to check against the results of [42] to assess whether the variants we detected were present in the summary statistics, marginally significant (*p* < 5 × 10^−8^), or reported as likely causal (defined in [42]).

### Association analysis for higher frequency variants

For genome-wide association analyses of higher frequency variants and height, we matched filtering criteria used in [40], retaining imputed variants that satisfy the basic filters listed above, as well as MAF ≥ 0.1%. Using these criteria, we estimated a permutation-based genome-wide significance threshold of 4.5×10^−9^ (95% CI: [2.2×10^−9^, 9.6×10^−9^], Supplementary Table 1). To facilitate direct comparison, we aimed to select parameters that would result in a comparable significance threshold for the ARG-MLMA analysis. Two sets of parameters satisfied this requirement: 3.4 × 10^−9^ (95% CI: [2.4 × 10^−9^, 5 × 10^−9^]), obtained for *μ* = 10^−5^, MAF ≥ 1%; and 4 × 10^−9^ (95% CI: [3.1 × 10^−9^, 5.3 × 10^−9^]), obtained for *μ* = 10^−6^, MAF ≥ 0.1%. We selected the former set of parameters, as a low sampling rate of *μ* = 10^−6^ leads to worse signal-to-noise and lower association power. We thus used a significance threshold of *p* < 3 × 10^−9^ for all analyses of higher frequency variants.

To perform COJO analyses (Fig. 5c-d, Supplementary Fig. S9e) we first performed LD clumping of associated variants using PLINK (v1.90b6.21) with flags `--clump-p1 0.0001 --clump-p2 0.0001 --clump-r2 0.5 --clump-kb 1000` for all data types. For ARG data we also pre-processed each of the 749 non-overlapping chunks by running LD-clumping with the same parameters but with a reduced `--clump-kb 100`. The GCTA software implementing the COJO procedure requires effect size estimates, which are not produced by the BOLT-LMM non-infinitesimal model. We therefore extracted genotype variants with MLMA *p* < 5 × 10^−7^ for array data and *p* < 5 × 10^−8^ for imputed and ARG data and performed further association analysis using PLINK (v2.00a3LM, `--glm` flag). To create merged data sets (e.g. ARG + SNP array data), we started from the variants that were selected from the LD clumping step and merged them using PLINK. Because imputed data contains genotype array markers, we first removed any overlapping markers from the set of LD-clumped imputed variants in cases where both imputed and array data were merged and separately considered. COJO analyses were performed using GCTA (v1.93.2) using the `--cojo-slct` flag and COJO *p* < 3 × 10^−9^.

## Acknowledgements

We thank Po-Ru Loh, Alexander Gusev, Simon R. Myers, Robert Davies, Nicola Whiffin, Andrew Dahl, Árni Gunnarsson, and Romain Fournier for helpful discussions and suggestions; Fergus Cooper, Sinan Shi, Juba Nait Saada, and Georgios Kalantzis for sharing code used for various parts of the analysis. This work was conducted using the UK Biobank resource (Application #43206). We thank the participants of the UK Biobank project. This work was supported by the Clarendon Scholarship (to B.C.Z.); NIH grant R21-HG010748-01 (to P.F.P. and A.B.); Wellcome Trust ISSF grant 204826/Z/16/Z (to P.F.P.); and ERC Starting Grant ARGPHENO 850869 (to P.F.P., A.B., and B.C.Z). Computation used the Oxford Biomedical Research Computing (BMRC) facility, a joint development between the Wellcome Centre for Human Genetics and the Big Data Institute supported by Health Data Research UK and the NIHR Oxford Biomedical Research Centre. Financial support was provided by the Wellcome Trust Core Award Grant Number 203141/Z/16/Z. The views expressed are those of the author(s) and not necessarily those of the NHS, the NIHR or the Department of Health.

## Competing Interests

The authors declare no competing interests.

## Data Availability

VEP annotations were generated using the Ensembl VEP tool (v101.0, output produced February 2021), https://www.ensembl.org/info/docs/tools/vep/index.html. Other datasets were downloaded from the following URLs: summary statistics from whole exome imputation from 50K sequences [42], https://data.broadinstitute.org/lohlab/UKB_exomeWAS/; likely causal associations from whole exome imputation from 50K sequences [42], https://www.nature.com/ articles/s41588-021-00892-1 Supplementary Table 3; GIANT consortium summary statistics in ∼700K [43], https://portals.broadinstitute.org/collaboration/giant/index.php/GIANT_consortium_data_files.

## Code Availability

The software package ARG-Needle will be made freely available for academic use prior to publication at https://palamaralab.github.io/software/. The ARG-Needle package implements the ARG-Needle and ASMC-clust ARG inference methods as well as various analysis methods using inferred ARGs. External software used in the current study were obtained from the following URLs: msprime (v0.7.4), https://pypi.org/project/msprime/; tsinfer (v0.1.4), https://pypi.org/project/tsinfer/; Relate (v1.0.15), https://myersgroup.github.io/relate/; IMPUTE4 (v4.1.2), https://jmarchini.org/software/#impute-4; Beagle (v5.1), https://faculty.washington.edu/browning/beagle/b5_1.html; PLINK (v1.90b6.21), https://www.cog-genomics.org/plink/; PLINK (v2.00a3LM), https://www.cog-genomics.org/plink/2.0/; GCTA (v1.93.2), https://cnsgenomics.com/software/gcta/; BOLT-LMM (v2.3.2), https://alkesgroup.broadinstitute.org/BOLT-LMM/downloads/; LiftOver (used April 2021), https://genome.ucsc.edu/cgi-bin/hgLiftOver.

## Notes

### Competing Interest Statement

The authors have declared no competing interest.

## References (Main Text and Methods)

[1] Michael Bamshad and Stephen P Wooding. Signatures of natural selection in the human genome. Nature Reviews Genetics, 4(2):99–110, 2003.

[2] Annabel C Beichman, Emilia Huerta-Sanchez, and Kirk E Lohmueller. Using genomic data to infer historic population dynamics of nonmodel organisms. Annual Review of Ecology, Evolution, and Systematics, 2018.

[3] Sharon R Browning and Brian L Browning. Haplotype phasing: existing methods and new developments. Nature Reviews Genetics, 12(10):703–714, 2011.

[4] Jonathan Marchini and Bryan Howie. Genotype imputation for genome-wide association studies. Nature Reviews Genetics, 11(7):499–511, 2010.

[5] Gilean AT McVean and Niall J Cardin. Approximating the coalescent with recombination. Philosophical Transactions of the Royal Society B: Biological Sciences, 360(1459):1387–1393, 2005.

[6] Heng Li and Richard Durbin. Inference of human population history from individual whole-genome sequences. Nature, 475(7357):493–496, 2011.

[7] Sara Sheehan, Kelley Harris, and Yun S Song. Estimating variable effective population sizes from multiple genomes: a sequentially Markov conditional sampling distribution approach. Genetics, 194(3):647–662, 2013.

[8] Stephan Schiffels and Richard Durbin. Inferring human population size and separation history from multiple genome sequences. Nature Genetics, 46(8):919–925, 2014.

[9] Matthew D Rasmussen, Melissa J Hubisz, Ilan Gronau, and Adam Siepel. Genome-wide inference of ancestral recombination graphs. PLoS Genetics, 10(5):e1004342, 2014.

[10] Jonathan Terhorst, John A Kamm, and Yun S Song. Robust and scalable inference of population history from hundreds of unphased whole genomes. Nature Genetics, 49(2):303–309, 2017.

[11] Pier Francesco Palamara, Jonathan Terhorst, Yun S Song, and Alkes L Price. High-throughput inference of pairwise coalescence times identifies signals of selection and enriched disease heritability. Nature Genetics, 50(9):1311–1317, 2018.

[12] Rune B Lyngsø, Yun S Song, and Jotun Hein. Minimum recombination histories by branch and bound. In International Workshop on Algorithms in Bioinformatics, pages 239–250. Springer, 2005.

[13] Mark J Minichiello and Richard Durbin. Mapping trait loci by use of inferred ancestral recombination graphs. The American Journal of Human Genetics, 79(5):910–922, 2006.

[14] Sajad Mirzaei and Yufeng Wu. RENT+: an improved method for inferring local genealogical trees from haplotypes with recombination. Bioinformatics, 33(7):1021–1030, 2017.

[15] Jerome Kelleher, Yan Wong, Anthony W Wohns, Chaimaa Fadil, Patrick K Albers, and Gil McVean. Inferring whole-genome histories in large population datasets. Nature Genetics, 51(9):1330–1338, 2019.

[16] Nathan K Schaefer, Beth Shapiro, and Richard E Green. An ancestral recombination graph of human, Neanderthal, and Denisovan genomes. Science Advances, 7(29):eabc0776, 2021.

[17] Leo Speidel, Marie Forest, Sinan Shi, and Simon R Myers. A method for genome-wide genealogy estimation for thousands of samples. Nature Genetics, 51(9):1321–1329, 2019.

[18] Leo Speidel, Lara Cassidy, Robert W Davies, Garrett Hellenthal, Pontus Skoglund, and Simon R Myers. Inferring population histories for ancient genomes using genome-wide genealogies. Molecular Biology and Evolution, 2021.

[19] Sebastian Zöllner and Jonathan K Pritchard. Coalescent-based association mapping and fine mapping of complex trait loci. Genetics, 169(2):1071–1092, 2005.

[20] Hyun Min Kang, Noah A Zaitlen, Claire M Wade, Andrew Kirby, David Heckerman, Mark J Daly, and Eleazar Eskin. Efficient control of population structure in model organism association mapping. Genetics, 178(3):1709–1723, 2008.

[21] Jian Yang, S Hong Lee, Michael E Goddard, and Peter M Visscher. GCTA: a tool for genome-wide complex trait analysis. The American Journal of Human Genetics, 88(1):76–82, 2011.

[22] Po-Ru Loh, Gleb Kichaev, Steven Gazal, Armin P Schoech, and Alkes L Price. Mixed-model association for biobank-scale datasets. Nature Genetics, 50(7):906–908, 2018.

[23] Robert C Griffiths and Paul Marjoram. An ancestral recombination graph. Institute for Mathematics and its Applications, 87:257, 1997.

[24] Alexander Gusev, Jennifer K Lowe, Markus Stoffel, Mark J Daly, David Altshuler, Jan L Breslow, Jeffrey M Friedman, and Itsik Pe’er. Whole population, genome-wide mapping of hidden relatedness. Genome Research, 19(2):318–326, 2009.

[25] Juba Nait Saada, Georgios Kalantzis, Derek Shyr, Fergus Cooper, Martin Robinson, Alexander Gusev, and Pier Francesco Palamara. Identity-by-descent detection across 487,409 British samples reveals fine scale population structure and ultra-rare variant associations. Nature Communications, 11(1):1–15, 2020.

[26] David F Robinson and Leslie R Foulds. Comparison of phylogenetic trees. Mathematical Biosciences, 53(1-2):131–147, 1981.

[27] Michelle Kendall and Caroline Colijn. Mapping phylogenetic trees to reveal distinct patterns of evolution. Molecular Biology and Evolution, 33(10):2735–2743, 2016.

[28] Jian Yang, Beben Benyamin, Brian P McEvoy, Scott Gordon, Anjali K Henders, Dale R Nyholt, Pamela A Madden, Andrew C Heath, Nicholas G Martin, Grant W Montgomery, et al. Common SNPs explain a large proportion of the heritability for human height. Nature Genetics, 42(7):565, 2010.

[29] Luke M Evans, Rasool Tahmasbi, Scott I Vrieze, Gonçalo R Abecasis, Sayantan Das, Steven Gazal, Douglas W Bjelland, Teresa R De Candia, Michael E Goddard, Benjamin M Neale, et al. Comparison of methods that use whole genome data to estimate the heritability and genetic architecture of complex traits. Nature Genetics, 50(5):737, 2018.

[30] Naomi R Wray, Jian Yang, Ben J Hayes, Alkes L Price, Michael E Goddard, and Peter M Visscher. Pitfalls of predicting complex traits from SNPs. Nature Reviews Genetics, 14(7):507–515, 2013.

[31] Jian Yang, Noah A Zaitlen, Michael E Goddard, Peter M Visscher, and Alkes L Price. Advantages and pitfalls in the application of mixed-model association methods. Nature Genetics, 46(2):100, 2014.

[32] Roderick HJ Houwen, Siamak Baharloo, Kathleen Blankenship, Peter Raeymaekers, Jenneke Juyn, Lodewijk A Sandkuijl, and Nelson B Freimer. Genome screening by searching for shared segments: mapping a gene for benign recurrent intrahepatic cholestasis. Nature Genetics, 8(4):380–386, 1994.

[33] Alexander Gusev, Eimear E Kenny, Jennifer K Lowe, Jaqueline Salit, Richa Saxena, Sekar Kathiresan, David M Altshuler, Jeffrey M Friedman, Jan L Breslow, and Itsik Pe’er. DASH: a method for identical-by-descent haplotype mapping uncovers association with recent variation. The American Journal of Human Genetics, 88(6):706–717, 2011.

[34] Sharon R Browning and Elizabeth A Thompson. Detecting rare variant associations by identity-by-descent mapping in case-control studies. Genetics, 190(4):1521–1531, 2012.

[35] Jian Yang, Andrew Bakshi, Zhihong Zhu, Gibran Hemani, Anna AE Vinkhuyzen, Sang Hong Lee, Matthew R Robinson, John RB Perry, Ilja M Nolte, Jana V van Vliet-Ostaptchouk, et al. Genetic variance estimation with imputed variants finds negligible missing heritability for human height and body mass index. Nature Genetics, 47(10):1114, 2015.

[36] Pierrick Wainschtein, Deepti Jain, Zhili Zheng, L Adrienne Cupples, Aladdin H Shadyab, Barbara McKnight, Benjamin M Shoemaker, Braxton D Mitchell, Bruce M Psaty, Charles Kooperberg, et al. Recovery of trait heritability from whole genome sequence data. bioRxiv, page 588020, 2021.

[37] Po-Ru Loh, George Tucker, Brendan K Bulik-Sullivan, Bjarni J Vilhjalmsson, Hilary K Finucane, Rany M Salem, Daniel I Chasman, Paul M Ridker, Benjamin M Neale, Bonnie Berger, et al. Efficient Bayesian mixed-model analysis increases association power in large cohorts. Nature Genetics, 47(3):284, 2015.

[38] Jie Huang, Bryan Howie, Shane McCarthy, Yasin Memari, Klaudia Walter, Josine L Min, Petr Danecek, Giovanni Malerba, Elisabetta Trabetti, Hou-Feng Zheng, et al. Improved imputation of low-frequency and rare variants using the UK10K haplotype reference panel. Nature Communications, 6(1):1–9, 2015.

[39] Shane McCarthy, Sayantan Das, Warren Kretzschmar, Olivier Delaneau, Andrew R Wood, Alexander Teumer, Hyun Min Kang, Christian Fuchsberger, Petr Danecek, Kevin Sharp, et al. A reference panel of 64,976 haplotypes for genotype imputation. Nature Genetics, 48(10):1279, 2016.

[40] Clare Bycroft, Colin Freeman, Desislava Petkova, Gavin Band, Lloyd T Elliott, Kevin Sharp, Allan Motyer, Damjan Vukcevic, Olivier Delaneau, Jared O’Connell, et al. The UK Biobank resource with deep phenotyping and genomic data. Nature, 562(7726):203–209, 2018.

[41] Masahiro Kanai, Toshihiro Tanaka, and Yukinori Okada. Empirical estimation of genome-wide significance thresholds based on the 1000 Genomes Project data set. Journal of Human Genetics, 61(10):861–866, 2016.

[42] Alison R Barton, Maxwell A Sherman, Ronen E Mukamel, and Po-Ru Loh. Whole-exome imputation within UK Biobank powers rare coding variant association and fine-mapping analyses. Nature Genetics, 53(8):1260–1269, 2021.

[43] Loic Yengo, Julia Sidorenko, Kathryn E Kemper, Zhili Zheng, Andrew R Wood, Michael N Weedon, Timothy M Frayling, Joel Hirschhorn, Jian Yang, Peter M Visscher, et al. Meta-analysis of genome-wide association studies for height and body mass index in ⇠700000 individuals of European ancestry. Human Molecular Genetics, 27(20):3641–3649, 2018.

[44] Jian Yang, Teresa Ferreira, Andrew P Morris, Sarah E Medland, Pamela AF Madden, Andrew C Heath, Nicholas G Martin, Grant W Montgomery, Michael N Weedon, Ruth J Loos, et al. Conditional and joint multiple-SNP analysis of GWAS summary statistics identifies additional variants influencing complex traits. Nature Genetics, 44(4):369–375, 2012.

[45] David E Reich, Michele Cargill, Stacey Bolk, James Ireland, Pardis C Sabeti, Daniel J Richter, Thomas Lavery, Rose Kouyoumjian, Shelli F Farhadian, Ryk Ward, et al. Linkage disequilibrium in the human genome. Nature, 411(6834):199–204, 2001.

[46] Daniel Taliun, Daniel N Harris, Michael D Kessler, Jedidiah Carlson, Zachary A Szpiech, Raul Torres, Sarah A Gagliano Taliun, André Corvelo, Stephanie M Gogarten, Hyun Min Kang, et al. Sequencing of 53,831 diverse genomes from the NHLBI TOPMed Program. Nature, 590(7845):290–299, 2021.

[47] Alicia R Martin, Masahiro Kanai, Yoichiro Kamatani, Yukinori Okada, Benjamin M Neale, and Mark J Daly. Clinical use of current polygenic risk scores may exacerbate health disparities. Nature Genetics, 51(4):584–591, 2019.

[48] Po-Ru Loh, Gaurav Bhatia, Alexander Gusev, Hilary K Finucane, Brendan K Bulik-Sullivan, Samuela J Pollack, Teresa R de Candia, Sang Hong Lee, Naomi R Wray, Kenneth S Kendler, et al. Contrasting genetic architectures of schizophrenia and other complex diseases using fast variance-components analysis. Nature Genetics, 47(12):1385, 2015.

[49] Ali Pazokitoroudi, Yue Wu, Kathryn S Burch, Kangcheng Hou, Aaron Zhou, Bogdan Pasaniuc, and Sriram Sankararaman. Efficient variance components analysis across millions of genomes. Nature Communications, 11(1):1–10, 2020.

[50] Steven Gazal, Carla Marquez-Luna, Hilary K Finucane, and Alkes L Price. Reconciling S-LDSC and LDAK functional enrichment estimates. Nature Genetics, 51(8):1202–1204, 2019.

[51] Alexander I Young, Michael L Frigge, Daniel F Gudbjartsson, Gudmar Thorleifsson, Gyda Bjornsdottir, Patrick Sulem, Gisli Masson, Unnur Thorsteinsdottir, Kari Stefansson, and Augustine Kong. Relatedness disequilibrium regression estimates heritability without environmental bias. Nature Genetics, 50(9):1304–1310, 2018.

[52] Yichen Si, Brett Vanderwerff, and Sebastian Zöllner. Why are rare variants hard to impute? Coalescent models reveal theoretical limits in existing algorithms. Genetics, 217(4):iyab011, 2021.

[53] Yoshiaki Yasumizu, Saori Sakaue, Takahiro Konuma, Ken Suzuki, Koichi Matsuda, Yoshinori Murakami, Michiaki Kubo, Pier Francesco Palamara, Yoichiro Kamatani, and Yukinori Okada. Genome-wide natural selection signatures are linked to genetic risk of modern phenotypes in the Japanese population. Molecular Biology and Evolution, 37(5):1306–1316, 2020.

[54] Aaron J Stern, Leo Speidel, Noah A Zaitlen, and Rasmus Nielsen. Disentangling selection on genetically correlated polygenic traits via whole-genome genealogies. The American Journal of Human Genetics, 108(2):219–239, 2021.

[55] Peter HA Sneath and Robert R Sokal. Numerical taxonomy. The principles and practice of numerical classification. W. H.. Freeman and Co., 1973.

[56] Ilan Gronau and Shlomo Moran. Optimal implementations of UPGMA and other common clustering algorithms. Information Processing Letters, 104(6):205–210, 2007.

[57] Daniel Müllner. fastcluster: Fast hierarchical, agglomerative clustering routines for R and Python. Journal of Statistical Software, 53(9):1–18, 2013.

[58] Jerome Kelleher, Alison M Etheridge, and Gilean McVean. Efficient coalescent simulation and genealogical analysis for large sample sizes. PLoS Computational Biology, 12(5):e1004842, 2016.

[59] Doug Speed, Gibran Hemani, Michael R Johnson, and David J Balding. Improved heritability estimation from genome-wide SNPs. The American Journal of Human Genetics, 91(6):1011–1021, 2012.

[60] S Hong Lee, Jian Yang, Guo-Bo Chen, Stephan Ripke, Eli A Stahl, Christina M Hultman, Pamela Sklar, Peter M Visscher, Patrick F Sullivan, Michael E Goddard, et al. Estimation of SNP heritability from dense genotype data. The American Journal of Human Genetics, 93(6):1151–1155, 2013.

[61] Steven Gazal, Hilary K Finucane, Nicholas A Furlotte, Po-Ru Loh, Pier Francesco Palamara, Xuanyao Liu, Armin Schoech, Brendan Bulik-Sullivan, Benjamin M Neale, Alexander Gusev, et al. Linkage disequilibrium–dependent architecture of human complex traits shows action of negative selection. Nature Genetics, 49(10):1421–1427, 2017.

[62] Hans D Daetwyler, Beatriz Villanueva, and John A Woolliams. Accuracy of predicting the genetic risk of disease using a genome-wide approach. PLoS ONE, 3(10):e3395, 2008.

[63] Joel Mefford, Danny Park, Zhili Zheng, Arthur Ko, Mika Ala-Korpela, Markku Laakso, Päivi Pajukanta, Jian Yang, John Witte, and Noah Zaitlen. Efficient estimation and applications of cross-validated genetic predictions to polygenic risk scores and linear mixed models. Journal of Computational Biology, 27(4):599–612, 2020.

[64] Brian L Browning, Ying Zhou, and Sharon R Browning. A one-penny imputed genome from next-generation reference panels. The American Journal of Human Genetics, 103(3):338–348, 2018.

[65] Sharon R Browning and Brian L Browning. Rapid and accurate haplotype phasing and missing-data inference for whole-genome association studies by use of localized haplotype clustering. The American Journal of Human Genetics, 81(5):1084–1097, 2007.

[66] Gary A Churchill and Rebecca W Doerge. Empirical threshold values for quantitative trait mapping. Genetics, 138(3):963–971, 1994.

[67] William McLaren, Laurent Gil, Sarah E Hunt, Harpreet Singh Riat, Graham RS Ritchie, Anja Thormann, Paul Flicek, and Fiona Cunningham. The Ensembl Variant Effect Predictor. Genome Biology, 17(1):1–14, 2016.

