## Supplementary Figures and Notes for "Biobank-scale inference of ancestral recombination graphs enables genealogy-based mixed model association of complex traits"

Zhang et al.

### Supplementary Figures

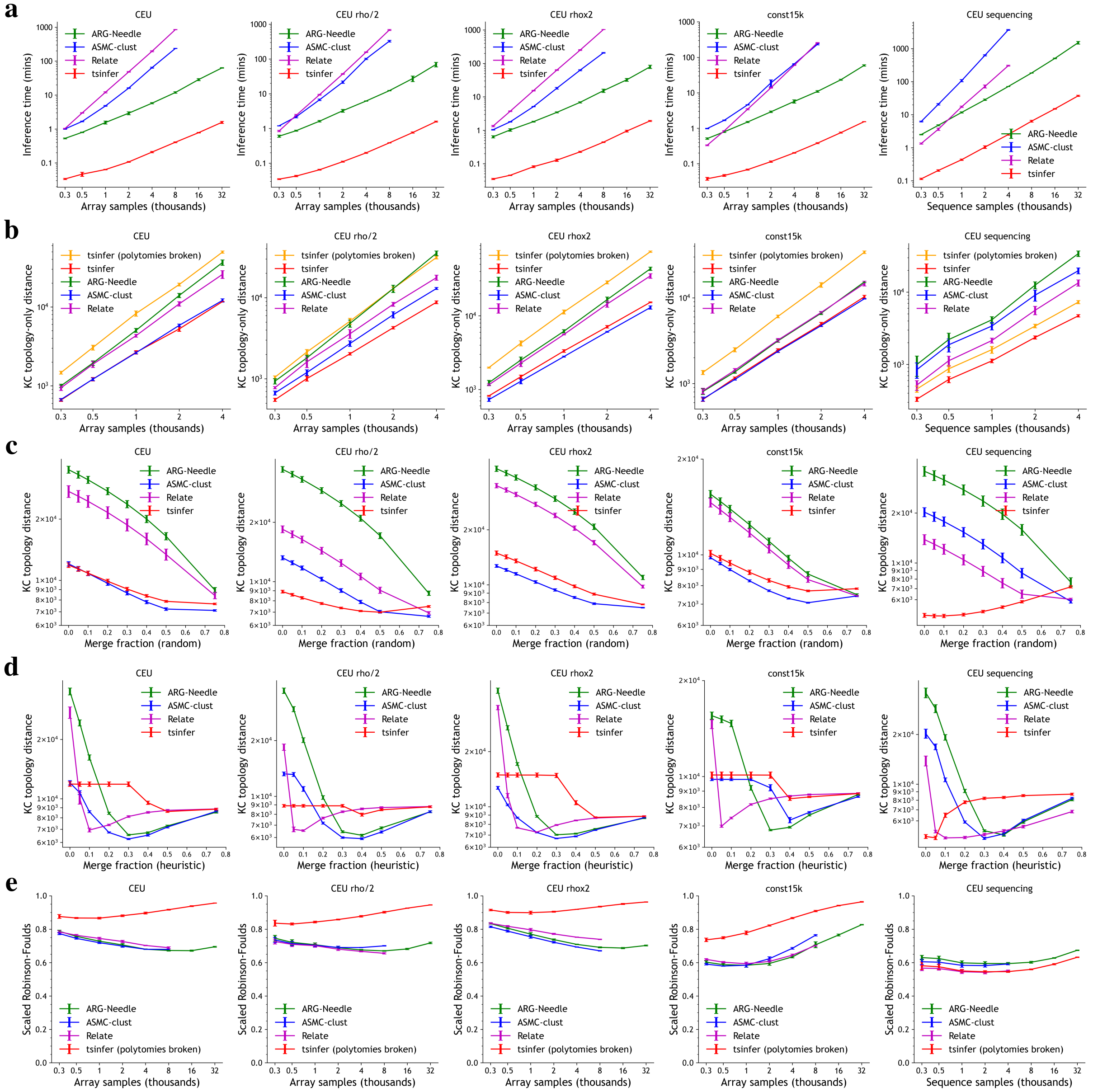

Figure S1: **Additional comparison of ARG inference methods (topology-only metrics).** We compare methods on runtime and topology-only metrics, as in Fig. 2 but with additional simulation conditions. Columns correspond to 5 Mb of CEU (Central European demography) array data with standard parameters (see Methods), 5 Mb of CEU array data with a factor of 2 smaller recombination rate ($\rho = 6 \times 10^{-9}$), 5 Mb of CEU array data with a factor of 2 larger recombination rate ($\rho = 2.4 \times 10^{-8}$), 5 Mb of array data under a constant population size demography of 15,000 individuals and otherwise standard parameters, and 1 Mb of CEU sequencing data with otherwise standard parameters. **a**. Inference time as a function of the number of samples $N$. **b**. KC topology-only distance as a function of $N$, additionally showing the results of tsinfer with randomly resolved polytomies. **c**. KC topology-only distance for $N = 4{,}000$ samples, showing performance as branches in marginal inferred trees are randomly collapsed to form polytomies. **d**. The same as **c**, except using a heuristic to preferentially merge branches that are least certain (see Supplementary Note 2). **e**. Robinson-Foulds distance as a function of $N$, where values are scaled to lie between 0 and 1, and with randomly resolved polytomies. Error bars represent 2 s.e.

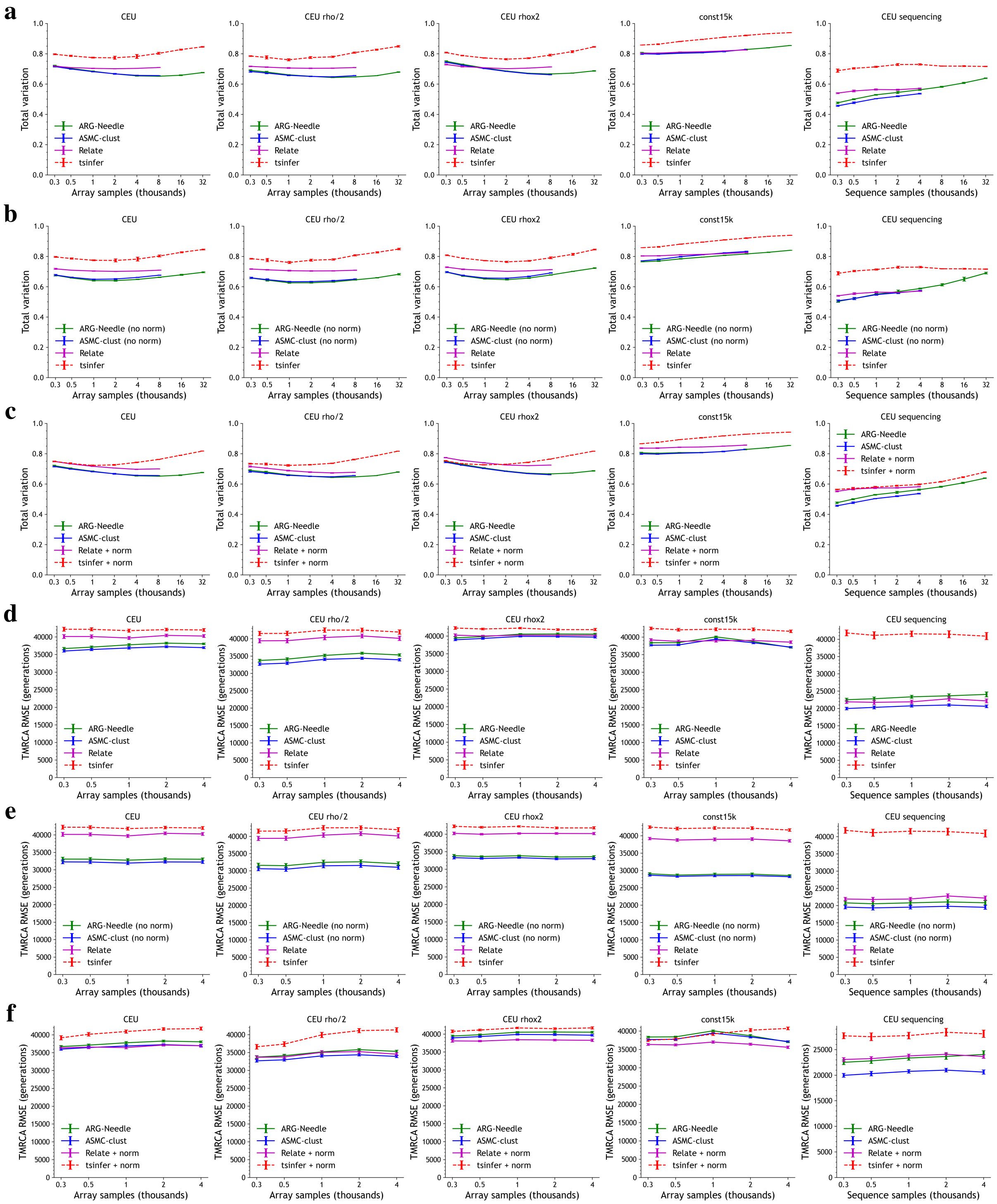

Figure S2: **Additional comparison of ARG inference methods for metrics that take into account branch length.** As in Fig. 2b-c but with additional simulation conditions. Columns correspond to 5 Mb of CEU array data with standard parameters (see Methods), 5 Mb of CEU array data with a factor of 2 smaller recombination rate ($\rho = 6 \times 10^{-9}$), 5 Mb of CEU array data with a factor of 2 larger recombination rate ($\rho = 2.4 \times 10^{-8}$), 5 Mb of array data under a demography with a constant population size of 15,000 diploids and otherwise standard parameters, and 1 Mb of CEU sequencing data with otherwise standard parameters. We show results for the ARG total variation distance (**a**-**c**) and pairwise TMRCA RMSE (**d**-**f**) across different conditions with and without ARG normalization for the various methods, as these metrics are sensitive to branch length. tsinfer is shown with dotted lines as it focuses on topologies. Error bars represent 2 s.e.

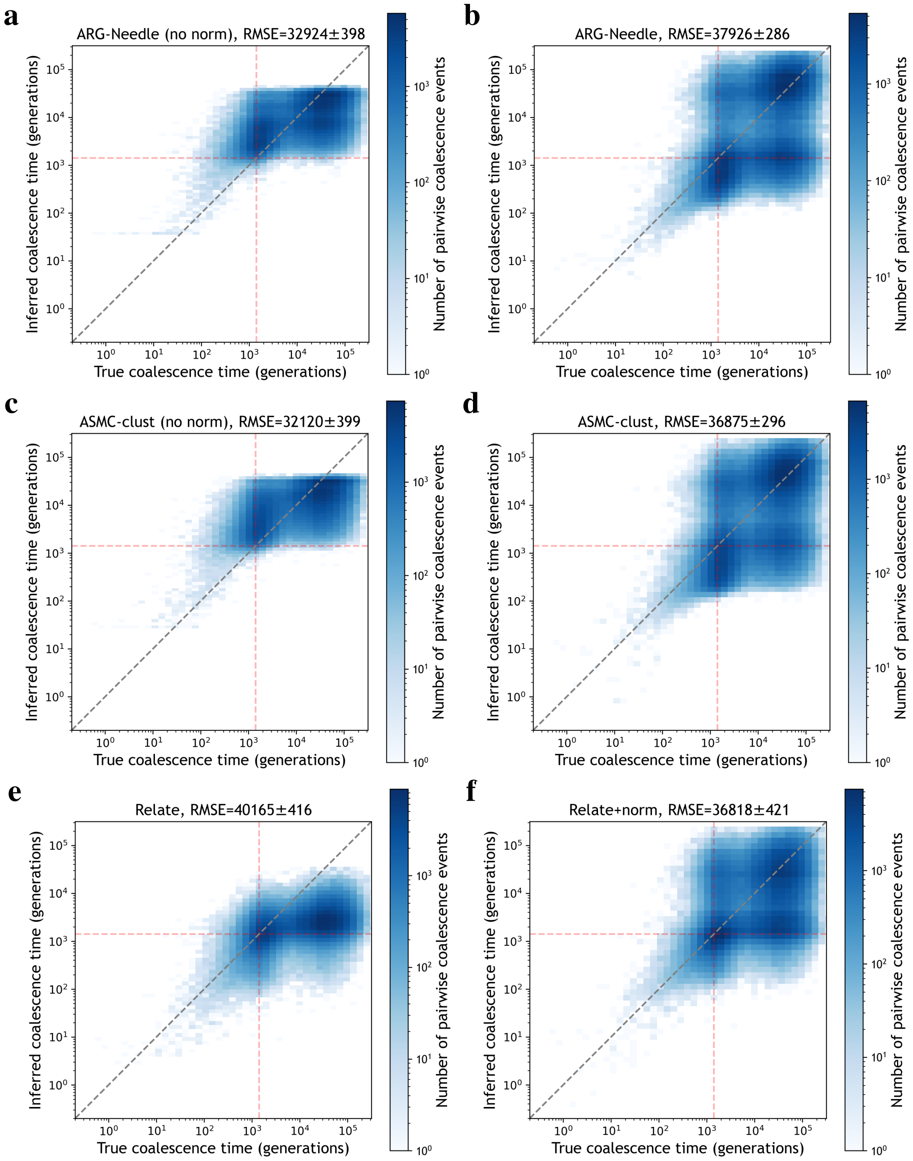

Figure S3: **Scatter plots of true vs. inferred pairwise TMRCAs.** We show scatter plots for ARG-Needle (**a**-**b**), ASMC-clust (**c**-**d**), and Relate (**e**-**f**), for $N = 4{,}000$ array samples, showing an aggregate over 20,000 randomly sampled pairs for each of 25 simulations. The left column corresponds to no ARG normalization and the right column corresponds to including ARG normalization. Titles display TMRCA RMSE with 2 s.e. from bootstrap sampling of the 25 simulations used. Removing ARG normalization decreases the pairwise TMRCA RMSE for ARG-Needle and ASMC-clust, but skews the distribution of pairwise TMRCAs towards the center. Dotted red lines show the time of the CEU demography population bottleneck.

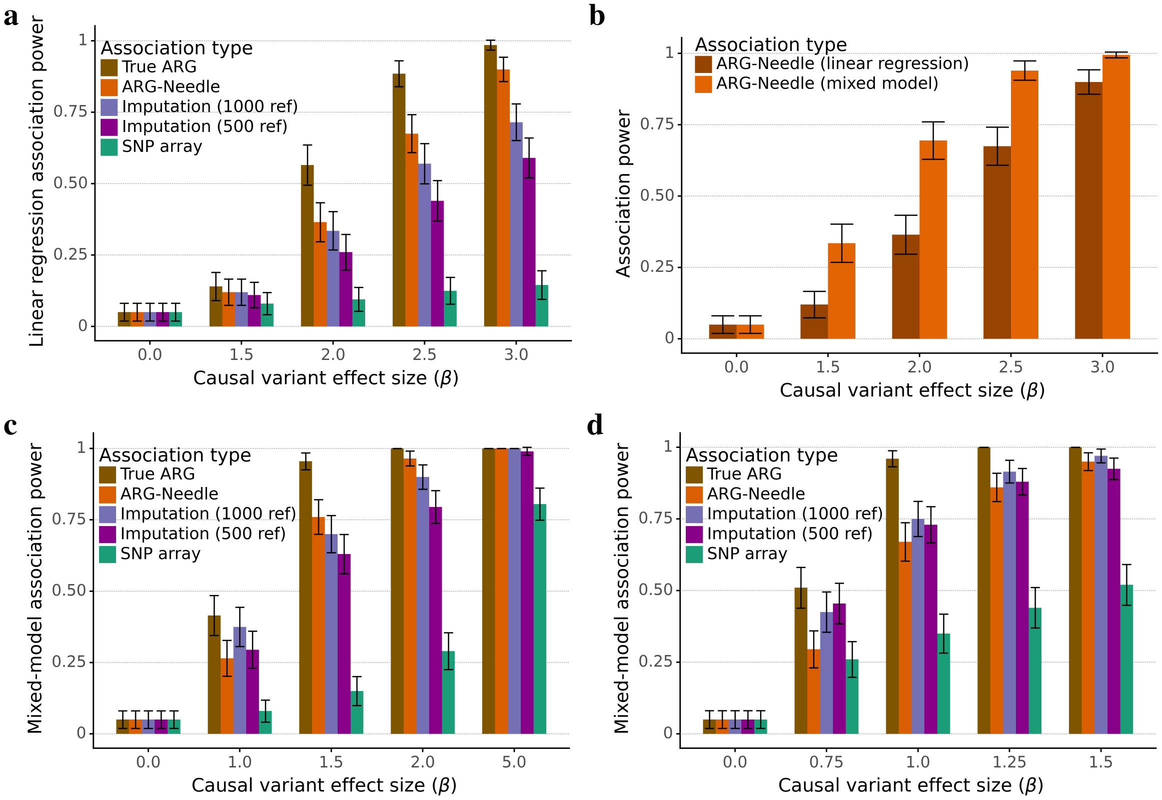

Figure S4: **Additional simulations of ARG-MLMA genealogy-wide association power. a**. Similar to Fig. 3a, except using linear regression instead of the linear mixed model to test for association. **b**. We combine the association power results of ARG-Needle association from **a** and Fig. 3a, highlighting the improvement of ARG-MLMA compared to directly testing ARG clades using linear regression. **c**,**d**. Similar to Fig. 3a, except with the causal variant MAF chosen to be 0.5% (**c**) and 1% (**d**) instead of 0.25%. *ref* indicates the number of haploid reference samples used for imputation.

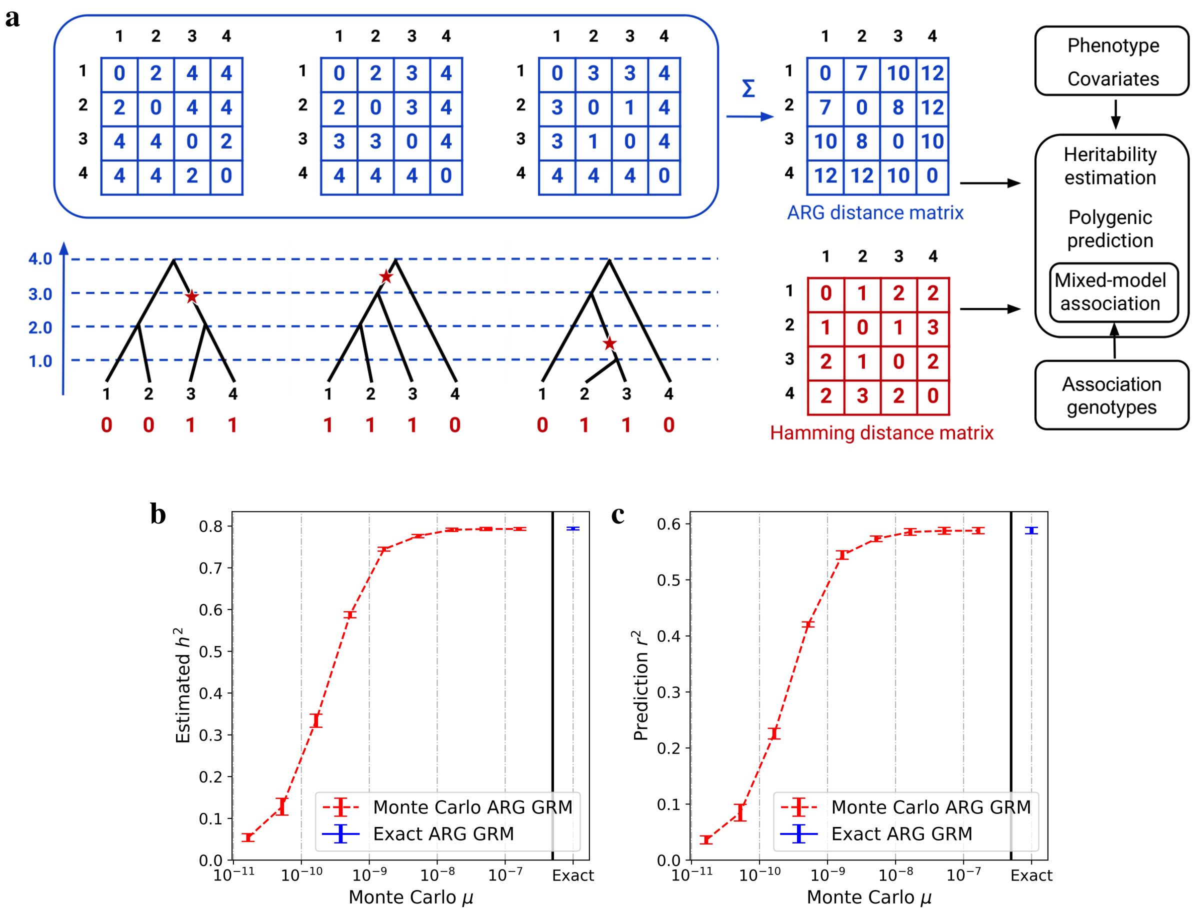

Figure S5: **Overview of ARG-GRM definition and Monte Carlo estimator. a**. Schematic of ARG-GRMs. Given an ARG between samples, we can compute the TMRCA matrix at each site and sum this over the genome to obtain the $\alpha = 0$ ARG distance matrix (top, in blue). This equals a scaled version of the expected Hamming distance matrix (bottom, in red), which is formed by counting the number of differences between the genotypes of samples. By applying a series of simple matrix transformations to the ARG distance matrix (see Supplementary Note 3), we obtain the ARG-GRM, which can subsequently be used in complex trait analysis just like genotype-based GRMs. **b**,**c**. We compare the use of an exact $\alpha = 0$ ARG-GRM to Monte Carlo $\alpha = 0$ ARG-GRMs for heritability estimation (**b**) and polygenic prediction (**c**). As we increase the mutation rate for the Monte Carlo ARG-GRMs (rightmost value of $\mu = 1.65 \times 10^{-7}$), we approach results from using the exact ARG-GRM. Simulations use $N = 2{,}000$ haploid samples, $h^2 = 0.8$, $\alpha = 0$, and 10 Mb. We show mean values over 5 runs. Error bars represent 2 s.e. from meta-analysis.

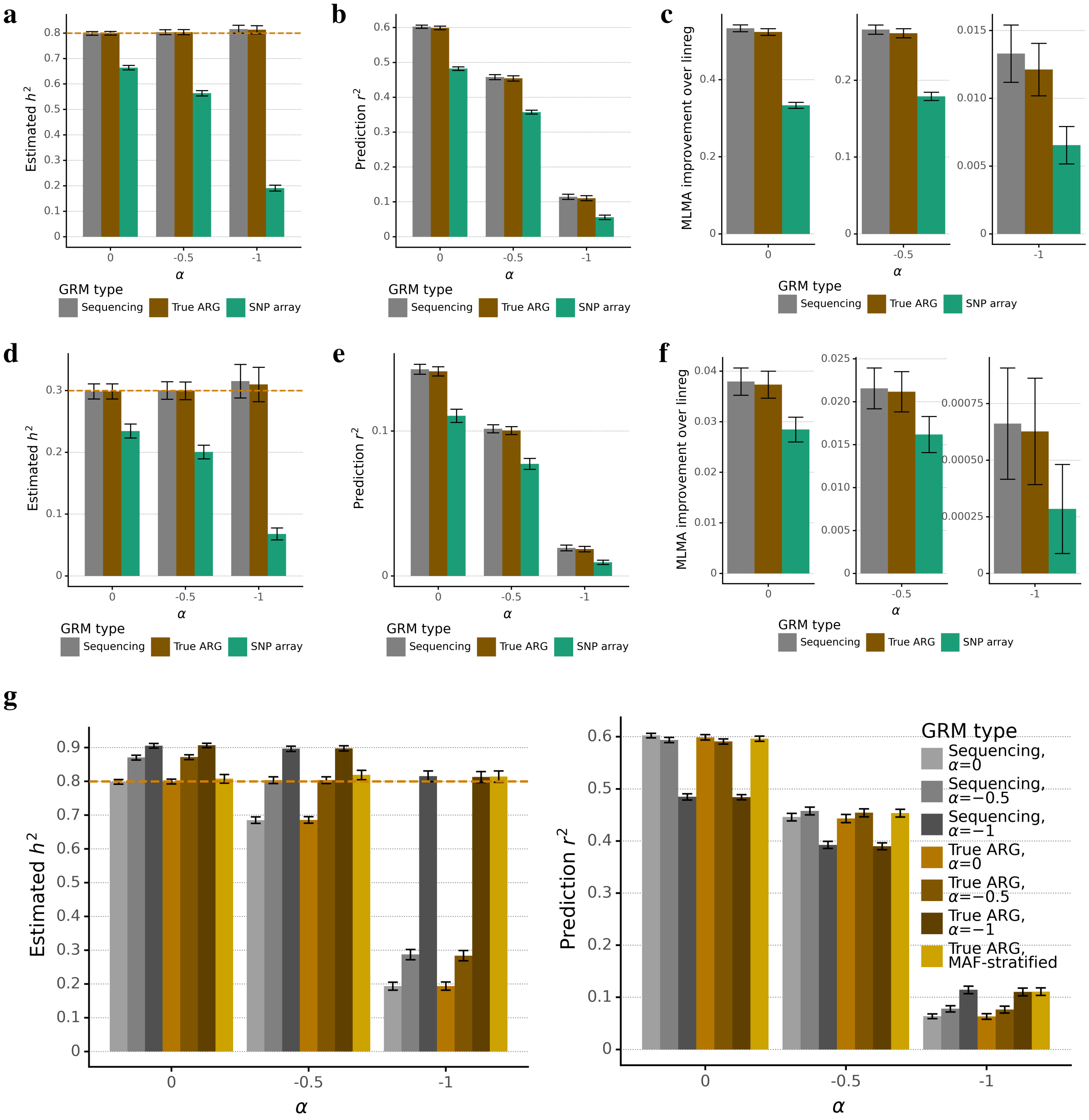

Figure S6: **Additional simulations for ground-truth ARG-GRMs.** **a**-**f**. As in Fig. 3c, with $N = 10{,}000$ haploid samples, except we vary $h^2 \in \{0.8, 0.3\}$ and $\alpha \in \{0, -0.5, -1\}$. **a**,**d**. Heritability estimation for a 50 Mb region for $h^2 = 0.8$ (**a**) and $h^2 = 0.3$ (**d**). **b**,**e**. Polygenic prediction for a 50 Mb region for $h^2 = 0.8$ (**b**) and $h^2 = 0.3$ (**e**). **c**,**f**. Mixed-model association for 22 chromosomes of 2.5 Mb each for $h^2 = 0.8$ (**c**) and $h^2 = 0.3$ (**f**). **g**. Panels **a**-**f** assumed it is possible to infer $\alpha$ and used the true $\alpha$ when building genotype-based or ARG-GRMs. If this value of $\alpha$ is misspecified, heritability estimation is biased and prediction $r^2$ is hampered. This is true both for ARG-GRMs and sequencing GRMs. However, using MAF-stratified ARG-GRMs provides a robust way to estimate the true heritability when $\alpha$ is unknown, and achieves prediction $r^2$ comparable to using the true $\alpha$ ($N = 10{,}000$ haploid samples, 50 Mb, $h^2 = 0.8$). For all panels, heritability and prediction experiments involve 5 simulations per bar, and most association experiments involve 50 simulations per bar, except for the $h^2 = 0.3, \alpha = -1$ condition in **f**, which involved 500 simulations. Error bars represent 2 s.e. (from meta-analysis in the case of heritability estimation).

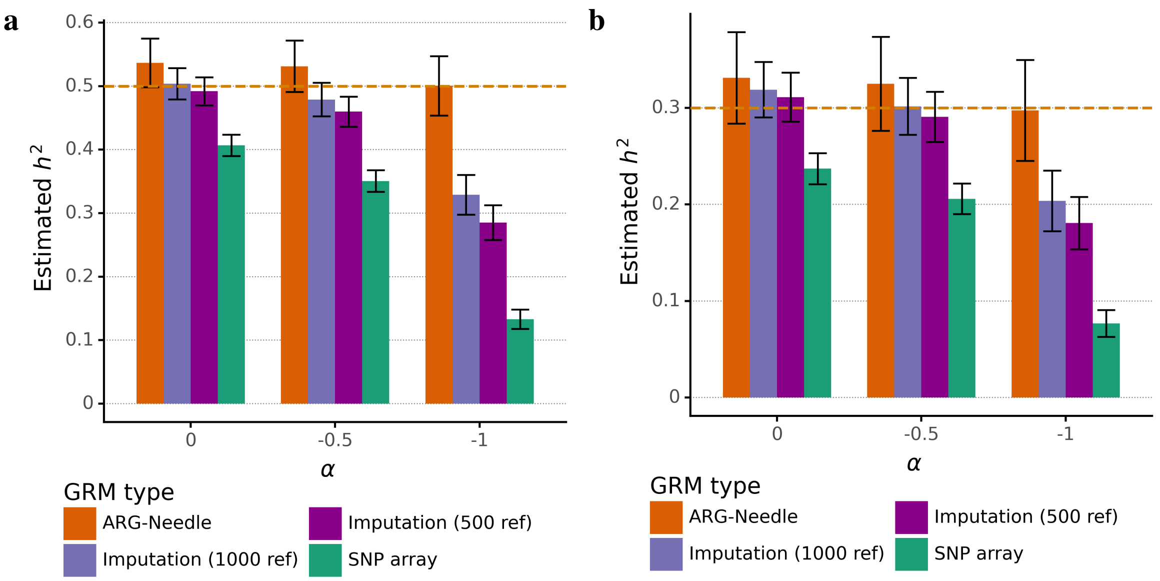

Figure S7: **Heritability estimation using ARG-Needle and ARG-GRMs.** Heritability estimation using ARG-GRMs with ARGs inferred from SNP data, compared to using GRMs of imputed data or array SNPs. As in Fig. 3b but with $h^2 = 0.5$ (**a**) and $h^2 = 0.3$ (**b**). Results are for 5 runs with $N = 5{,}000$ haploid samples, 25 Mb, and $\alpha \in \{0, -0.5, -1\}$. Error bars represent 2 s.e. from meta-analysis. *ref* indicates the number of haploid reference samples used for imputation.

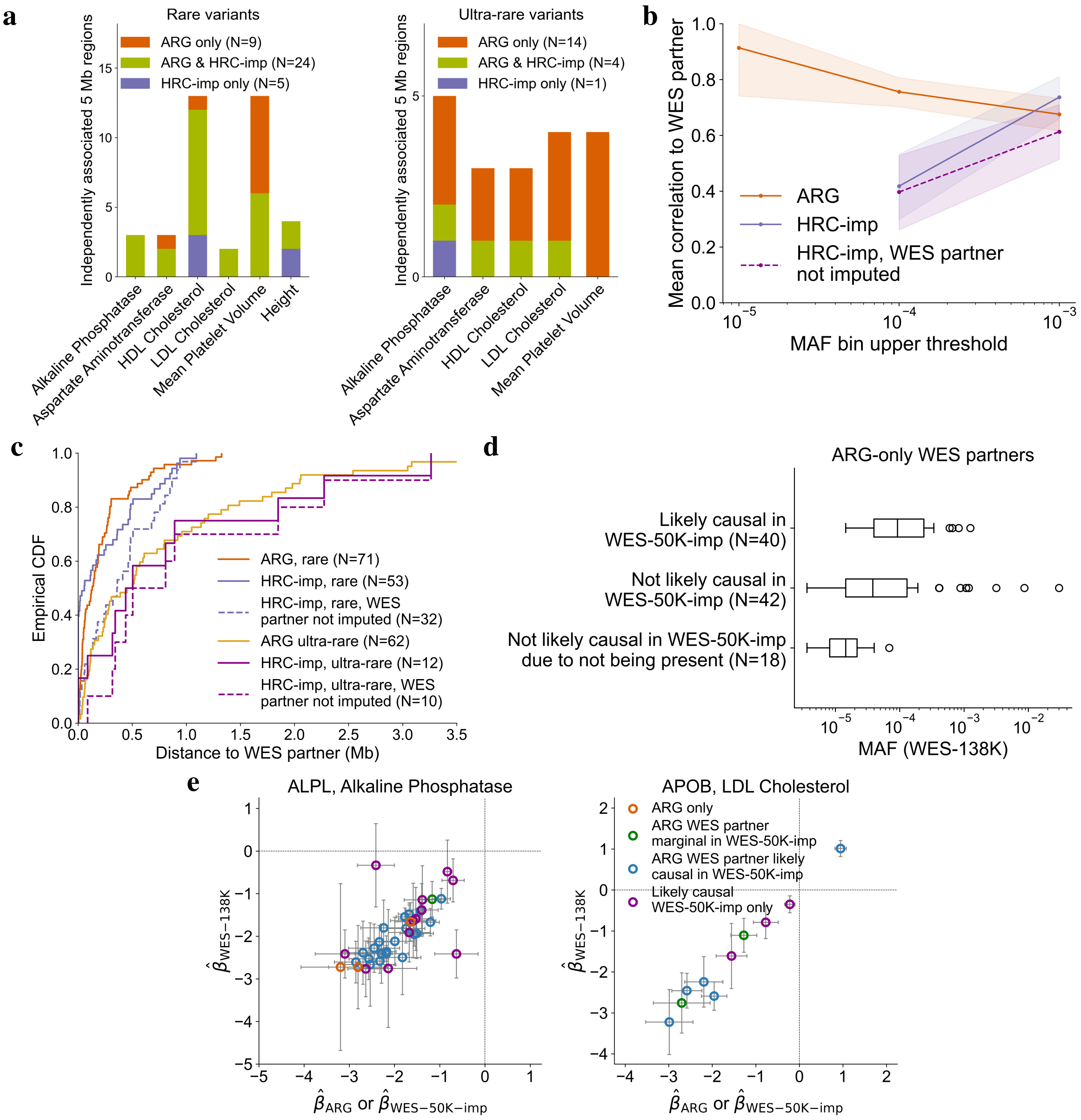

Figure S8: **Further results for rare and ultra-rare variant associations.** **a**. Counts of implicated 5 Mb regions containing ARG and HRC+UK10K imputation ("HRC-imp") independent associations, partitioned by traits and frequency and showing overlap. Total bilirubin was not associated at these frequencies. **b**. Average Pearson correlation between independent variants and their WES partners as a function of frequency, for ARG-derived variants, HRC+UK10K imputed variants, and HRC+UK10K imputed variants for which the WES partner was not the imputed variant. Dots represent the upper end of a frequency range. Shaded areas represent 95% bootstrap confidence intervals. **c**. Cumulative distribution function for the distance between independent variants and their WES partners, partitioned by frequency. As in Fig. 4b, but also showing HRC+UK10K imputed variants for which the WES partner was not the imputed variant. **d**. Box plots of MAF for WES partners found by ARG-derived but not HRC+UK10K imputed independent variants (center line, median; box limits, upper and lower quartiles, whiskers, 1.5x interquartile range; points, outliers), stratifying by status in WES-50K-imp (imputation from WES-50K). **e**. Scatter plot of $\hat{\beta}$ (estimated effect) for ARG-derived independent variants against $\hat{\beta}$ for their WES partners, as in Fig. 4f but for associations with alkaline phosphatase in the *ALPL* gene and with LDL cholesterol in the *APOB* gene. We color points based on whether the WES partner is likely causal in WES-50K-imp, not likely causal but marginally significant in WES-50K-imp, or not marginally significant in WES-50K-imp ("ARG only" in figure). We also plot the $\hat{\beta}$ for the additional likely causal variants in WES-50K-imp against the $\hat{\beta}$ in WES-138K. Error bars represent 1.96 s.e.

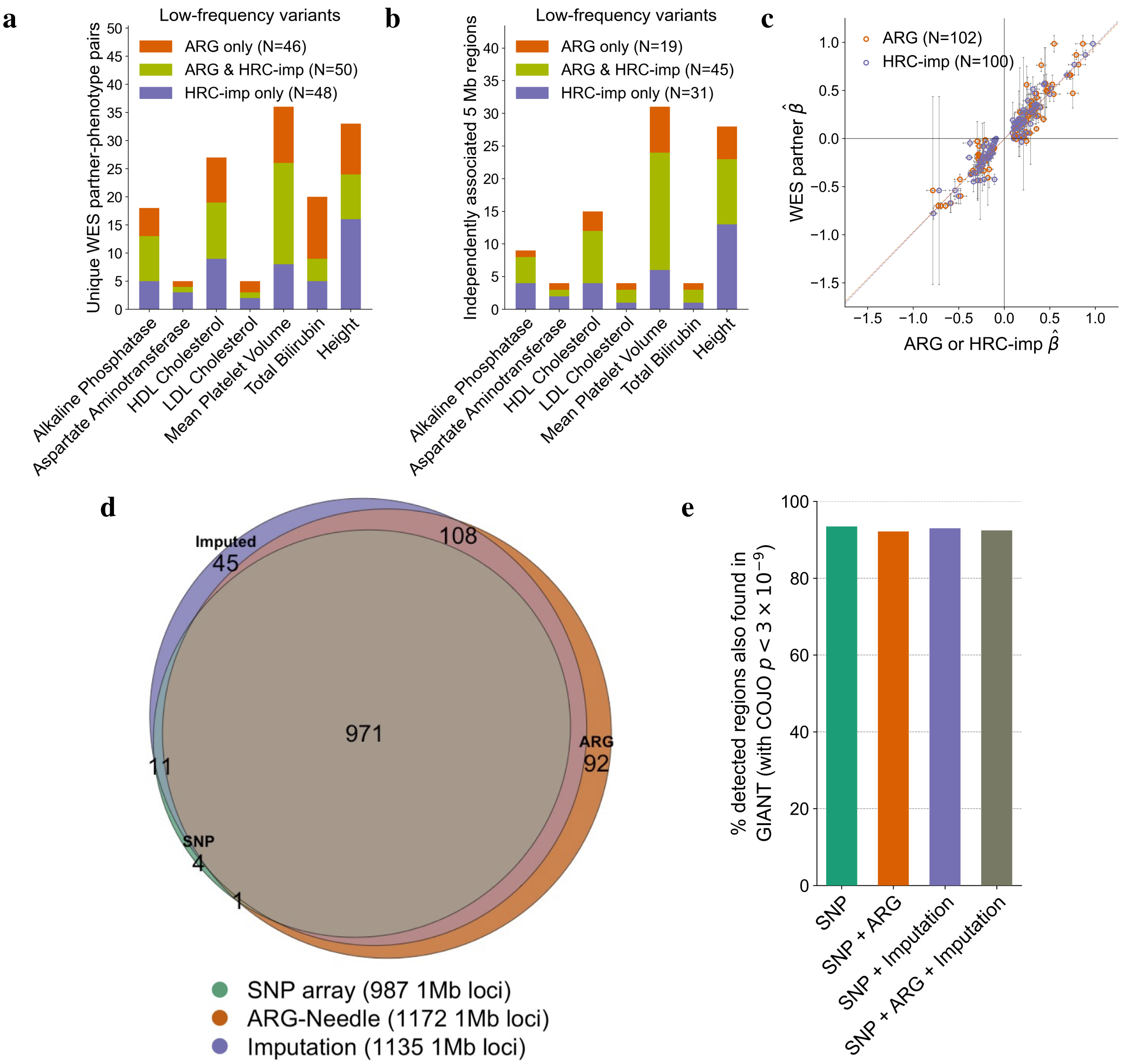

Figure S9: **Additional results for low ($0.1\% \leq$ MAF $< 1\%$) and high frequency (MAF $\geq 1\%$) variant associations. a-c**. Association of ARG-derived and imputed low-frequency variants with 7 quantitative traits. **a**. Counts of unique WES partners for ARG and HRC+UK10K imputed ("HRC-imp") independent associations, partitioned by traits and showing overlap. **b**. Counts of implicated 5 Mb regions containing ARG and HRC+UK10K imputation independent associations, partitioned by traits and showing overlap. **c**. Scatter plot of estimated effect ($\hat{\beta}$) for independent variants against estimated effect for their WES partners, with linear model fit. Error bars represent 1.96 s.e. **d-e**. Association of higher frequency variants with height. **d**. Venn diagram of number of 1 Mb regions containing a significant hit at $p < 3 \times 10^{-9}$ for ARG-Needle ($\mu = 10^{-5}$), HRC+UK10K imputed, and SNP array association. ARG-Needle association detected 971 out of 982 (98.9%) 1 Mb regions found by both imputation and array, 108 out of 153 (71%) 1 Mb regions found by imputation but not array, and an additional 92 (8% increase upon 1140) 1 Mb regions to those already found by imputation and array. **e**. Percent of 1 Mb regions containing independent associations (COJO $p < 3 \times 10^{-9}$) in association scans of 337,464 UK Biobank individuals that were also present in a GIANT consortium meta-analysis of $\sim$700K samples.

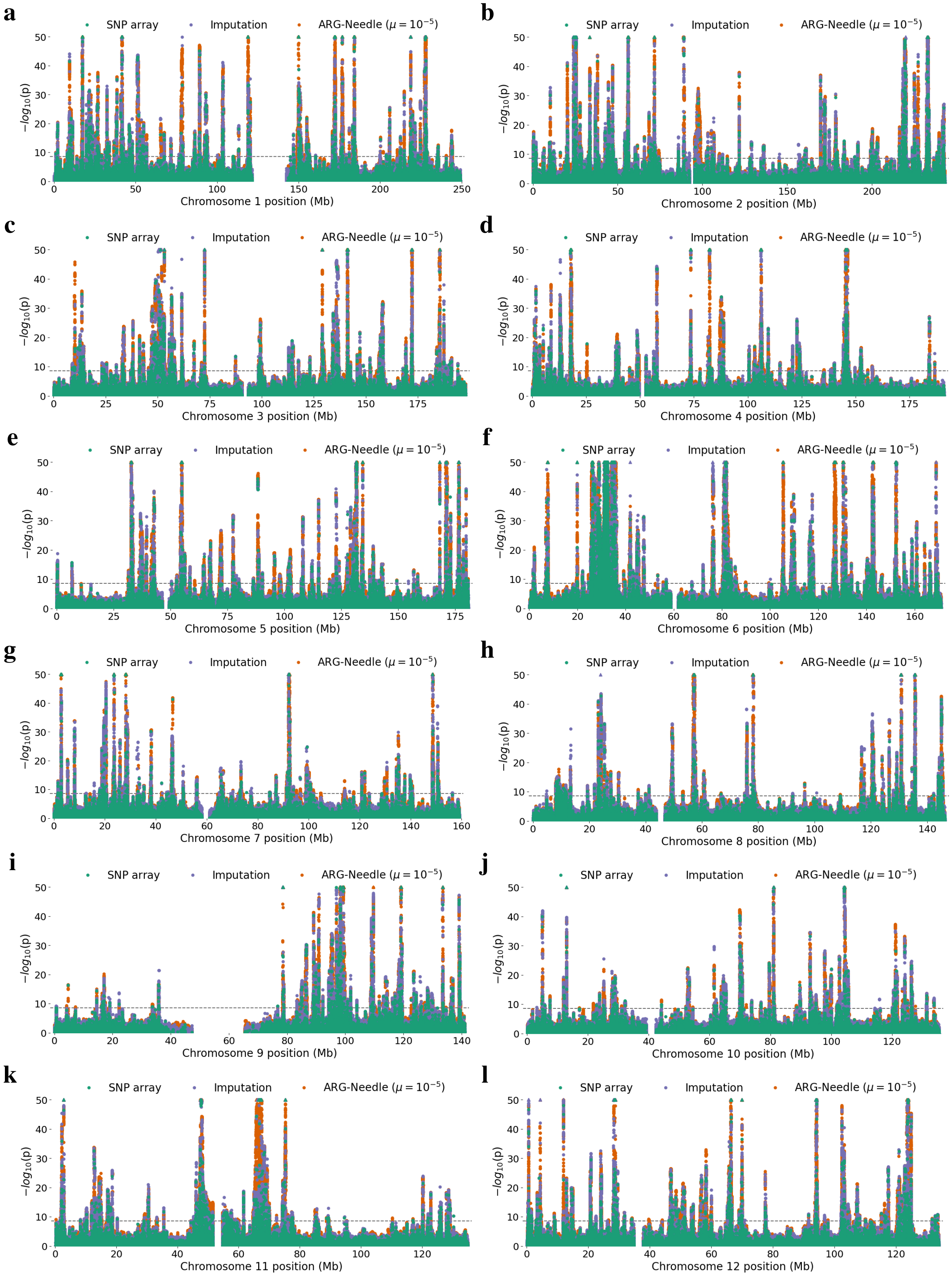

a
SNP array
Imputation
ARG-Needle ($\mu = 10^{-5}$)
$-log_{10}(p)$
Chromosome 1 position (Mb)
b
SNP array
Imputation
ARG-Needle ($\mu = 10^{-5}$)
$-log_{10}(p)$
Chromosome 2 position (Mb)
c
SNP array
Imputation
ARG-Needle ($\mu = 10^{-5}$)
$-log_{10}(p)$
Chromosome 3 position (Mb)
d
SNP array
Imputation
ARG-Needle ($\mu = 10^{-5}$)
$-log_{10}(p)$
Chromosome 4 position (Mb)
e
SNP array
Imputation
ARG-Needle ($\mu = 10^{-5}$)
$-log_{10}(p)$
Chromosome 5 position (Mb)
f
SNP array
Imputation
ARG-Needle ($\mu = 10^{-5}$)
$-log_{10}(p)$
Chromosome 6 position (Mb)
g
SNP array
Imputation
ARG-Needle ($\mu = 10^{-5}$)
$-log_{10}(p)$
Chromosome 7 position (Mb)
h
SNP array
Imputation
ARG-Needle ($\mu = 10^{-5}$)
$-log_{10}(p)$
Chromosome 8 position (Mb)
i
SNP array
Imputation
ARG-Needle ($\mu = 10^{-5}$)
$-log_{10}(p)$
Chromosome 9 position (Mb)
j
SNP array
Imputation
ARG-Needle ($\mu = 10^{-5}$)
$-log_{10}(p)$
Chromosome 10 position (Mb)
k
SNP array
Imputation
ARG-Needle ($\mu = 10^{-5}$)
$-log_{10}(p)$
Chromosome 11 position (Mb)
l
SNP array
Imputation
ARG-Needle ($\mu = 10^{-5}$)
$-log_{10}(p)$
Chromosome 12 position (Mb)

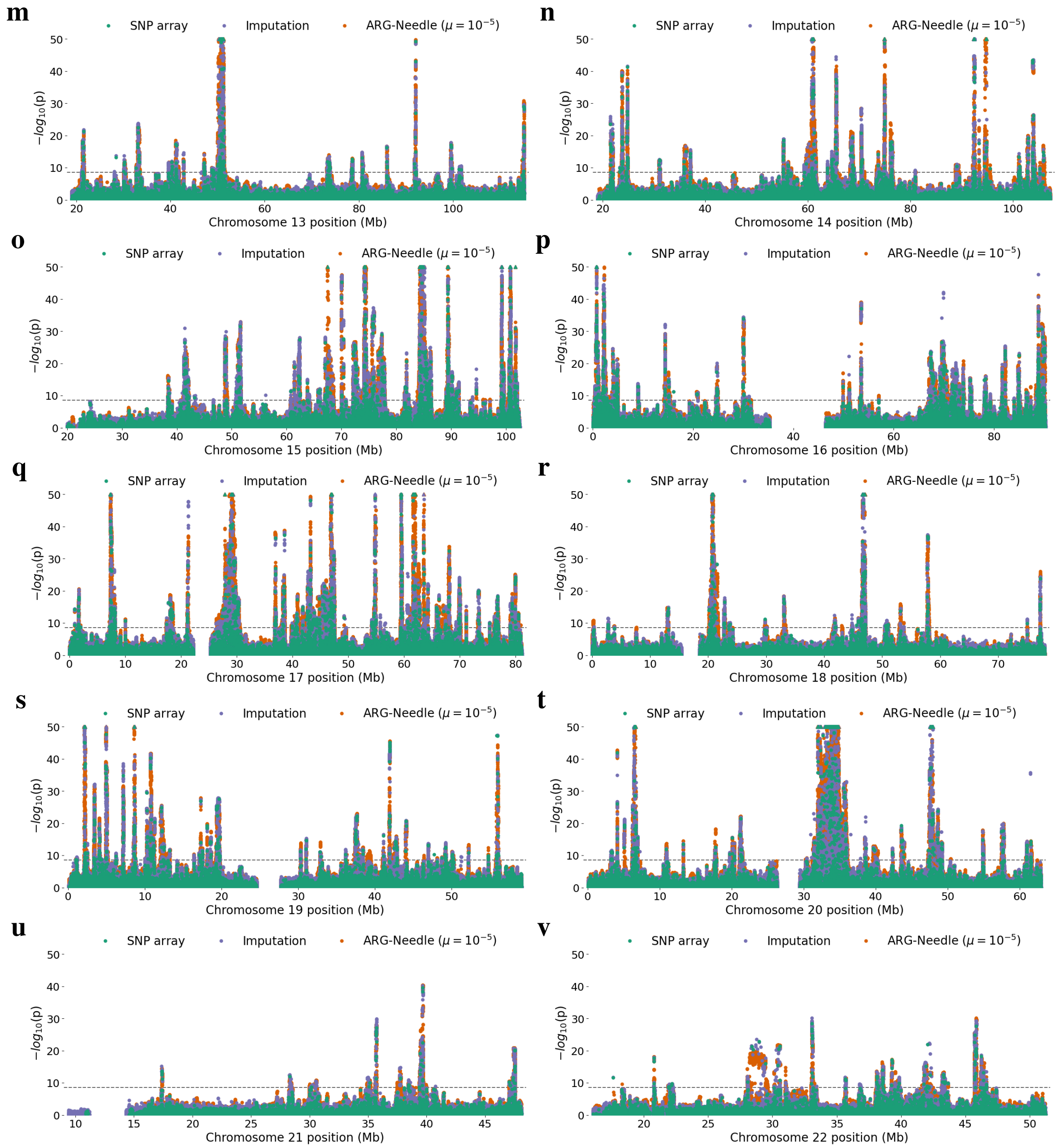

Figure S10: (Continued from previous page.) **Additional chromosome-wide Manhattan plots of mixed-model association of higher frequency variants with height.** Manhattan plots showing ARG-Needle, HRC+UK10K imputed variants, and SNP array association, as in Fig. 5a-b but with all methods on one plot and for all 22 chromosomes. Dotted lines correspond to $p = 3 \times 10^{-9}$ (see Methods). Triangles indicate associations with $p < 10^{-50}$. The order of plotting is ARG-Needle with $\mu = 10^{-5}$, then imputation, then SNP array variants on top.

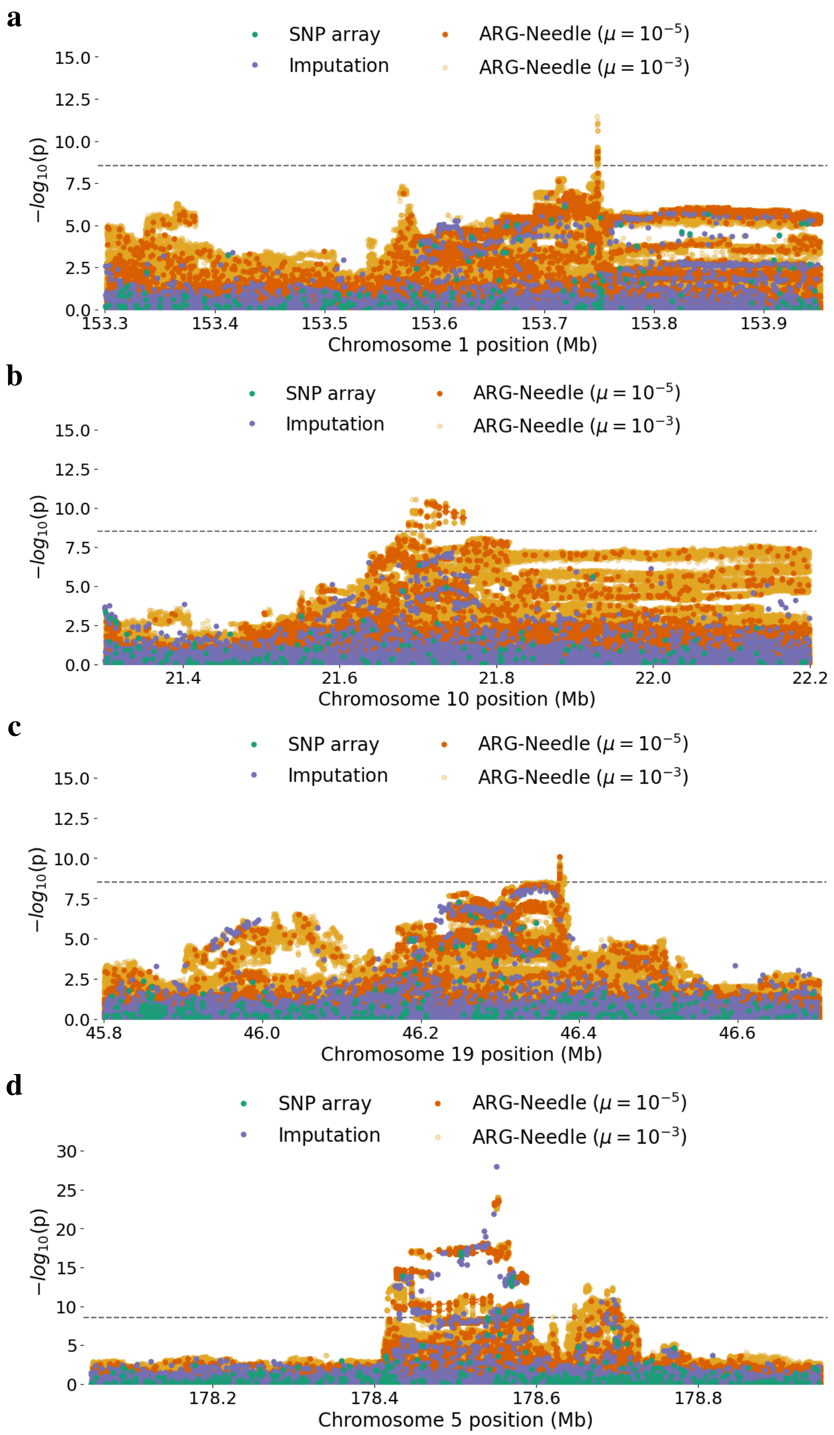
a
SNP array
Imputation
ARG-Needle ($\mu = 10^{-5}$)
ARG-Needle ($\mu = 10^{-3}$)
$-log_{10}(p)$
15.0
12.5
10.0
7.5
5.0
2.5
0.0
153.3
153.4
153.5
153.6
153.7
153.8
153.9
Chromosome 1 position (Mb)
b
SNP array
Imputation
ARG-Needle ($\mu = 10^{-5}$)
ARG-Needle ($\mu = 10^{-3}$)
$-log_{10}(p)$
15.0
12.5
10.0
7.5
5.0
2.5
0.0
21.4
21.6
21.8
22.0
22.2
Chromosome 10 position (Mb)
c
SNP array
Imputation
ARG-Needle ($\mu = 10^{-5}$)
ARG-Needle ($\mu = 10^{-3}$)
$-log_{10}(p)$
15.0
12.5
10.0
7.5
5.0
2.5
0.0
45.8
46.0
46.2
46.4
46.6
Chromosome 19 position (Mb)
d
SNP array
Imputation
ARG-Needle ($\mu = 10^{-5}$)
ARG-Needle ($\mu = 10^{-3}$)
$-log_{10}(p)$
30
25
20
15
10
5
0
178.2
178.4
178.6
178.8
Chromosome 5 position (Mb)

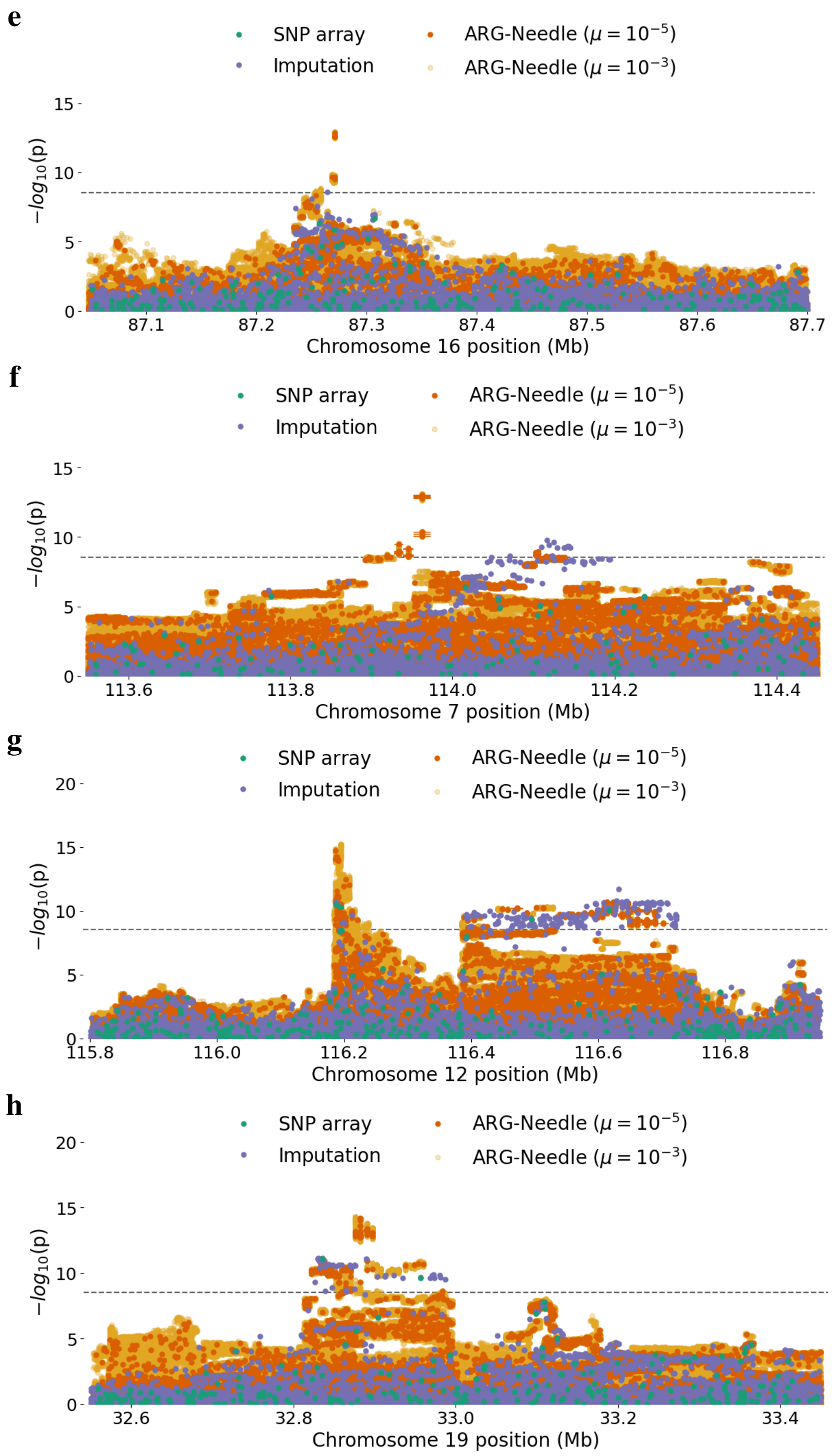

Figure S11: (Continued from previous page.) **Manhattan plots of higher frequency loci associated with height.** As in Fig. 5e-f but with eight additional loci of interest. **a-c**. Three regions where ARG-Needle ($\mu = 10^{-5}$) alone detects associations passing $p < 3 \times 10^{-9}$ significance. **d-e**. Two loci where ARG-Needle detects an association peak within 10 kb as HRC+UK10K imputation from $\sim$65K haploid references, despite only using SNP array data. **f-h**. Three loci where ARG-Needle detects a different primary association peak than SNP array or imputed data association. Dotted lines correspond to $p = 3 \times 10^{-9}$ (see Methods). The order of plotting is ARG-Needle with $\mu = 10^{-3}$, then ARG-Needle with $\mu = 10^{-5}$, then imputation, then SNP array variants on top. For the $\mu = 10^{-5}$ ARG associations crossing significance (here and in Fig. 5e-f), we additionally plotted a horizontal line showing the extent of the corresponding ARG clade.

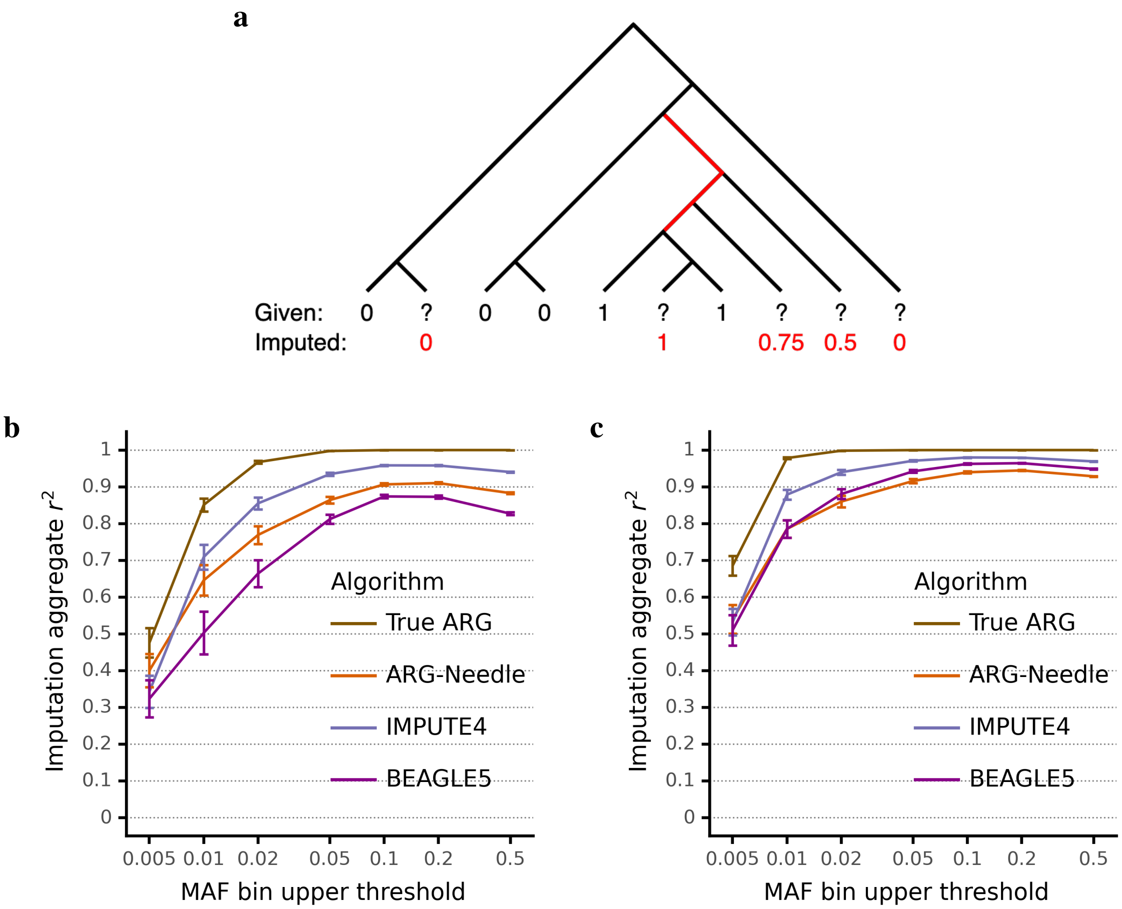

Figure S12: **Joint array and sequencing ARG inference for genotype imputation.** **a**. Given a polymorphic sequenced site containing sequenced samples, unobserved genotypes for array samples, and a marginal coalescent tree relating all samples, we perform genotype imputation as follows. We first identify all branches in the tree for which a mutation on that branch best explains the observed sequencing data in terms of Hamming distance (red branches in the example). Each branch implies genotypes of 0 or 1 for the array samples, and we weight by branch length to produce a weighted predicted dosage for each array sample. In this example, the three branches have lengths in ratio 1:1:2, resulting in the predicted dosages shown in red. **b**-**c**. We perform ARG-based imputation using the true ARG and an ARG-Needle inferred ARG and compare to IMPUTE4 and Beagle 5. Simulations use a 10 Mb region, 300 (**b**) or 1000 (**c**) haploid sequencing samples, and 1000 haploid array samples. For IMPUTE4 and Beagle 5, we input a genetic map corresponding to the recombination rate used for simulation, and otherwise use default parameters. Variants are binned by MAF in the sequencing samples, and we report the aggregate $r^2$ within each bin (mean over 25 runs), with each bin represented by its maximal MAF. Error bars represent 2 s.e.

### Supplementary Text Contents

**Supplementary Note 1** **18**

**References (Supplementary Information)** **42**

#### Supplementary Note 1: Additional details on ARG inference

In this note we clarify our definition of the ARG, provide detail for the three steps of the ARG-Needle algorithm (see Fig. 1), describe the ARG normalization step, and discuss a memory-efficient extension of the ASMC-clust algorithm.

##### ARG definition and representation

Several definitions of the ancestral recombination graph [1] (ARG) are used in the literature. The definitions depend on factors such as whether recombination events are explicitly represented, and whether the inferred structure is a graph or a collection of marginal trees. We therefore briefly clarify the definition and representation of the ARG used in this work.

##### Representing recombination events and individuals

The coalescent with recombination process [2, 3] (hereafter "the coalescent") is a stochastic process that produces an ARG. It is convenient to describe the coalescent by means of the simulation algorithm that samples a specific ARG instance. In its simplest form (ignoring e.g. mutations, which do not affect the shape of the ARG, as well as migration and gene conversion), backwards-in-time coalescent simulation proceeds by sampling coalescence and recombination events. Individuals are represented by collections of haplotypes, which may be encoded as a set of nodes with properties such as start/end position and time (or age). As simulation proceeds, a graph (the ARG) is built: new nodes are created and connected by new edges as a result of coalescence or recombination. Note that here we always refer to Hudson's ARG [3, 4], or the "small ARG", where only relevant ancestral material is considered [5].

The genomic location and time of recombination events may be explicitly modeled using the nodes and edges of the ARG, but their recording has no direct effect on the most common downstream analyses, such as the sampling of mutations and the generation of sequencing data. Although some authors refer to ARGs that explicitly model the location and time of recombination events (e.g. [6]), we do not make that distinction in this work. In particular, although we infer recombination locations using a heuristic as part of the ARG-Needle threading procedure (see the description of Step 3 below), the ARGs we produce do not store explicit recombination times. These may be added by performing a post-processing of the ARG to infer the most plausible time of recombination events, which we do not explore in this work. Our inferred ARGs also do not distinguish between an ancestral haplotype and an ancestor (which may be modeled as a collection of ancestral haplotypes), so that the ancestors in our ARGs never include "trapped material" [7].

##### Tree-based vs graph-based ARG representation

One key property of the ARG is that although it is a potentially cyclic graph (ignoring directionality of the edges; haplotypes may recombine and subsequently coalesce again), it simplifies to a tree if we only consider the set of nodes and edges spanning a specific genomic position. For this reason, an ARG may also be interpreted as a sequence of trees, and vice versa (if trees are defined for the same samples). This property is leveraged in formulations of the coalescent as a process that generates a series of correlated marginal trees along the genome, and in simulators of this process [8, 9, 10, 11, 12, 13]. However, despite this equivalence between tree-based and graph-based encoding of an ARG, the graph-based representation that more naturally arises in the backwards-in-time view of the process is more memory-efficient than a representation based on a collection of marginal trees [14], which requires storing of the same ancestral haplotype (ARG node) multiple times in neighboring trees.

Due to these different interpretations and properties, other approaches for genealogical inference sometimes refer to sequences of trees and ARGs with subtle distinctions. [14] and [15] (within the `tskit` library) use the graph-based ARG representation (a set of nodes and edges), but refer to it as a tree sequence, highlighting the sequential interpretation of an ARG. The output of Relate [16] is referred to as a sequence of trees approximating the ARG. Although these are equivalent in our definition, this places an emphasis on the fact that the Relate algorithm does not strongly enforce correlation across neighboring marginal trees (making the graph-based representation converge towards the tree-based representation). In our work we use a graph-based representation of the ARG adapted from the representation used within the ARGON simulator [17]. Although, due to intrinsic properties of ARGs, this graph may also be interpreted as representing a sequence of trees, we solely refer to it as an ARG, implying no particular difference. Because the ARG-Needle algorithm enforces some degree of node sharing across neighboring marginal trees, the graph-based representation of ARG-Needle inferred ARGs is more parsimonious than a representation based on independent marginal trees. Similarly to the Relate algorithm, on the other hand, the ASMC-clust algorithm does not encourage substantial correlation across marginal trees.

##### ARG-Needle ARG format

ARG-Needle is coded in C++ and Python. In an ARG-Needle ARG, nodes represent ancestral haplotypes, storing metadata such as their age (in generations). Edges connect pairs of nodes implying ancestor/descendant relationships, and store start and end coordinates for the inherited chromosomal region. Access to a node's ancestor or children at a particular position requires $O(\log K)$ computation, where $K$ is the total number of ancestors or children proceeding from the node. ARG-Needle implements various algorithms for complex trait analysis and can export and import ARGs in `tskit` format.

##### ARG-Needle algorithm description

ARG-Needle starts with an empty ARG, and iteratively adds one new haploid sample at a time to the ARG. Adding the first sample generates a single ARG node at the present time spanning the whole genomic region. For each additional sample, three steps are used to generate a threading instruction and add the new sample to the ARG: (1) perform hash table queries; (2) run ASMC [18] to form the threading instruction; (3) thread the new sample to the ARG. We describe each of these three steps in detail.

**ARG-Needle step 1: Shortlisting of closest relatives via genotype hashing**

Let the haploid data consist of $N$ samples and $M$ sites. Let the genetic distances of the sites be $g_1, \ldots, g_M$, and let the genotypes be $x_{ij} \in \{0, 1\}$, for $1 \leq i \leq N$ and $1 \leq j \leq M$. The parameters of the hash table operations are a hash word size $S$, a hash region size $L$, a hash tolerance $T$, and a hashing output size $K$. The units of the hash region size $L$ are in genetic distance, while all other parameters are integers. $S$ and $L$ are used to partition the sites into words and regions. $S$ simply denotes the number of sites in each word. For $1 \leq k \leq \lceil M/S \rceil$, word $k$ consists of sites $a(k) = (k-1) \times S + 1$ to $b(k) = \max(k \times S, M)$ inclusive, resulting in a total of $W = \lceil M/S \rceil$ words. For $1 \leq i \leq N$ and $1 \leq j \leq W$, let $w_{ik}$ be the integer resulting from interpreting the $W$ bits of the $k$th word for sample $i$ as a binary number, i.e. $w_{ik} = (x_{ib(k)} \ldots x_{ia(k)})_2$.

The words are further grouped into non-overlapping regions such that each region spans a genetic distance of at least $L$, with the exception of when $L > g_M - g_1$, in which case all words are placed in the same region. Starting with word 1, we find the first index $k$ such that $g_{b(k)} - g_1 \geq L$. This makes up the first region, spanning words 1 to $k$ inclusive. For the second region, we start at word $k+1$ and find the first index $k'$ such that $g_{b(k')} - g_{a(k)} \geq L$. The second region then spans words $k+1$ to $k'$. We continue in a similar fashion. When we reach the end, if we perfectly have just formed a last region covering all words, the algorithm terminates. Otherwise, we combine any remaining words on to the previous region so that all regions are of genetic distance approximately or just over $L$. Let the number of regions resulting from this procedure be $R$, and for $1 \leq r \leq R$, let $c(r)$ and $d(r)$ be the start and end words, inclusive, $1 \leq c(r) \leq d(r) \leq W$.

The hashing query takes a new sample in input and outputs the top $K$ candidate closest relatives to this sample out of those already in the ARG, for each of the $R$ regions. The purpose of having the hash region size $L$ be in genetic distance is to control for the expected number of recombination events within each region. Recombination will alter the set of closest relatives, but by keeping $L$ sufficiently low it is possible to obtain closest relatives that remain consistent through the region. Within each region, the hashing query selects the top $K$ candidate closest relatives by a score which we call the $T$-tolerant IBS score (for identical-by-state). Given two samples $i$ and $i'$ and a hash tolerance $T$, we say that $i$ and $i'$ are $T$-tolerant IBS over $[f, g]$, $1 \leq f \leq g \leq W$, if

$$|\{f \leq k \leq g | w_{ik} \neq w_{i'k}\}| \leq T.$$

The $T$-tolerant IBS score of samples $i$ and $i'$ over a region $r$ is the number of words in the longest $T$-tolerant IBS stretch with at least one word of overlap with region $r$. In notation, we consider the set

$$\{[f, g] | 1 \leq f \leq g \leq W, f \leq d(r), g \geq c(r), |\{f \leq k \leq g | w_{ik} \neq w_{i'k}\}| \leq T\}$$

and take the maximum value of $g - f + 1$ over this set. For each region $r$, we then take the top $K$ samples with the largest $T$-tolerant IBS score over this region. In the case of ties, we select the sample(s) that have been added to the ARG more recently, though it is also possible to break ties randomly. If the number of samples with a nonzero score is less than $K$, only this many results are returned, or in other words, samples scoring zero are not returned. In summary, the hashing query returns a list of at most $K$ top-scoring sample IDs for each of the $R$ regions.

Our hashing data structure consists of a vector of $W$ hash tables, one for each of the $W$ words. The $k$th hash table maps from possible values for the $k$th word, guaranteed to be a number between 0 and $2^S - 1$ inclusive, to a vector of sample IDs which contain that value for the $k$th word. Every time a sample $i'$ is added to the ARG, we perform $O(W)$ operations to update this data structure. For each $k$, $1 \leq k \leq W$, we access the entry in the $k$th hash table at $w_{i'k}$, initializing an empty vector if no entry exists, and appending $i'$ to this vector. Similarly, when we want to query a new sample $i'$, we can index $w_{i'k}$ into the $k$th hash table to find the samples which match sample $i'$ at the $k$th word. In this way, we only need to visit the values $(i, k)$ for which there is a match, $w_{ik} = w_{i'k}$.

Suppose there are $N'$ samples already in the ARG, and we want to query the closest relatives for a new sample $i'$. Our hashing-based implementation runs in $O\left(\overline{N}_{overlap}W\right)$, where $0 \leq \overline{N}_{overlap} \leq N'$ is the average number of samples already in the ARG which match sample $i'$ per word, i.e. the average size of $\{0 \leq i \leq N' | w_{ik} = w_{i'k}\}$ as $k$ varies from 1 to $W$. As one increases the word size $S$, $W = \lceil M/S \rceil$ decreases and $\overline{N}_{overlap}$ also decreases because longer words are less likely to be shared. The pattern of this latter trend as a function of $S$ is data set-specific due to factors such as the demographic history of the population, but in summary, $S$ can be used as a parameter to increase hashing speed at the cost of coarser hashing-based IBS detection.

In building ARGs on 337K UK Biobank samples, the hashing step dominates runtime. We implemented a form of dynamic hashing that uses a primary hash word size $S_1$ and falls back to a backup hash word size $S_2$ in some cases, with $S_1 > S_2$. The criterion for falling back to the backup hash word size is parameterized by a factor $F$. For each region, we compute the sum of the top $K$ $T$-tolerant IBS scores in that region, and check that this is greater than $F$ times the number of words in the region (in notation, $F \times (d(r) - c(r) + 1)$). If the check fails for any of the regions, then we fall back to the backup hash word size, repeating the entire hash query with $S_2$ instead of $S_1$. This has the effect of performing the initial hashing queries with $S_2$, then transitioning to more and more queries done with $S_1$ as the ARG-Needle algorithm proceeds. For our run on 337K diploid individuals, we started with $S_1 = 16, S_2 = 8, F = 4$ for adding the first 50K individuals, then progressed to $S_1 = 64, S_2 = 16, F = 8$ for the remaining individuals. Other parameters were set to $T = 1$, $K = 64$, and $L = 0.5$ cM.

##### ARG-Needle step 2: ASMC queries

In step 2 of the ARG-Needle algorithm, the candidate closest relatives output by hashing are validated using ASMC to yield a more certain estimate of true closest relatives and their TMRCA to the target sample. Each pairwise ASMC comparison takes $O(MD)$ time to run, where $D$ is the number of discretized time bins used to represent the posterior distribution of pairwise coalescence time (we use $D = 69$, the default for ASMC v1.0). With no hashing, each new sample being threaded would need to be compared against all existing samples in the ARG using ASMC, yielding $O(N^2MD)$ runtime overall. By only selecting $K$ candidate closest relatives for each sample being threaded, the total runtime is instead linear in $N$. Step 1 of ARG-Needle thus functions as a heuristic pre-selection strategy, similar to the use of genotype hashing or the positional Burrows-Wheeler transform [19] to speed up genotype imputation [20, 21, 22], phasing [23], and identity-by-descent (IBD) detection [24].

For each of the $R$ regions (partitioned based on genetic distance, see above), step 1 of ARG-Needle returns up to $K$ IDs representing the candidate closest relatives to this sample within the region. Using our earlier definitions, region $r$ contains words with indices from $c(r)$ to $d(r)$ inclusive, with $1 \leq c(r) \leq d(r) \leq W$, which means it contains SNPs with indices from $a(c(r))$ to $b(d(r))$ inclusive, $1 \leq a(c(r)) \leq b(d(r)) \leq M$. We use ASMC (run in array or sequencing mode, depending on the experiment) to predict the pairwise coalescence time posterior distribution between the new sample and each of the $K$ samples at each of these SNP positions, resulting in an output of size $K \times (b(d(r)) - a(c(r)) + 1) \times D$. Because the ASMC Hidden Markov Model (HMM) takes into account sequential context when making a prediction, we pad the input region by adding 1 cM on either side. This increases accuracy at the cost of slightly increased runtime. The ASMC output for each pair and site is a posterior distribution, given as probabilities over the $D$ discretized time bins. We use it to extract the posterior mean and the mode, or maximum a posteriori (MAP) values. This does not affect the runtime complexity, as for each site and pair computing these summaries takes $O(D)$ time. Before proceeding to step 3, we aggregate these summaries so that at each site we have at most $K$ triplets, which represent the ID of the candidate closest relative, the posterior mean of the TMRCA with the target sample, and the MAP.

##### ARG-Needle Step 3: processing the ASMC output and performing threading

In the final step of the ARG-Needle algorithm, we use the ASMC output to add the new sample to the ARG. This takes place using a "threading" operation, named after the terminology coined by ARGweaver [6]. We first describe the ARG-Needle threading operation, then show that it can be used to accurately reconstruct an ARG, and finally explain how we perform threading using the ASMC output.

We define a "threading instruction" to contain the sample ID of a closest relative and their TMRCA to the target sample, at each position along the genome. If we assume a genomic extent $[s, t)$ and sample IDs $\{1, \ldots, N'\}$ already in the ARG, a threading instruction may be represented as a function $f : [s, t) \to \{1, \ldots, N'\} \times (0, \infty)$, assigning to each position $x$ a pair $f(x) = (i(x), T(x))$, where $i(x)$ is a sample ID and $T(x) > 0$ is a time.

We now describe the threading operation in terms of how it affects each marginal tree. (Our actual implementation considers threading as an ARG-wide problem and is more efficient; we take this perspective for simplicity.) Consider a position $x$ with a marginal tree over $N'$ samples, and suppose the threading instruction has $f(x) = (i, T)$. First, a new node $u$ for sample $N' + 1$ is created. Next, let

$v$ be the oldest ancestor node of sample $i$ with a time less than or equal to $T$. One of three conditions holds:

1. If $v$ has time less than $T$ and $v$ has a parent node $v'$ (necessarily with time greater than $T$), create a new node $w$ with time $T$. Delete the edge between $v$ and $v'$ and add three new edges between pairs ($u$, $w$), ($v$, $w$), and ($w$, $v'$).
2. If $v$ has time less than $T$ but does not have a parent node, then $v$ is the root node of the marginal tree at $x$. Create a new node $w$ with time $T$ and add two new edges between pairs ($u$, $w$) and ($v$, $w$). (Node $w$ becomes the new root node.)
3. If $v$ has time $T$, then create an edge going from $u$ to $v$. (This operation creates polytomies: nodes with more than two children at a position.)

Next, we show that iteratively threading samples to an ARG using accurate threading instructions (reflecting true TMRCAs) leads to recovering the true ARG. Let tmrca$(a, b)$ denote the pairwise TMRCA between samples $a$ and $b$ (position $x$ is implicit). We begin with a few observations (these claims are easily verified, and we omit a proof):

*Claim 1.* For samples $1 \leq a < b \leq N'$ (samples already in the ARG), tmrca$(a, b)$ is the same before and after the above threading operation.

*Claim 2.* After the threading operation, tmrca$(N' + 1, i) = T$.

The remaining pairwise TMRCAs after threading can also be determined:

*Claim 3.* Let $j$ be any sample in the ARG other than $i$. After the threading operation, we have

$$\text{tmrca}(N' + 1, j) = \max\left(T, \text{tmrca}(i, j)\right).$$

*Proof.* Let $v$ be the pairwise MRCA node of samples $i$ and $j$, and $w$ be the pairwise MRCA node of samples $N' + 1$ and $i$ after threading (having time $T$). There are now three cases:

1. If tmrca$(i, j) < T$, then $v$ lies below $w$, and there is a path from the node for sample $i$ to $w$ which must pass through $v$. Therefore $w$ is the MRCA node of $j$ and $N' + 1$, so tmrca$(N' + 1, j) = T = \max\left(T, \text{tmrca}(i, j)\right)$.

2. If tmrca$(i, j) > T$, then $w$ lies below $v$. The MRCA of $w$ (a non-leaf node) and sample $j$ is $v$, and sample $N' + 1$ is a descendant of $w$, so tmrca$(N' + 1, j) =$ tmrca$(i, j) = \max\left(T, \text{tmrca}(i, j)\right)$.

3. If tmrca$(i, j) = T$, then when sample $N' + 1$ was being threaded to the sample $i$, it would have found node $v$ already at time $T$, leading to the second threading case above and adding a single edge between the node for sample $N' + 1$ and $v$. Therefore tmrca$(N' + 1, j) = T = \max\left(T, \text{tmrca}(i, j)\right)$.

□

We leverage these observations to verify that the threading algorithm reconstructs the true ARG when the threading instructions are correct (i.e., when at each position the threading instruction provides a true closest relative and their true TMRCA to the target sample).

*Theorem.* Let $\mathcal{A}$ be an ARG over $N$ samples, and let $\mathcal{B}$ be an ARG constructed via threading from an initially trivial ARG with one sample. Suppose that for threading sample $N'$, one uses the threading instruction $f(x) = (i(x), T(x))$ consisting of $i(x) = \text{argmin}_{1 \leq i' < N'}(\text{tmrca}_{\mathcal{A},x}(N', i'))$ and $T(x) = \min_{1 \leq i' < N'}(\text{tmrca}_{\mathcal{A},x}(N', i'))$. Then at each $x$, the marginal trees of $\mathcal{A}$ and $\mathcal{B}$ will be identical (not considering the possible creation or deletion of unary nodes).

*Proof.* It suffices to consider an arbitrary position $x$ and show that the marginal trees of $\mathcal{A}$ and $\mathcal{B}$ are equivalent. We therefore rely on the fact that the full set of TMRCAs uniquely determines a rooted tree [25], and show that the $\binom{N}{2}$ pairwise TMRCAs between any two samples are the same within $\mathcal{A}$ and $\mathcal{B}$. We proceed by induction, and assume that the pairwise TMRCAs between the first $N'$ samples are the same in $\mathcal{A}$ and $\mathcal{B}$. The pairwise TMRCAs between $N' + 1$ and the first $N'$ samples are set when sample $N' + 1$ is threaded to $\mathcal{B}$, and unchanged after (by Claim 1). Let $i$ and $T$ denote the chosen sample and time as defined above, then by Claim 2, $\text{tmrca}_{\mathcal{B}}(N' + 1, i) = T = \text{tmrca}_{\mathcal{A}}(N' + 1, i)$. It remains to show that $\text{tmrca}_{\mathcal{B}}(N' + 1, j) = \text{tmrca}_{\mathcal{A}}(N' + 1, j)$ for any $j \neq i$ among the first $N'$ samples. We have

$$\begin{aligned} \text{tmrca}_{\mathcal{B}}(N' + 1, j) &= \max\left(T, \text{tmrca}_{\mathcal{B}}(i, j)\right) \\ &= \max\left(T, \text{tmrca}_{\mathcal{A}}(i, j)\right), \end{aligned}$$

by Claim 3 and the inductive hypothesis. By the definition of $T$,

$$\begin{aligned} \text{tmrca}_{\mathcal{B}}(N' + 1, j) &= \max\left(T, \text{tmrca}_{\mathcal{A}}(i, j)\right) \\ &\leq T \\ &= \min_{1 \leq i' \leq N'}(\text{tmrca}_{\mathcal{A}}(N' + 1, i')) \\ &\leq \text{tmrca}_{\mathcal{A}}(N' + 1, j). \end{aligned}$$

However, by the ultrametric property,

$$\begin{aligned} \text{tmrca}_{\mathcal{B}}(N' + 1, j) &= \max\left(T, \text{tmrca}_{\mathcal{A}}(i, j)\right) \\ &= \max\left(\text{tmrca}_{\mathcal{A}}(N' + 1, i), \text{tmrca}_{\mathcal{A}}(i, j)\right) \\ &\geq \text{tmrca}_{\mathcal{A}}(N' + 1, j). \end{aligned}$$

Putting these together, $\text{tmrca}_{\mathcal{B}}(N' + 1, j) = \text{tmrca}_{\mathcal{A}}(N' + 1, j)$. □

Having verified that iteratively threading all samples using accurate threading instructions recovers the true ARG, we describe our strategy to estimate the threading instructions from the output of step 2. At each site, the closest relative is selected to be the individual with smallest ASMC posterior mean TMRCA among the $K$ candidate closest relatives (ties are arbitrarily broken). To compute the TMRCA for the threading instruction, we select regions where both the closest relative and their TMRCA MAP remain constant and compute the average posterior mean value within each such region. This has several benefits over using the raw value of either the MAP or the posterior mean at each site. Although changes in the MAP better reflect IBD segment breakpoints, the MAP output by ASMC only takes one of $D$ possible values, which would lead to a large number of polytomies in the ARG. The MAP also has a higher RMSE for the TMRCA compared to the posterior mean. The posterior mean, on the other hand, takes different values at each site, which would reduce haplotype sharing across marginal trees and produce a less compact ARG (as in the case of ASMC-clust). By combining the MAP and posterior mean in the threading instruction we thus obtain better estimates for the location of recombination breakpoints without sacrificing accuracy or compactness of the inferred ARG.

##### ARG normalization

Although the posterior mean has a lower RMSE than the MAP in estimating TMRCAs, it tends to be more biased towards the average coalescent time induced by the demographic prior, particularly in sparser array data. For this reason, we observed that ARGs inferred using the above definition of threading instruction tend to overestimate the time of recent coalescence events and underestimate the height of root nodes. We thus developed a procedure, which we call ARG normalization, to further leverage the demographic prior to reduce the bias due to the use of posterior mean estimates, while preserving the inferred ordering of coalescent events (see Supplementary Fig. S3). ARG normalization performs a quantile normalization of the heights of inferred ARG nodes, rescaling them to match the quantiles observed in 1,000 independent trees sampled from the demographic prior (after accounting for the span of inferred ARG nodes).

More in detail, assuming the 1,000 simulations generate $Q$ non-leaf nodes with times that can be ordered increasingly from $t_1$ to $t_Q$, we compute quantiles from the simulated trees by assigning quantile $(2i-1)/(2Q)$ to time $t_i$. We also assign quantile 0 to time 0 and quantile 1 to time $1.05 \times t_Q$ to ensure that the mapping is strictly increasing. Given an inferred ARG, we sought to compute an analogous quantile distribution of node times that was sensitive to differences between nodes spanning short vs. long segments of ancestral material and sensitive to polytomies. For each edge in the inferred ARG, we recorded the time of its parent node and the distance in base pairs it spanned. We aggregated these time-distance pairs to obtain $Q'$ distinct parent node times, ordered increasingly from $t_1$ to $t_{Q'}$, and corresponding aggregate distances spanned by edges with that parent node time, $d_1$ to $d_{Q'}$. We assigned quantile value $(d_1 + d_2 + \ldots + d_{i-1} + d_i/2)/(d_1 + \ldots + d_{Q'})$ to time $t_i$. We then matched the inferred node time quantile distribution with the target node time quantile distribution using linear interpolation on the quantiles. We used this mapping to rewrite all nodes of time $t_i$ to the new corresponding time.

##### Memory-efficient ASMC-clust extension

Assuming $N$ samples and $M$ sites, the default ASMC-clust algorithm requires $O(N^2M)$ memory and runtime complexity. We implemented an option for ASMC-clust to instead use $O(N^2 + NM)$ memory, at the cost of more runtime. In the default algorithm, we run ASMC on all $\binom{N}{2}$ pairs of samples across

$M$ sites, and store the results in an $\binom{N}{2}$ by $M$ matrix. We then iterate through the TMRCA values at each of the $M$ sites and perform UPGMA hierarchical clustering. The resulting ARG consists of $M$ trees, each over $N$ nodes, which can be represented in $O(NM)$ memory. Therefore, the memory bottleneck comes from the storage of the TMRCA results, which takes up $O(N^2M)$ memory.

To reduce the memory required from storing the TMRCA results, we iterate through groups of $M_{max}$ sites, each time computing and storing the TMRCA results for those sites, performing UPGMA clustering, and saving the clustered trees. This results in $O(N^2M_{max} + NM)$ memory usage. As described earlier, however, we pad additional sites (by default, 1 cM) on either end of the region to provide the ASMC HMM with some context. Using this approach, each site will therefore be processed multiple times by ASMC, leading to a time complexity somewhere between $O(N^2M)$ and $O(N^2M^2)$, depending on the choice of $M_{max}$ and the number of SNPs provided as context in each ASMC run.

In our simulations we fixed $M_{max} = 300$ for SNP data and $M_{max} = 2000$ for sequencing data. This led to memory usage that scaled quadratically as a function of $N$. Relate faces a similar issue of potentially quadratic memory, which is also solved by only storing distance matrices for a certain number of sites at a time. However, instead of fixing the number of sites, Relate enforces a maximum memory for the storage of distance matrices and uses that to determine the appropriate number of sites. We ran Relate using its default setting, which sets the maximum amount of memory for these calculations to 5 GB.

#### Supplementary Note 2: Additional details about ARG evaluation metrics

We studied four specific metrics to compare true and inferred ARGs: Robinson-Foulds distance, ARG total variation distance, pairwise TMRCA RMSE, and Kendall-Colijn (KC) topology-only distance. We also looked at scatter plots of the predicted vs. true TMRCAs. We first describe our use of stabbing queries to calculate these metrics, then discuss the various evaluation methods in more detail.

##### Evaluating metrics via stabbing queries

Most of the metrics we considered—Robinson-Foulds distance, pairwise TMRCA RMSE, and KC topology-only distance—are originally defined on two trees. (The ARG total variation distance is defined directly between two ARGs and is treated separately.) To generalize these metrics to ARGs, we consider the metrics as comparing two marginal trees at the same position and take a genome-wide average of the metrics over all positions.

Consider two ARGs $\mathcal{A}$ and $\mathcal{B}$, which may for instance be the true and inferred ARGs, with a common genomic extent $[s, t) \subset \mathbb{R}$. (Note that we are modeling the genome as continuous.) Let Tree be a binary operator that takes an ARG and a position and returns the tree at that position, and suppose $d : \mathcal{T}_N \times \mathcal{T}_N \to \mathbb{R}$ is a metric of interest that operates on the space of rooted trees on $N$ leaves $\mathcal{T}_N$. Then the genome-wide average of $d$ is

$$\frac{1}{t-s}\int_s^t d(\text{Tree}(\mathcal{A}, x), \text{Tree}(\mathcal{B}, x))dx.$$

Although it is possible to compute such integrals exactly, we found that such a method is slowed down by needing to iterate over all marginal trees in both ARGs. In particular, the true ARG often contains many recombination events, each of which generates a new marginal tree. Therefore, we instead approximated the integral using sampling. Given a set of points $x_1, \ldots, x_n$ uniformly distributed among $[s, t)$, we have

$$\frac{1}{t-s}\int_s^t d(\text{Tree}(\mathcal{A}, x), \text{Tree}(\mathcal{B}, x))dx \approx \frac{1}{n}\sum_{i=1}^{n} d(\text{Tree}(\mathcal{A}, x_i), \text{Tree}(\mathcal{B}, x_i)).$$

We call each such $d(\text{Tree}(\mathcal{A}, x_i), \text{Tree}(\mathcal{B}, x_i))$ term a "stabbing query", because we are sampling two trees at a position along the ARG, which entails finding the edges of the ARG that overlap this position. The estimate is then an unweighted average over $n$ stabbing queries. For choosing the points $\{x_i\}_{i=1}^n$, we set $x_i = s + (i \cdot \phi - \lfloor i \cdot \phi \rfloor) \times (t - s)$, where $\phi = (1 + \sqrt{5})/2$ is the golden ratio. For all evaluations, we used $n =$ 5,000 stabbing queries.

##### Robinson-Foulds distance and ARG total variation distance

For our use of the Robinson-Foulds metric, we averaged the scaled Robinson-Foulds distance, which we defined in Methods, over $n =$ 5,000 genome-wide stabbing queries. This quantity lies between 0 (perfect match of present mutations) and 1 (complete mismatch of non-singleton mutations). Note that Robinson-Foulds weights all possible mutations in a marginal tree equally when considering dissimilarity. However, rare variants in these marginal trees tend to correspond to recent, short branches and are less likely to occur through random mutations. Furthermore, these short and recent branches tend to represent long haplotypes and thus appear in multiple neighboring trees, leading to a disproportionate contribution to the overall distance. Our proposed metric, the ARG total variation distance, generalizes the Robinson-Foulds distance to overcome these limitations when comparing ARGs.

To define the ARG total variation distance we focus on the probability distributions that two ARGs encode over the set of possible mutations found in the genome and compute the total variation distance between these distributions. To this end, given an ARG $\mathcal{A}$, we consider the probability that an ARG generates a variant represented by a specific bitset at a given genomic location. We are interested in this probability because we often infer the ARG from a subset of available markers but leverage it to infer the presence of other underlying variants. This probability thus encodes the confidence placed by the inferred ARG in the presence of an underlying variant, and can be compared to the likelihood of such a variant being generated by the true ARG.

More formally, given $N$ samples, the possible non-trivial mutational patterns that can occur over these samples are given by the set $\{0,1\}^N \setminus \{(0,\dots,0),(1,\dots,1)\}$. We call each element of this set an $N$-bitset; there are $2^N - 2$ possible $N$-bitsets, which we denote as $\mathcal{S}_N$. We choose to represent each mutation using its position $x \in [s,t)$ and the $N$-bitset $b \in \mathcal{S}_N$. We thus consider the probability distribution corresponding to the mutations $(x,b) \in [s,t) \times \mathcal{S}_N$ that can be observed. Assuming a constant mutation rate, each simulated mutation is uniformly distributed over the area of the ARG, which consists of the physical distance times the height of each mutation-generating branch. We can capture such a distribution via a probability density function (PDF) $f_{\mathcal{A}} : [s,t) \times \mathcal{S}_N \to \mathbb{R}$ which is induced by uniform sampling over the ARG. We can use two equivalent expressions for the total variation distance and apply them to compute the total variation distance between two ARGs $\mathcal{A}$ and $\mathcal{B}$ in terms of their PDFs:

$$TV_{ARG}(\mathcal{A},\mathcal{B}) = \frac{1}{2}\int_s^t \sum_{b\in\mathcal{S}_N} |f_{\mathcal{A}}(x,b) - f_{\mathcal{B}}(x,b)|dx \tag{4}$$

$$= 1 - \int_s^t \sum_{b\in\mathcal{S}_N} \min(f_{\mathcal{A}}(x,b), f_{\mathcal{B}}(x,b))dx. \tag{5}$$

The distance is 0 for identical distributions and 1 if the two distributions contain no overlapping support.

To evaluate the ARG total variation distance in our simulations, we used the second expression, where we approximated the integral by sampling $n =$ 5,000 stabbing queries and performed a sum over all present mutations for each stabbing query. We benchmarked the various methods across our simulation conditions with and without ARG normalization. Overall, ARG normalization either kept the total variation distance the same or provided a modest benefit (Supplementary Fig. S2b for methods without ARG normalization, and S2c for methods with ARG normalization).

To highlight the connection between the ARG total variation distance and Robinson-Foulds more explicitly, we first describe how we compute the PDF $f_{\mathcal{A}}(x,b)$ of an ARG $\mathcal{A}$. The value $f_{\mathcal{A}}(x,b)$ should be 0 if no branch yielding $N$-bitset $b$ exists at the tree at position $x$. Otherwise, because we are sampling uniformly, $f_{\mathcal{A}}(x,b)$ should be proportional to the length of the branch yielding $b$. Define a function $l_{\mathcal{A}} : [s,t) \times \mathcal{S}_N \to \mathbb{R}$ which will be a scaled version of $f_{\mathcal{A}}(x,b)$, and which we can use to describe $f_{\mathcal{A}}(x,b)$ explicitly:

$$l_{\mathcal{A}}(x,b) = \begin{cases} 0, & \text{if no branch giving } b \text{ exists at Tree}(\mathcal{A},x), \\ \text{the length of a branch giving } b \text{ at Tree}(\mathcal{A},x), & \text{otherwise.} \end{cases} \tag{6}$$

We can then write

$$f_{\mathcal{A}}(x,b) = \frac{1}{Z_{\mathcal{A}}} l_{\mathcal{A}}(x,b) \tag{7}$$

for some scaling constant $Z_{\mathcal{A}}$. We would like all the properties of a PDF to hold, e.g.

$$\int_s^t \sum_{b \in \mathcal{S}_N} f_{\mathcal{A}}(x,b) dx = 1. \tag{8}$$

Combining (7) and (8), we obtain

$$Z_{\mathcal{A}} = \int_s^t \sum_{b \in \mathcal{S}_N} l_{\mathcal{A}}(x,b) dx. \tag{9}$$

For a given ARG $\mathcal{A}$, equations (6), (7), and (9) provide a way to evaluate its PDF. First, we compute $Z_{\mathcal{A}}$ using stabbing queries. Then, to get the value of $f_{\mathcal{A}}(x,b)$, we look up $l_{\mathcal{A}}(x,b)$ using the marginal tree at $x$, and divide by $Z_{\mathcal{A}}$.

Consider again the definition of $l_{\mathcal{A}}(x,b)$. We can obtain a topology-only version of the ARG total variation distance by rewriting $l_{\mathcal{A}}(x,b)$ to not use branch lengths:

$$l'_{\mathcal{A}}(x,b) = \begin{cases} 0, & \text{if no branch giving } b \text{ exists at Tree}(\mathcal{A},x), \\ 1, & \text{otherwise.} \end{cases}$$

If we use the same definitions for $TV_{ARG}$ but replace $l_{\mathcal{A}}(x,b)$ with $l'_{\mathcal{A}}(x,b)$ and assume no polytomies in $\mathcal{A}$ or $\mathcal{B}$, the ARG total variation distance reduces to a scaled version of the Robinson-Foulds metric.

We briefly comment on limitations of the ARG total variation distance. In our definition, we considered the space of possible mutations as consisting of a position $x \in [s,t)$ and an $N$-bitset $b \in \mathcal{S}_N$. Including the position enables us to measure the ability to correctly localize mutations in the genome. It may also be worthwhile to also consider the time (height) of the mutations, which enable measuring the ability to correctly localize the time of mutation events. Additionally, the total variation distance relies on a hard 0-1 loss in its definition: it does not consider whether two mutations that are correlated (small Hamming distance) or occur at close but disjoint positions in two ARGs, a limitation it shares with the Robinson-Foulds distance. One possible direction to ameliorate this drawback is to use the Wasserstein distance, which generalizes the total variation distance with the aid of a metric on the probability space, thus allowing "margin for error" with a smooth loss.

##### Pairwise TMRCA RMSE and TMRCA scatter plots

We provide additional details on the TMRCA RMSE and its interpretation. Recall that we introduced ARG normalization as a way to adjust the node times of an inferred ARG to be more consistent with a demographic prior. We observed that ARG normalization improves the pairwise TMRCA RMSE of Relate in array data, though not in sequencing data (compare Supplementary Fig. S2d,f), and tends to improve performance for tsinfer as well. This suggests that ARG normalization provides a reasonable branch length estimation heuristic when branch lengths are not modeled or inferred under model misspecification, as in the case of Relate on array data. For ARG-Needle and ASMC-clust, ARG normalization improves the overall calibration of TMRCAs by making the range of predicted coalescence times closer to that expected from the demographic prior (see Supplementary Fig. S2a-d). Interestingly, however, ARG normalization decreases the TMRCA RMSE performance of ARG-Needle and ASMC-clust in both array and sequencing data (compare Supplementary Fig. S2d, e), such that ARG-Needle and ASMC-clust without ARG normalization consistently achieve the best TMRCA RMSE across methods (Supplementary Fig. S2e). This is likely linked to our use of ASMC's posterior mean TMRCA estimator, which is biased towards the average prior TMRCA but leads to good RMSE performance due to the weight placed by an L2 norm on outliers. ARG normalization is also likely to reduce the accuracy of the inferred height for root nodes in ARGs built using ARG-Needle and ASMC-clust, which have a large impact on TMRCA RMSE as a large fraction of pairwise coalescence events involve the root. It may be worthwhile to develop additional metrics that incorporate pairwise TMRCA values without relying so heavily on the root event or on large TMRCA values, for instance using an L1 norm instead of an L2 norm, or by measuring the RMSE of log TMRCA or of the square root of TMRCA. Note, however, that accuracy on the pairwise TMRCA RMSE is connected to several analyses discussed in this work in the context of ARG-GRMs. In particular increased performance for pairwise TMRCA RMSE reflects increased similarity (under the Frobenius norm) between true and inferred ARG-GRMs (when we assume $\alpha = 0$, see Methods), which we have shown may be utilized for heritability estimation, polygenic prediction, and mixed-model association.

##### KC distance

The Kendall-Colijn topology-only distance [26] (henceforth "KC distance" for short) has been used to evaluate inferred ARGs (e.g. in [15]). As a topology-only metric, it does not consider branch length or coalescence time information. Instead, it compares counts of the number of nodes that occur on the path from a desired most recent common ancestor node to the root node. The KC topology-only distance is affected whenever two virtually coincident coalescence events are joined to form a polytomy, or when polytomies are broken to form strictly bifurcating trees, since these operations create or remove nodes and thus alter the distances from internal nodes to the root. We provide further details on analyses we performed that show that the KC distance is systematically lower for methods that generate polytomies, as well as interpretations of this property in the context of ARG inference.

We first discuss the tree-wise KC distance and describe how we compute the KC distance across ARGs. Given two marginal trees $T_1$ and $T_2$, each with the same set of samples labeled 1 to $N$, the KC distance computation first calculates a vector of length $N(N-1)/2$ for each tree. For one of the trees $T$, let root($T$) denote the root node of the tree, and for any two nodes $a$ and $b$ in the tree, let $\text{mrca}_T(a, b)$ denote the most recent common ancestor of $a$ and $b$ and let $\text{dist}_T(a, b)$ denote the tree-path distance between nodes $a$ and $b$ in the tree, defined as the number of edges in the tree that need to be traversed to travel between $a$ and $b$. For any two distinct samples $1 \leq i < j \leq N$, let $\text{leaf}_T(i)$ and $\text{leaf}_T(j)$ denote the corresponding leaf nodes. We consider the MRCA of the two leaf nodes and count the distance from the root of the tree, computing

$$m_T(i,j) = \text{dist}_T\left(\text{root}(T), \text{mrca}_T(\text{leaf}_T(i), \text{leaf}_T(j))\right).$$

We then form vectors

$$\mathbf{m}_{T_1} = \left(m_{T_1}(1,2), m_{T_1}(1,3), \ldots, m_{T_1}(N-1,N)\right),$$
$$\mathbf{m}_{T_2} = \left(m_{T_2}(1,2), m_{T_2}(1,3), \ldots, m_{T_2}(N-1,N)\right).$$

(Note that in [26] the $\mathbf{m}_T$ vectors are augmented with additional entries, which are however irrelevant in our case.) Finally, we define the KC distance as the Euclidean distance between these two length $N(N-1)/2$ vectors:

$$\text{KC}(T_1, T_2) = ||\mathbf{m}_{T_1} - \mathbf{m}_{T_2}||_2 = \left(\sum_{1 \leq i < j \leq N} \left(m_{T_1}(i,j) - m_{T_2}(i,j)\right)^2\right)^{1/2}.$$

One option to combine tree-wise KC distances to get a comparison between ARGs is to weigh the KC metric by the genomic distance spanned by each tree, as done in [15]. For efficiency, this can be approximated by taking an unweighted average of the KC metric for several stabbing queries. In this work we opted to instead average the squared KC distance over the stabbing queries, then perform the square root (similar to our TMRCA RMSE calculations). A benefit of this approach is that it preserves the interpretation of the (ARG) KC distance as a Euclidean norm. (Suppose $\mathbf{m}_{1x}$ and $\mathbf{m}_{1y}$ are vectors from the first ARG at two locations $x$ and $y$, and $\mathbf{m}_{2x}$ and $\mathbf{m}_{2y}$ are vectors from the second ARG at two locations. Assume both locations receive a weight of $1/2$. Then our metric is equivalent to computing $||((\mathbf{m}_{1x}, \mathbf{m}_{1y}) - (\mathbf{m}_{2x}, \mathbf{m}_{2y}))/2||_2$, whereas the other approach results in $(||\mathbf{m}_{1x} - \mathbf{m}_{2x}||_2 + ||\mathbf{m}_{1y} - \mathbf{m}_{2y})||_2)/2$.)

To randomly resolve polytomies in marginal trees produced by tsinfer we replicated the approach used in [15]. At each polytomy with $k$ child edges coalescing, a random binary tree with $k$ leaves was generated and was substituted in place of the polytomy. In Supplementary Fig. S1b, we show the result of tsinfer with polytomies broken, where we sample the breaking procedure 10 times and average the KC results. As in [15], we observed that randomly resolving polytomies in tsinfer leads to reduced performance measured using KC distance.

We performed additional experiments where instead of breaking polytomies into bifurcations, we collapsed branches in a marginal tree to create polytomies. Because our goal is to measure similarity to the true ARG, we applied this operation only on inferred ARGs and not the true ARG. The merging operation takes a real parameter $f$ between 0 and 1 corresponding to the fraction of branches that are collapsed in each marginal tree of the inferred ARG. We implemented two types of merging: random merging and heuristic merging. In random merging, for each branch we sample a uniform random real number between 0 and 1, and if it is less than $f$ we collapse that branch. In heuristic merging, we aim to instead first collapse branches that are predicted with least confidence. We order the branches in the tree by computing the ratio of the branch length divided by the height of the parent node (to take into account that coalescent events in recent time tend to have short branches). After ordering, we select the fraction $f$ of the least certain branches and merge these to form polytomies. For either method of merging, we use a single merged tree per site and evaluate the KC distance against the marginal tree in the true ARG.

We tested values $f \in \{0, 0.05, 0.1, 0.2, 0.3, 0.4, 0.5, 0.75\}$, and measured the KC distance after applying merging for ARGs inferred by tsinfer, Relate, ASMC-clust, and ARG-Needle (Supplementary Fig. S1c-d). We used 4,000 sequences and 1 Mb for inference from sequences, and we used 4,000 array samples and 5 Mb for inference from array data. Relate, ASMC-clust, and ARG-Needle achieved lower KC distance when nodes were merged to form polytomies, including when the random merging strategy was used, suggesting that the KC distance is systematically lower for inferred trees that contain polytomies. Using heuristic merging in array data led to improvements for all methods, with Relate, ASMC-clust, and ARG-Needle performing better than tsinfer at the optimal merging fraction. In sequencing data, the KC performance of tsinfer was not improved by heuristic merging, and was matched by Relate, ASMC-clust, and ARG-Needle as $f$ was varied.

We briefly speculate on why collapsing branches to form polytomies may improve performance on the KC distance. The KC distance compares trees by computing the Euclidean norm between the two vectors $\mathbf{m}_{T_1}$ and $\mathbf{m}_{T_2}$, where we can take $T_1$ to be the true tree and $T_2$ to be the inferred tree, which will penalize large values. In our coalescent simulations, $T_1$ is a relatively well-balanced binary tree, so the values of $\mathbf{m}_{T_1}$ will range from $0$ (a pair of samples that coalesces at the root) to $O(\log N)$, with a mode and mean of $O(1)$. The introduction of polytomies may result in shrinkage of the values of $\mathbf{m}_{T_2}$, because the paths from MRCA nodes to the root contain fewer intermediate nodes, reducing variance but introducing bias (similar to what is observed for TMRCA RMSE). Additional analysis may help further understand the connections between the KC distance and properties of inferred ARGs that perform well under this metric, including their performance in downstream analyses.

#### Supplementary Note 3: Additional details on ARG-GRMs

In Methods we introduced ARG-GRMs for the case of haploid samples and $\alpha = 0$, as well as Monte Carlo ARG-GRMs, which sample new mutations on the ARG and use those markers to construct GRMs. Monte Carlo ARG-GRMs enable easily taking into account diploid samples, modeling varying values of $\alpha$, and working with stratified ARG-GRMs. In this Note, we describe computing an exact ARG-GRM for the cases of diploid samples, general $\alpha$, and stratification. We also discuss various invariances of the GRM used in mixed-model analysis and show how all methods compute an expected version of the sequence-based GRM up to invariance.

We first provide notation for the various sequence-based GRMs which we seek to approximate using ARGs. In all cases we have $M$ markers and $N$ individuals. While we have focused on haploid individuals with genotypes $x_{ik} \in \{0, 1\}$, $1 \leq i \leq N$ and $1 \leq i \leq M$, we also describe the case of diploid individuals with genotypes $x_{ik} \in \{0, 1, 2\}$, $1 \leq i \leq N$ and $1 \leq i \leq M$.

We begin by discussing the sequence-based GRM for haploid genotypes and general $\alpha$, with allele frequencies $p_k = \frac{1}{N} \sum_{i=1}^{N} x_{ik}$:

$$K_{\alpha,hap}(i, j) = \frac{1}{M} \sum_{k=1}^{M} \frac{(x_{ik} - p_k)(x_{jk} - p_k)}{[p_k(1 - p_k)]^{-\alpha}}. \tag{10}$$

We then describe stratified ARG-GRMs, which are obtained by partitioning the markers into various bins via MAF, LD, time intervals (to capture allele age), or other annotations. The haploid stratified ARG-GRM for a bin containing SNPs $B \subseteq \{1, \ldots, M\}$ using value $\alpha$ is given by:

$$K_{\alpha,hap,B}(i, j) = \frac{1}{|B|} \sum_{k=1}^{M} \mathbb{1}_{k \in B} \frac{(x_{ik} - p_k)(x_{jk} - p_k)}{[p_k(1 - p_k)]^{-\alpha}}. \tag{11}$$

(We use $\mathbb{1}_A$ to represent the indicator function of event $A$, taking value 1 if $A$ holds and 0 otherwise.)

Lastly, we consider the general $\alpha$ sequence-based GRM for diploid genotypes, with allele frequencies $p_k = \frac{1}{2N} \sum_{i=1}^{N} x_{ik}$:

$$K_{\alpha,dip}(i, j) = \frac{1}{M} \sum_{k=1}^{M} \frac{(x_{ik} - 2p_k)(x_{jk} - 2p_k)}{[2p_k(1 - p_k)]^{-\alpha}}. \tag{12}$$

(We omit a discussion on diploid stratified ARG-GRMs, which are easily derived as a generalization of the above two cases.)

##### From sequence-based GRMs to ARG-GRMs

The above GRM expressions assume we have access to a set of markers. Under the infinite sites assumption, each marker corresponds to an event that occurred at some time in the past. For simplicity, we refer to these events as mutations, though they may also consist of other variant types, such as short indels. Each such mutation occurs somewhere on the ARG, with position $x$ and time $t$, and affects a set of descendants that will carry the derived allele. In the haploid case, if we let the descendants of a mutation $m$ be $d(m) \subset \{1, \ldots, N\}$, we can also define the allele frequency $p(m)$ of a mutation as $p(m) = |d(m)|/N$.

Assuming a uniform mutation rate and the infinite-sites model, mutations arise uniformly over the area of the ARG, as a Poisson process characterized by the mutation rate $\mu$. The area of the ARG consists of all its edges, but an ARG edge does not always have a constant set of descendants, if a recombination event occurs on the path between the edge and its descendants. Therefore, we may further partition the edges of the ARG into "branches", such that each branch $b$ is valid for only part of the genomic extent of an edge and carries a constant set of descendants. We let $A(b)$ denote the area of branch $b$, obtained by multiplying its extent in time (e.g. generations) by its extent along the genome, and assume that the set $B$ enumerates all branches. Each branch $b$ then also inherits a set of descendants $d(b) \subset \{1, \ldots, N\}$ and an allele frequency $p(b)$, just as for mutations. The area of the ARG is then the sum over all these branches, or $\sum_{b \in B} A(b)$.

In the case where we do not have access to all underlying markers needed to compute a GRM but have access to the ground truth ARG or an inferred ARG, we may compute the expectation for the GRM entries. For the haploid single-component ARG-GRM, we compute

$$K_{\alpha,hap}(i,j) = \mathbb{E}_{m \sim Uniform(\{m_1,\ldots,m_M\})} \frac{\left(\mathbb{1}_{i \in d(m)} - p(m)\right)\left(\mathbb{1}_{j \in d(m)} - p(m)\right)}{\left[p(m)\left(1-p(m)\right)\right]^{-\alpha}}. \tag{13}$$

Since mutations are uniformly distributed over the area of an ARG, we replace the set of known $M$ mutations with the distribution over all possible mutations induced by uniform sampling over the ARG:

$$K_{\alpha,hap,ARG}(i,j) = \mathbb{E}_{m \sim Uniform(ARG)} \frac{\left(\mathbb{1}_{i \in d(m)} - p(m)\right)\left(\mathbb{1}_{j \in d(m)} - p(m)\right)}{\left[p(m)\left(1-p(m)\right)\right]^{-\alpha}}. \tag{14}$$

(14) reflects the expected value of (13) given the genealogical relationships of the ARG, without observing any markers.

$$K_{\alpha,hap,ARG}(i,j) = \mathbb{E}\left[K_{\alpha,hap}(i,j)|ARG\right]. \tag{15}$$

We can compute the ARG-wide expectation given by (14) using Monte Carlo, adopting a mutation rate $\mu$ to uniformly sample $M'$ variants on the ARG. We then use these $M'$ sampled variants to evaluate (14).

We can also take this expectation analytically, by weighting all possible branches of the ARG by their area. Using our earlier notation,

$$\begin{aligned} K_{\alpha,hap,ARG}(i,j) &= \mathbb{E}_{m \sim Uniform(ARG)} \frac{\left(\mathbb{1}_{i \in d(m)} - p(m)\right)\left(\mathbb{1}_{j \in d(m)} - p(m)\right)}{\left[p(m)\left(1-p(m)\right)\right]^{-\alpha}} \\ &= \frac{1}{\sum_{b \in B} A(b)} \sum_{b \in B} \left[A(b) \cdot \frac{\left(\mathbb{1}_{i \in d(b)} - p(b)\right)\left(\mathbb{1}_{j \in d(b)} - p(b)\right)}{\left[p(b)\left(1-p(b)\right)\right]^{-\alpha}}\right] \end{aligned} \tag{16}$$

This formulation, however, requires traversing all branches of the ARG and updating $N^2$ values for each branch, which is less efficient than using the Monte Carlo method with a sufficiently high mutation rate. The Monte Carlo method is also $O(N^2)$ per mutation, but fewer mutations are needed to be sampled compared to the exact ARG-GRM, which visits all branches.

The case of stratified GRMs is an extension of the above, where we assign each mutation to different GRMs according to the stratification criteria. In our experiments, this corresponded to selecting the appropriate GRM based on the allele frequency ($p(m)$ or $p(b)$) of each sampled mutation.

For diploid GRMs, each individual genotype $x_{ik}$ is modeled as the sum of two haplotypes. We introduce notation where we consider an ARG of $2N$ haploid samples, numbered 1 to $2N$. Without loss of generality, each individual $i$ consists of haplotypes $2i-1$ and $2i$, for $1 \leq i \leq N$. For shorthand, we define $i_1 = 2i-1$ and $i_2 = 2i$. Then we can rewrite (12) as

$$K_{\alpha,dip}(i,j) = \frac{1}{M}\sum_{k=1}^{M} \frac{(x_{i_1k} + x_{i_2k} - 2p_k)(x_{j_1k} + x_{j_2k} - 2p_k)}{[2p_k(1-p_k)]^{-\alpha}}. \tag{17}$$

If we denote descendants of a mutation in the ARG as $d(m) \subset \{1,\dots,2N\}$, and $p(m) = |d(m)|/2N$, the diploid ARG-GRM can be computed using the ARG as

$$\begin{aligned} &K_{\alpha,dip,ARG}(i,j) \\ &= \mathbb{E}_{m\sim Uniform(ARG)} \frac{\left(\mathbb{1}_{i_1\in d(m)} + \mathbb{1}_{i_2\in d(m)} - 2p(m)\right)\left(\mathbb{1}_{j_1\in d(m)} + \mathbb{1}_{j_1\in d(m)} - 2p(m)\right)}{[2p(m)(1-p(m))]^{-\alpha}} \\ &= \frac{1}{\sum_{b\in B} A(b)} \sum_{b\in B}\left[A(b)\cdot \frac{\left(\mathbb{1}_{i_1\in d(b)} + \mathbb{1}_{i_2\in d(b)} - 2p(b)\right)\left(\mathbb{1}_{j_1\in d(b)} + \mathbb{1}_{j_2\in d(b)} - 2p(b)\right)}{[2p(b)(1-p(b))]^{-\alpha}}\right] \end{aligned}$$

The exact ARG-GRM is obtained by iterating over all branches, and a Monte Carlo ARG-GRM is obtained by sampling, as previously described.

In the remainder we derive simpler expressions for ARG-GRMs. We begin by describing three useful invariances of GRMs in mixed-model analysis.

##### Three GRM invariances

We discuss three invariances under which a GRM may be altered without changing downstream results, which we use to facilitate mixed model analysis using ARG-GRMs and in some of the proofs below. We refer to these invariances as scale invariance, data shift invariance, and constant shift invariance.

The two aspects of the ARG-LMM pipeline that are linked to these invariances are Gower centering and the inclusion of a centering covariate. We first describe Gower centering. Given an $N$ by $N$ GRM $K$, define the $N$ by $N$ identity matrix $I_N$, the size $N$ column vector $1_N$, and the projection matrix $P_N$ (which is symmetric and idempotent and corresponds to projection onto the subspace orthogonal to $1_N$) to be

$$P_N = I_N - \frac{1}{N} 1_N 1_N^T.$$

Then, the Gower centered version of $K$ is defined as

$$C_{Gower}(K) = \frac{N-1}{\text{Tr}(P_N K P_N)} K.$$

To obtain correct heritability estimates for sequence-based GRMs with general $\alpha$ (e.g. (10) above), we Gower centered ARG-GRMs provided in input to GCTA. (In the special case of $\alpha = 0$, the sequence-based GRM in (10) is approximately Gower centered already, and Gower centering merely multiplies by the factor $(N-1)/N$.) Consider the multiplication of $K$ by any nonzero scalar constant $\gamma$:

$$C_{Gower}(\gamma K) = \frac{N-1}{\text{Tr}(P_N(\gamma K)P_N)}\gamma K = \frac{N-1}{\gamma\text{Tr}(P_N K P_N)}\gamma K = C_{Gower}(K).$$

Thus, multiplying a GRM by any nonzero scalar and then applying Gower centering will lead to identical downstream results, which we refer to as *scale invariance* of GRMs.

We next consider the inclusion of a centering covariate. Even when no covariates are specified in an analysis, most software packages for complex trait analysis (e.g. PLINK [27], GCTA [28], BOLT-LMM [29, 30], and BOLT-REML [31]) implicitly or explicitly include a centering covariate. This can be thought of as a length $N$ covariate vector consisting of all 1s. Including this covariate is equivalent to mean-centering the phenotype as well as the genotype vector for each marker. It can also be implemented by applying the projection operator $P_N$, defined above, to project out the component parallel to $1_N$ in the data.

In our analyses involving GCTA, we provided the GRM and a phenotype in input, rather than providing markers from which to compute the GRM. In this case, the phenotype is mean-centered, and although one does not have access to the underlying genotypes, the centering covariate is still applied implicitly during the relevant mixed-model calculations.

We also describe the inclusion of a centering covariate as a transformation on the GRM itself. In our analyses we performed this operation, which we call data centering, prior to Gower centering, and after both steps were completed, we provided the transformed GRM in input to GCTA. The data centered GRM is given by

$$C_{Data}(K) = P_N K P_N.$$

Note that right-multiplying by $P_N$ corresponds to subtracting the mean column of a matrix from each column and left-multiplying by $P_N$ corresponds to subtracting the mean row of a matrix from each row. Also note that the transformations $C_{Gower}$ and $C_{Data}$ commute, so that the order in which they are applied does not matter, and that they are both idempotent, so that data centering outside of GCTA does not interfere with the application of the centering covariate inside.

We highlight the second invariance of GRMs we leveraged in these analyses using an example of the effects of data centering. Consider the GRM that would arise if we used raw genotypes instead of centered genotypes in (10):

$$\widetilde{K}_{\alpha,hap}(i,j) = \frac{1}{M}\sum_{k=1}^{M}\frac{(x_{ik})(x_{jk})}{[p_k(1-p_k)]^{-\alpha}}.$$

We verify that data centering gives the expression (10) with centered genotypes. Notice that the GRM $\widetilde{K}_{\alpha,hap}$ can be written as a product of matrices:

$$\widetilde{K}_{\alpha,hap} = XDX^T$$

where $X$ is of size $N$ by $M$ with entries $x_{ij}$, and $D$ is a diagonal matrix of size $M$ by $M$ with diagonal entries

$$d_{kk} = \left[p_k\left(1-p_k\right)\right]^{\alpha}/M.$$

The data centered version of $\widetilde{K}_{\alpha,hap}$ is then

$$C_{Data}\left(\widetilde{K}_{\alpha,hap}\right) = P_N\widetilde{K}_{\alpha,hap}P_N = P_N(XDX^T)P_N = (P_NX)D(P_NX)^T.$$

Also notice that

$$P_NX = \left(I_N - \frac{1}{N}1_N1_N^T\right)X = X - 1_N\left(\frac{1}{N}1_N^TX\right) = X - 1_N\mu^T$$

where $\mu^T = (p_1, p_2, \ldots, p_M)$ is a row vector consisting of the average row of $X$, or the collection of allele frequencies. (So that left-multiplying by $P_N$ corresponds to subtracting the mean row of a matrix from each row.) Hence,

$$C_{Data}\left(\widetilde{K}_{\alpha,hap}\right) = (X - 1_N\mu^T)D(X - 1_N\mu^T)^T.$$

This is equivalent to forming a GRM using the centered data, and hence $C_{Data}(\widetilde{K}_{\alpha,hap}) = K_{\alpha,hap}$.

The invariance highlighted in this example, where centered data is replaced by raw genotypes, is more general. Rather than using raw genotypes, we can add a marker-specific constant to the genotypes for each marker before multiplying:

$$\widehat{K}_{\alpha,hap}(i,j) = \frac{1}{M}\sum_{k=1}^{M}\frac{\left(x_{ik}+c_k\right)\left(x_{jk}+c_k\right)}{\left[p_k\left(1-p_k\right)\right]^{-\alpha}},$$

where the $c_k$ are any real scalars for $1 \leq k \leq M$. If we let $\rho^T = (c_1, \ldots, c_M)$, we see that

$$\widehat{K}_{\alpha,hap} = (X + 1_N\rho^T)D(X + 1_N\rho^T)^T.$$

We have

$$\begin{aligned}
P_N(X + 1_N\rho^T) &= P_NX + P_N1_N\rho^T \\
&= (X - 1_N\mu^T) + \left(I_N - \frac{1}{N}1_N1_N^T\right)1_N\rho^T \\
&= (X - 1_N\mu^T) + \left(1_N - \frac{1}{N}1_NN\right)\rho^T \\
&= X - 1_N\mu^T.
\end{aligned}$$

Hence

$$C_{Data}\left(\widehat{K}_{\alpha,hap}\right) = P_N\widehat{K}_{\alpha,hap}P_N = (X - 1_N\mu^T)D(X - 1_N\mu^T)^T = K_{\alpha,hap}.$$

We refer to this invariance as *data shift invariance*: by using a centering covariate, the markers used to construct a GRM can have a constant shift per marker applied to the genotypes before multiplying, without affecting downstream results. Although we have only described a detailed derivation in the case of haploid GRMs, the same considerations apply to other cases, starting from (11) and (12). Note that this form of invariance was also discussed in [32], Appendix B.

Finally, *constant shift invariance* allows adding or subtracting a constant scalar to each entry of a GRM without affecting downstream results. This invariance also follows from the inclusion of a centering covariate. Assume data centering and consider adding a constant $c$ to each entry of a GRM $K$:

$$\begin{aligned} C_{Data}(K + c1_N 1_N^T) &= P_N(K + c1_N 1_N^T)P_N \\ &= C_{Data}(K) + \left(I_N - \frac{1}{N}1_N 1_N^T\right)(c1_N 1_N^T)\left(I_N - \frac{1}{N}1_N 1_N^T\right) \\ &= C_{Data}(K) + c\left(1_N 1_N^T - 1_N 1_N^T - 1_N 1_N^T + 1_N 1_N^T\right) \\ &= C_{Data}(K). \end{aligned}$$

In our experiments, we applied data centering directly on the GRM before passing into GCTA, which guaranteed constant shift invariance. We observed that if we omitted the data centering transformation, only relying on GCTA's implementation of a centering covariate, constant shift invariance still held for a range of shift values. However, some experiments involved adding a large negative shift to a GRM such that it is no longer positive semidefinite, and we observed that passing such a GRM directly into GCTA led to errors. On the other hand, we found that GCTA was robust to the earlier described data shift invariance, including when we omitted the data centering transformation, possibly because data shift transformations preserve positive definiteness.

In summary, by always applying Gower centering and what we referred to as data centering to our GRMs, before passing into GCTA, allowed us to guarantee three invariances in the GRM: scale invariance, where the GRM is multiplied by a non-zero scalar; data shift invariance, where a marker-specific shift is applied to each marker used to compute the GRM; and constant shift invariance, where a constant scalar is added to each entry of the GRM.

##### Derivation of exact ARG-GRM, haploid and general $\alpha$

We use these invariances to describe simplified expressions for computing exact ARG-GRMs. Due to data shift invariance, we may replace the $p_k$ terms in the numerator of (10) with the value $1/2$. Let $\equiv$ denote equivalence under invariance. We can write

$$K_{\alpha,hap}(i,j) \equiv \frac{1}{M}\sum_{k=1}^{M}\frac{(x_{ik}-1/2)(x_{jk}-1/2)}{[p_k(1-p_k)]^{-\alpha}}. \tag{18}$$

(Note that with this notation we are referring to equivalence under invariance for the entire GRM, not for an individual entry.) Since $x_{ik}, x_{jk} \in \{0, 1\}$,

$$\begin{aligned}(x_{ik} - 1/2)(x_{jk} - 1/2) &= \frac{1}{4}(2x_{ik} - 1)(2x_{jk} - 1) \\ &= \frac{1}{4}(1 - 2(x_{ik} \oplus x_{jk})),\end{aligned}$$

where $\oplus$ refers to the XOR of two binary values. Substituting into (18) and leveraging scale invariance and constant shift invariance,

$$\begin{aligned}K_{\alpha,hap}(i,j) &\equiv \frac{1}{4M}\sum_{k=1}^{M}\frac{(1 - 2(x_{ik} \oplus x_{jk}))}{[p_k(1-p_k)]^{-\alpha}} \\ &= \frac{1}{4M}\sum_{k=1}^{M}\frac{1}{[p_k(1-p_k)]^{-\alpha}} - \frac{1}{2M}\sum_{k=1}^{M}\frac{x_{ik} \oplus x_{jk}}{[p_k(1-p_k)]^{-\alpha}} \\ &\equiv \sum_{k=1}^{M}\frac{x_{ik} \oplus x_{jk}}{[p_k(1-p_k)]^{-\alpha}}. \end{aligned} \tag{19}$$

In the case of $\alpha = 0$, this simplifies to

$$K_{\alpha=0,hap}(i,j) \equiv \sum_{k=1}^{M} x_{ik} \oplus x_{jk}.$$

Note that this is the Hamming distance matrix between $i$ and $j$. Under the infinite-sites model, mutations for which samples $i$ and $j$ differ occur on the area of the 2-sample ARG from samples $i$ and $j$ to their TMRCA along the genome. The number of such mutations is a Poisson random variable with rate $\mu$ (the per-site per-generation mutation rate) times the area (sites times generations) of the 2-sample ARG. We express the areas as $2 \times L \times \bar{t}_{ij}$, where $\bar{t}_{ij}$ is the genome-wide average TMRCA of $i$ and $j$ and $L$ is the physical extent of the ARG. Then,

$$\begin{aligned}K_{\alpha=0,hap,ARG}(i,j) &= \mathbb{E}\left[K_{\alpha=0,hap}(i,j)|ARG\right] \\ &\equiv \mathbb{E}\left[\sum_{k=1}^{M} x_{ik} \oplus x_{jk}\middle| ARG\right] \\ &= \mathbb{E}\left[\text{Poisson}(2 \times L \times \bar{t}_{ij})\right] \\ &= 2 \times L \times \bar{t}_{ij} \\ &\equiv \bar{t}_{ij}.\end{aligned}$$

Assuming Gower centering and data centering are used, we may therefore use the average TMRCA matrix between samples to compute the $\alpha = 0$ exact ARG-GRM. (Note that in the case of $\alpha = 0$ allele frequencies are not involved, so that the ARG-GRM may be estimated using pairwise TMRCA estimates alone [33], e.g. using ASMC.)

For the case of general $\alpha$, consider (19), then

$$\begin{aligned}K_{\alpha,hap,ARG}(i,j) &\equiv \mathbb{E}\left[\frac{1}{M}\sum_{k=1}^{M}\frac{x_{ik}\oplus x_{jk}}{\left[p_k\left(1-p_k\right)\right]^{-\alpha}}\middle| ARG\right] \\ &= \mathbb{E}_{m\sim Uniform(ARG)}\left[\frac{\mathbb{1}_{i\in d(m)}\oplus \mathbb{1}_{j\in d(m)}}{\left[p(m)\left(1-p(m)\right)\right]^{-\alpha}}\right] \\ &= \sum_{b\in B}\left[A(b)\cdot\frac{\mathbb{1}_{i\in d(b)}\oplus \mathbb{1}_{j\in d(b)}}{\left[p(b)\left(1-p(b)\right)\right]^{-\alpha}}\right]\end{aligned}$$

The expression $\mathbb{1}_{i\in d(b)}\oplus \mathbb{1}_{j\in d(b)}$ is 1 if exactly one of $i$ and $j$ is a descendant of branch $b$, and 0 otherwise. Therefore we may rewrite it as $\mathbb{1}_{|d(b)\cap\{i,j\}|=1}$, where we test whether the intersection of $d(b)$ and $\{i,j\}$ contains exactly one element. Thus:

$$\begin{aligned}K_{\alpha,hap,ARG}(i,j) &\equiv \sum_{b\in B}\left[A(b)\cdot\frac{\mathbb{1}_{|d(b)\cap\{i,j\}|=1}}{\left[p(b)\left(1-p(b)\right)\right]^{-\alpha}}\right] \\ &= \sum_{b\in B,|d(b)\cap\{i,j\}|=1}\left[\frac{A(b)}{\left[p(b)\left(1-p(b)\right)\right]^{-\alpha}}\right] \qquad (20)\end{aligned}$$

The branches $b\in B$ with $|d(b)\cap\{i,j\}|=1$ are those that lie in the 2-sample ARG containing samples $i$ and $j$. Computing (20) is achieved by iterating over all these branches and summing their areas, weighted by a term involving $p(b)$ and $\alpha$. When $\alpha=0$, this reduces to summing the areas of all branches in the 2-sample ARG:

$$\begin{aligned}K_{\alpha=0,hap,ARG}(i,j) &\equiv \sum_{b\in B,|d(b)\cap\{i,j\}|=1} A(b) \\ &= 2\times L\times \bar{t}_{ij},\end{aligned}$$

which coincides with the earlier derivation for $\alpha=0$.

The MAF-stratified version of the haploid $\alpha$ GRM is similarly derived, except iteration is restricted to branches $b$ with allele frequency $p(b)$ within a specified range. This generalizes to other stratification criteria.

#### Derivation of exact ARG-GRM, diploid and general $\alpha$

Finally, we describe the case of diploid genotypes and general $\alpha$. We separate (17) into four terms:

$$
\begin{aligned}
K_{\alpha,dip}(i,j) &= \frac{1}{M}\sum_{k=1}^{M}\frac{(x_{i_1k}+x_{i_2k}-2p_k)(x_{j_1k}+x_{j_2k}-2p_k)}{[2p_k(1-p_k)]^{-\alpha}} \\
&= \frac{1}{M}\sum_{k=1}^{M}\frac{[(x_{i_1k}-p_k)+(x_{i_2k}-p_k)][(x_{j_1k}-p_k)+(x_{j_2k}-p_k)]}{[2p_k(1-p_k)]^{-\alpha}} \\
&= \frac{2^\alpha}{M}\left[\sum_{k=1}^{M}\frac{(x_{i_1k}-p_k)(x_{j_1k}-p_k)}{[p_k(1-p_k)]^{-\alpha}}+\sum_{k=1}^{M}\frac{(x_{i_1k}-p_k)(x_{j_2k}-p_k)}{[p_k(1-p_k)]^{-\alpha}}+\right. \\
&\qquad\left.\sum_{k=1}^{M}\frac{(x_{i_2k}-p_k)(x_{j_1k}-p_k)}{[p_k(1-p_k)]^{-\alpha}}+\sum_{k=1}^{M}\frac{(x_{i_2k}-p_k)(x_{j_2k}-p_k)}{[p_k(1-p_k)]^{-\alpha}}\right] \\
&= 2^\alpha\left[K_{\alpha,hap}(i_1,j_1)+K_{\alpha,hap}(i_1,j_2)+K_{\alpha,hap}(i_2,j_1)+K_{\alpha,hap}(i_2,j_2)\right],
\end{aligned}
$$

where terms such as $K_{\alpha,hap}(i_1,j_1)$ correspond to the haploid GRM over $2N$ samples given by (10). Using scale invariance, the factor of $2^\alpha$ can be removed to write

$$K_{\alpha,dip}(i,j) \equiv K_{\alpha,hap}(i_1,j_1)+K_{\alpha,hap}(i_1,j_2)+K_{\alpha,hap}(i_2,j_1)+K_{\alpha,hap}(i_2,j_2).$$

A similar expression may be obtained for the ARG-GRMs:

$$
\begin{aligned}
K_{\alpha,dip,ARG}(i,j) &= \mathbb{E}\left[K_{\alpha,dip}(i,j)|ARG\right] \\
&\equiv \mathbb{E}\left[K_{\alpha,hap}(i_1,j_1)|ARG\right]+\mathbb{E}\left[K_{\alpha,hap}(i_1,j_2)|ARG\right]+ \\
&\qquad \mathbb{E}\left[K_{\alpha,hap}(i_2,j_1)|ARG\right]+\mathbb{E}\left[K_{\alpha,hap}(i_2,j_2)|ARG\right] \\
&= K_{\alpha,hap,ARG}(i_1,j_1)+K_{\alpha,hap,ARG}(i_1,j_2)+K_{\alpha,hap,ARG}(i_2,j_1)+K_{\alpha,hap,ARG}(i_2,j_2).
\end{aligned}
$$

The diploid ARG-GRM is therefore obtained by first computing the exact ARG-GRM of size $2N$ by $2N$ over the haploid samples and then using terms involving $(i_1,j_1)$, $(i_1,j_2)$, $(i_2,j_1)$, and $(i_2,j_2)$ to compute the entry for pair $(i,j)$.

#### References (Supplementary Information)

[1] Robert C Griffiths and Paul Marjoram. An ancestral recombination graph. *Institute for Mathematics and its Applications*, 87:257, 1997.

[2] John Frank Charles Kingman. The coalescent. *Stochastic Processes and their Applications*, 13(3):235–248, 1982.

[3] Richard R Hudson. Properties of a neutral allele model with intragenic recombination. *Theoretical Population Biology*, 23(2):183–201, 1983.

[4] Richard R Hudson et al. Gene genealogies and the coalescent process. *Oxford Surveys in Evolutionary Biology*, 7(1):44, 1990.

[5] Jotun Hein, Mikkel Schierup, and Carsten Wiuf. *Gene genealogies, variation and evolution: a primer in coalescent theory*. Oxford University Press, 2004.

[6] Matthew D Rasmussen, Melissa J Hubisz, Ilan Gronau, and Adam Siepel. Genome-wide inference of ancestral recombination graphs. *PLoS Genetics*, 10(5):e1004342, 2014.

[7] Carsten Wiuf and Jotun Hein. The ancestry of a sample of sequences subject to recombination. *Genetics*, 151(3):1217–1228, 1999.

[8] Carsten Wiuf and Jotun Hein. Recombination as a point process along sequences. *Theoretical Population Biology*, 55(3):248–259, 1999.

[9] Gilean AT McVean and Niall J Cardin. Approximating the coalescent with recombination. *Philosophical Transactions of the Royal Society B: Biological Sciences*, 360(1459):1387–1393, 2005.

[10] Paul Marjoram and Jeff D Wall. Fast "coalescent" simulation. *BMC Genetics*, 7(1):1–9, 2006.

[11] Gary K Chen, Paul Marjoram, and Jeffrey D Wall. Fast and flexible simulation of DNA sequence data. *Genome Research*, 19(1):136–142, 2009.

[12] Laurent Excoffier and Matthieu Foll. Fastsimcoal: a continuous-time coalescent simulator of genomic diversity under arbitrarily complex evolutionary scenarios. *Bioinformatics*, 27(9):1332–1334, 2011.

[13] Paul R Staab, Sha Zhu, Dirk Metzler, and Gerton Lunter. scrm: Efficiently simulating long sequences using the approximated coalescent with recombination. *Bioinformatics*, 31(10):1680–1682, 2015.

[14] Jerome Kelleher, Alison M Etheridge, and Gilean McVean. Efficient coalescent simulation and genealogical analysis for large sample sizes. *PLoS Computational Biology*, 12(5):e1004842, 2016.

[15] Jerome Kelleher, Yan Wong, Anthony W Wohns, Chaimaa Fadil, Patrick K Albers, and Gil McVean. Inferring whole-genome histories in large population datasets. *Nature Genetics*, 51(9):1330–1338, 2019.

[16] Leo Speidel, Marie Forest, Sinan Shi, and Simon R Myers. A method for genome-wide genealogy estimation for thousands of samples. *Nature Genetics*, 51(9):1321–1329, 2019.

[17] Pier Francesco Palamara. ARGON: fast, whole-genome simulation of the discrete time Wright-Fisher process. *Bioinformatics*, 32(19):3032–3034, 2016.

[18] Pier Francesco Palamara, Jonathan Terhorst, Yun S Song, and Alkes L Price. High-throughput inference of pairwise coalescence times identifies signals of selection and enriched disease heritability. *Nature Genetics*, 50(9):1311–1317, 2018.

[19] Richard Durbin. Efficient haplotype matching and storage using the positional Burrows–Wheeler transform (PBWT). *Bioinformatics*, 30(9):1266–1272, 2014.

[20] Brian L Browning, Ying Zhou, and Sharon R Browning. A one-penny imputed genome from next-generation reference panels. *The American Journal of Human Genetics*, 103(3):338–348, 2018.

[21] Olivier Delaneau, Jean-François Zagury, Matthew R Robinson, Jonathan L Marchini, and Emmanouil T Dermitzakis. Accurate, scalable and integrative haplotype estimation. *Nature Communications*, 10(1):1–10, 2019.

[22] Simone Rubinacci, Olivier Delaneau, and Jonathan Marchini. Genotype imputation using the Positional Burrows Wheeler Transform. *PLoS Genetics*, 16(11):e1009049, 2020.

[23] Po-Ru Loh, Petr Danecek, Pier Francesco Palamara, Christian Fuchsberger, Yakir A Reshef, Hilary K Finucane, Sebastian Schoenherr, Lukas Forer, Shane McCarthy, Goncalo R Abecasis, et al. Reference-based phasing using the Haplotype Reference Consortium panel. *Nature Genetics*, 48(11):1443–1448, 2016.

[24] Juba Nait Saada, Georgios Kalantzis, Derek Shyr, Fergus Cooper, Martin Robinson, Alexander Gusev, and Pier Francesco Palamara. Identity-by-descent detection across 487,409 British samples reveals fine scale population structure and ultra-rare variant associations. *Nature Communications*, 11(1):1–15, 2020.

[25] Sebastian Böcker and Andreas WM Dress. Recovering symbolically dated, rooted trees from symbolic ultrametrics. *Advances in Mathematics*, 138(1):105–125, 1998.

[26] Michelle Kendall and Caroline Colijn. Mapping phylogenetic trees to reveal distinct patterns of evolution. *Molecular Biology and Evolution*, 33(10):2735–2743, 2016.

[27] Shaun Purcell, Benjamin Neale, Kathe Todd-Brown, Lori Thomas, Manuel AR Ferreira, David Bender, Julian Maller, Pamela Sklar, Paul IW De Bakker, Mark J Daly, et al. PLINK: a tool set for whole-genome association and population-based linkage analyses. *The American Journal of Human Genetics*, 81(3):559–575, 2007.

[28] Jian Yang, S Hong Lee, Michael E Goddard, and Peter M Visscher. GCTA: a tool for genome-wide complex trait analysis. *The American Journal of Human Genetics*, 88(1):76–82, 2011.

[29] Po-Ru Loh, George Tucker, Brendan K Bulik-Sullivan, Bjarni J Vilhjalmsson, Hilary K Finucane, Rany M Salem, Daniel I Chasman, Paul M Ridker, Benjamin M Neale, Bonnie Berger, et al. Efficient Bayesian mixed-model analysis increases association power in large cohorts. *Nature Genetics*, 47(3):284, 2015.

[30] Po-Ru Loh, Gleb Kichaev, Steven Gazal, Armin P Schoech, and Alkes L Price. Mixed-model association for biobank-scale datasets. *Nature Genetics*, 50(7):906–908, 2018.

[31] Po-Ru Loh, Gaurav Bhatia, Alexander Gusev, Hilary K Finucane, Brendan K Bulik-Sullivan, Samuela J Pollack, Teresa R de Candia, Sang Hong Lee, Naomi R Wray, Kenneth S Kendler, et al. Contrasting genetic architectures of schizophrenia and other complex diseases using fast variance-components analysis. *Nature Genetics*, 47(12):1385, 2015.

[32] Andy Dahl, Khiem Nguyen, Na Cai, Michael J Gandal, Jonathan Flint, and Noah Zaitlen. A robust method uncovers significant context-specific heritability in diverse complex traits. *The American Journal of Human Genetics*, 106(1):71–91, 2020.

[33] Pier Francesco Palamara, Jonathan Terhorst, Yun Song, and Alkes Price. Leveraging deep genealogical structure to estimate the phenotypic contribution of rare variants. Presented at the 66th Annual Meeting of The American Society of Human Genetics, Vancouver, 2016. Abstract 1931T.
